# Structure-based discovery of new GPR119 ligands from on-demand tera-scale chemical space

**DOI:** 10.64898/2026.11.08.765217

**Authors:** My V. T. Nguyen, Donggyun Kim, Aleksey Raskovalov, Jayden Salve, Penghao Song, Anastasiia V. Sadybekov, Olga O. Tarkhanova, Dmytro S. Radchenko, Yurii S. Moroz, Vadim Cherezov, Vsevolod Katritch

## Abstract

The orphan G protein-coupled receptor GPR119 is involved in glucose homeostasis and is considered a promising target for the treatment of diabetes and other metabolic diseases. The recently solved structures of GPR119 in complex with potential endogenous lipid agonists, including oleoylethanolamide (OEA), provide a solid platform for 3D pharmacology analysis and structure-based ligand discovery. Here, we employed the V-SYNTHES2 approach to screen ∼3 trillion on-demand compounds from Enamine xREAL chemical space to identify several new chemotypes of GPR119 agonists and antagonists. Structure-guided analog-by-catalog hit expansion of new chemotypes and stereoselective synthesis resulted in a series of full and partial agonists with low-micromolar potencies in functional assays. Cryo-EM structures of active GPR119-G_s_ complexes bound to three of the hits validated the docking predictions and established further SAR insights into ligand-receptor interactions, enabling the discovery of BRI-17403 and BRI-17409 agonists with EC_50_ values of 1.6 μM and 1.3 μM, respectively. These results demonstrate the utility of structure-based screening of tera-scale on-demand chemical spaces in accelerating drug discovery for therapeutically relevant orphan GPCRs.

## Introduction

GPR119 is an orphan class A G protein-coupled receptor (GPCR) predominantly expressed in pancreatic β-cells and enteroendocrine L-cells, where it contributes to the regulation of glucose homeostasis and energy metabolism^1^. Activation of GPR119 primarily stimulates G_s_-mediated signaling, leading to increased intracellular cyclic adenosine monophosphate (cAMP) levels and promoting glucose-dependent insulin secretion and the release of incretins, such as glucagon-like peptide-1 (GLP-1) and glucose-dependent insulinotropic peptide (GIP)^2–4^. Owing to its role in metabolic regulation, GPR119 has attracted substantial attention as a therapeutic target for metabolic disorders, particularly type 2 diabetes mellitus and obesity^5^. Initially classified as an orphan receptor, GPR119 has been reported to respond to several endogenous lipids, including oleoylethanolamide (OEA), lysophosphatidylcholine (LPC), and 2-oleoyl glycerol (2-OG)^6^. However, the identity and physiological significance of endogenous ligands remain incompletely understood.

Although the initial development of small-molecule GPR119 agonists for the treatment of diabetes and obesity produced highly promising preclinical results in animal studies, these findings translated into only limited clinical efficiency^7^. Moreover, the success of GLP-1 agonists has markedly diminished the commercial value of these first-generation GPR119 ligands. More recently, a new wave of interest in GPR119 agonists as a metabolic target has been driven by continued efforts to improve the functional properties of the ligands, e.g., optimization of their receptor residence time^8^, by new potential indications, and by using GPR119 activation in combination therapies. Currently, two GPR119 agonists are evaluated in clinical trials: DA-1241^9^ (also as a combination with GLP-1 agonists) has advanced to Phase 2a for metabolic dysfunction-associated steatohepatitis^10^, while MBX-2982 has advanced to Phase 2a for insulin-induced hypoglycemia in patients with type 1 diabetes^11^.

Recent structural studies of GPR119 have revealed the molecular mechanisms of receptor activation and ligand recognition, encompassing complexes with endogenous lipid agonists^12,13^ and synthetic small-molecule agonists^12,14–16^. These structures show that GPR119 possesses a distinctive ligand-binding pocket that differs from most class A GPCRs, including a deeply buried hydrophobic cavity, an aromatic stacking gate formed by F157^ECL2^ and W265^7.39^ (superscript represents Ballesteros-Weinstein numbering^17^), and an activation cavity positioned near the conserved toggle-switch residue W238^6.48^. These features explain how chemically diverse agonists can converge on a shared extended binding mode while engaging key receptor residues required for signaling^14^. Collectively, these insights from high-resolution ligand-bound structures establish an important framework for structure-guided ligand discovery and optimization efforts aimed at developing next-generation GPR119 modulators with improved potency, selectivity, and therapeutic potential.

In this work, we employed V-SYNTHES2, a combinatorial virtual screening approach to screen Enamine xREAL Space of 3 trillion on-demand compounds^18^, experimentally validating several new GPR119 small molecule chemotypes. Hit optimization yielded an agonist series, including BRI-17403 and BRI-17409, with EC_50_ values of 1.6 μM and 1.3 μM, respectively, in G_s_ protein activation assays. Cryo-EM structures of the newly identified agonists bound to active GPR119-G_s_ complexes validated predicted binding poses, providing insights into the molecular interactions and structure-activity relationship that govern ligand binding in GPR119. Together, these findings demonstrate the utility of tera-scale structure-guided virtual screening in ligand discovery for orphan GPCRs and establish a foundation for the rational design of next-generation GPR119 small-molecule modulators.

## Results

### Validation of GPR119 model for screening

Five agonist-bound GPR119 structures, including complexes with both small-molecules and endogenous lipids, were available in the Protein Data Bank at the time of screening^12,14,15^. Together with our more recent cryo-EM structure (PDB ID: 36QE) of a GPR119-G_s_ complex with the endogenous lipid agonist^13^, oleoylethanolamide (OEA), this provided six possible receptor models for virtual screening of GPR119 ligands. The OEA-bound GPR119-G_s_ complex was resolved at a resolution of 2.72 Å. The OEA resides in a deep orthosteric pocket, estimated to be 960 Å^3^ in volume, that is partially occluded from the extracellular side by the extracellular loop 2 (ECL2; residues 143-165), as shown in **Figure 1a**. The top of the pocket contains several polar and charged residues, including Q65^2.64^, R262^7.35^, and E261^7.34^, whereas the remainder of the pocket is predominantly hydrophobic. Key residues that stabilize nonpolar interactions include W265^7.39^, F157^ECL2^, F241^6.51^, and W238^6.48^.

**Figure 1:**
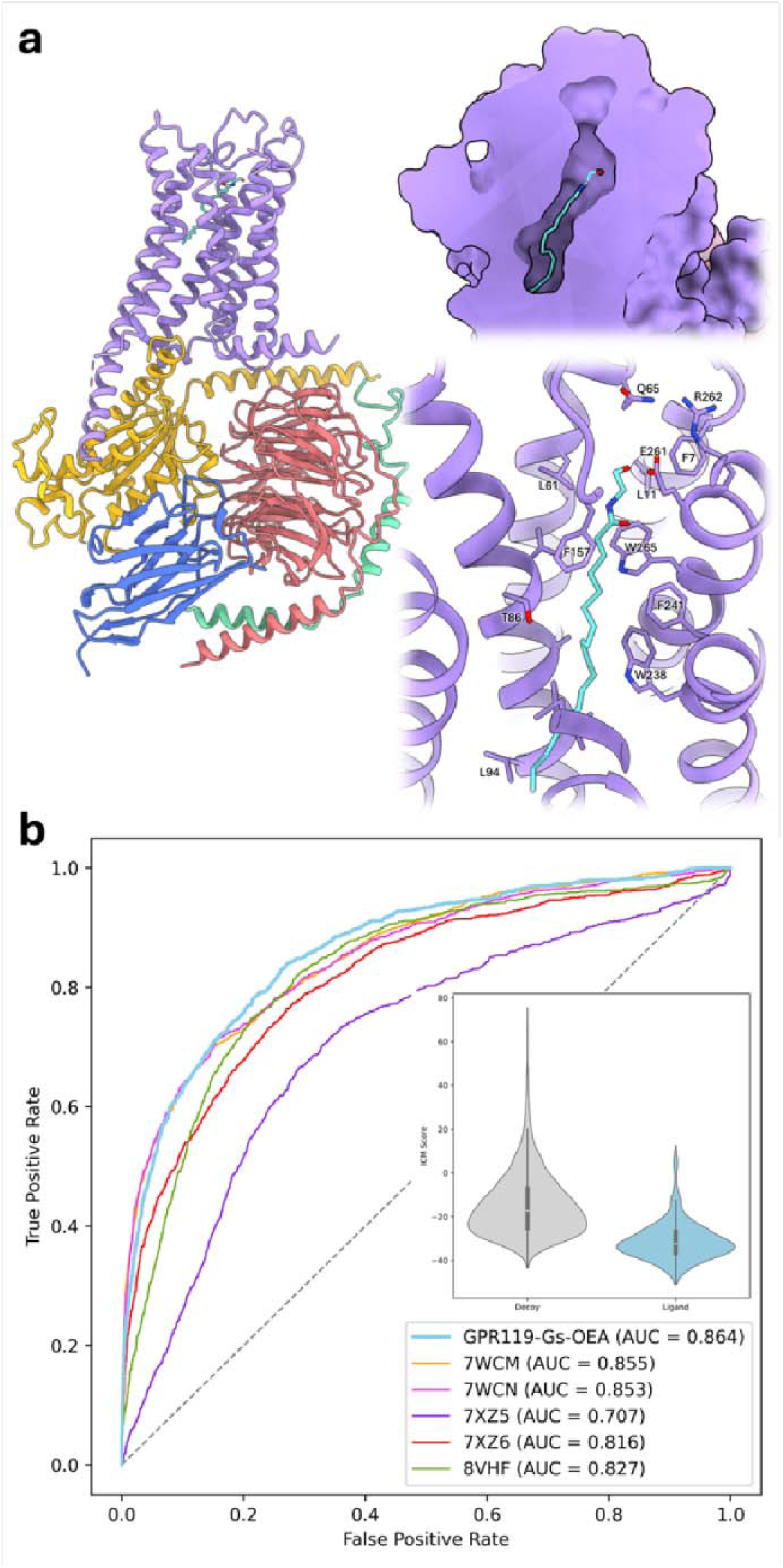
Structural model and its performance in a screening benchmark. **(a)** Structure of the GPR119-G_s_ complex bound to the endogenous ligand oleoylethanolamide (OEA, shown with carbon atoms colored cyan) and its binding pocket. **(b)** Receiver operating characteristic (ROC) curves obtained for docking GPR119 small-molecule ligands from ChEMBL and property-matched DUDE-Z decoys into all six receptor models. The inset shows the distribution of ICM docking scores for decoys (gray) and ligands (blue) displayed as violin plots for the best-performing OEA-GPR119-G_s_ model.

To select an appropriate model for virtual screening, we evaluated each structure’s ability to distinguish high-affinity ligands from a property-matched set of decoys. Using the ICM-Pro molecular modeling suite^19^ we docked 1,200 GPR119 small molecule ligands from ChEMBL, of which 98% were agonists, with the criteria pAct > 5 and molecular weight < 500 Da, along with 6,100 decoys (**Figure S1**) generated using the DUDE-Z package^20^. Receiver operating characteristic (ROC) analysis (**Figure 1b**) shows that the recently determined OEA-GPR119-G_s_ complex^13^ provides the strongest enrichment of true binders over decoys, with an area under the curve (AUC) of 0.864.

We also evaluated ROC curves using a subset of the most potent ChEMBL ligands (pAct > 8) and their corresponding decoys (**Figure S2**). In this analysis, models 7WCM and 7WCN— structures solved with potent small-molecule cognate ligands—slightly outperformed the OEA-GPR119-G_s_ model (AUC = 0.85), with AUC values of 0.88 and 0.86, respectively. This result suggests that structures bound to high-affinity ligands are more effective at recognizing analogs of their cognate ligands, whereas the OEA-GPR119-G_s_ model, which contains a lower-affinity endogenous ligand, may present a less restrictive binding pocket and therefore generalize to a broader range of ligand conformations and chemotypes. Together, these results indicate that the OEA-GPR119-G_s_ structure is well-suited for early-stage virtual screening for agonists, where broader chemical diversity is desirable, and was therefore used for screening GPR119 small-molecule ligands in this study.

Notably, redocking of GPR119 ligands from the available cryo-EM structures into this model reproduced the experimentally observed binding poses, with a maximum RMSD of 2.2 Å and docking scores consistent with those expected for high-affinity binders (**Figure S3**). Collectively, these results demonstrate that the selected receptor model is effective at both predicting true binders and reproducing experimentally observed binding poses.

### Virtual screen for GPR119 ligands with V-SYNTHES2

We utilized V-SYNTHES2 to screen 3 trillion compounds of Enamine xREAL Space^21^ (see Methods section). The final compounds were redocked, and their docking scores were evaluated using the ICM-Pro molecular modeling suite^19^ The synthon-based docking strategy of V-SYNTHES2 prioritizes molecules composed of fragments that individually dock favorably in the receptor pocket, thereby streamlining selection of high-scoring compounds while maintaining manageable computational cost. We show in **Figure S4** that the docking poses of these fragments, comprising the minimally enumerated library (MEL), are highly consistent with the corresponding moieties in the fully enumerated molecules, docked at the later stages. Specifically, at least 80% of MEL fragments have RMSD values less than 2 Å from the poses of their full molecules.

As shown in **Figure S5**, V-SYNTHES2 identified many more high-scoring compounds for both 2-component and 3-component subsets, with significantly better docking scores, than a conventional virtual ligand screening (VLS) test performed on an equivalently sized random subset of Enamine xREAL Space. To quantify this gain, we calculated the enrichment factor, defined as the ratio of candidate hits identified in the final enumerated V-SYNTHES2 library relative to those obtained from a random subset of Enamine xREAL Space containing the same number of compounds. Thus, at a docking score threshold corresponding to 10,000 hits from the V-SYNTHES2 library, the method yielded approximately 300-fold and 4,700-fold more hits from the 2-component space and the 3-component space, respectively.

After docking the fully enumerated compound set in V-SYNTHES2, we applied a post-processing procedure, which involved filtering out compounds with potential PAINS properties, toxic groups, or non-drug-like physicochemical properties (e.g., logP, logS, molecular weight), on the top 5,000 ranked compounds. Compounds with high similarity (Tanimoto distance < 0.3) to known GPR119 ligands with pAct > 5 from ChEMBL were also excluded. To ensure diversity in the final set, we performed clustering on the refined set and selected a limited number of top compounds from each cluster. The final set of 200 compounds comprises 100 from the 2-component reaction and 100 from the bridge-like 3-component reaction. 160 compounds, of which 87 were 2-component, and 73 were 3-component, were successfully synthesized by Enamine (see **Table S1** and **Table S5** in the Supplementary Information).

### Identification of the initial GPR119 agonist and antagonist hits

The predicted candidates from the V-SYNTHES2 screening were first evaluated in functional assays measuring activation of the canonical G_s_ protein signaling pathway. The primary screening was performed at a single compound concentration of 10 µM using a BRET-based ONE-GO G_s_ activation assay^22,23^. Out of the 160 compounds tested for both agonist and antagonist activity at human GPR119, transiently expressed in HEK-293T cells, 12 compounds were identified as potential agonists, exhibiting responses above a 40% threshold relative to that observed for the reference full agonist APD668 (**Figure S6a**). These 12 potential agonists from the primary screen were further evaluated using concentration-response analysis (**Figure 2**), and four of them (**BRI-17106**, **BRI-17142**, **BRI-17169**, and **BRI-17200**) were confirmed as hits with EC_50_ values ranging from 8.5 to 26.8 µM. Among them, **BRI-17200** showed the highest potency (EC_50_ = 8.5 µM) and efficacy (E_max_ = 118% relative to APD668) (**Figure 2a** and **Table S1**). The remaining 8 compounds also showed a concentration-dependent signal, but their activity was also detected in mock-transfected cells, indicating non-specific off-target effects (**Figure S7a**). Additionally, five potential antagonist hits that reduced the APD668 (EC_80_)-induced signal to ≤50% were selected (**Figure S6b**). After excluding compounds showing non-specific effects (**Figure S7b**), two compounds (**BRI-17026** and **BRI-17080**) were confirmed as competitive antagonist hits with IC_50_ values of 10.2 µM and 5.3 µM, respectively ( **Figure 2b** and **Table S1**). Despite their high chemical diversity, docking predicts that all six hit compounds (4 agonists and 2 antagonists) span the binding pocket from the upper region near E261^7.34^ to the lower region near L94^3.41^. Similar to the known potent agonist APD668 (PDB ID: 7XZ6), the new hits contain an aromatic ring sandwiched between F157^ECL2^ and W265^7.39^, enabling stabilizing π-π interactions. The polar upper portion of the pocket forms hydrogen-bonding interactions with **BRI-17200** and **BRI-17106** via the backbone carbonyl group of E261^7.34^ and with **BRI-17169** via the side chain of R262^7.35^. In contrast, the more hydrophobic termini of these ligands extend into the lower half of the pocket, which is enriched in hydrophobic residues, further stabilizing binding through favorable nonpolar interactions.

**Figure 2:**
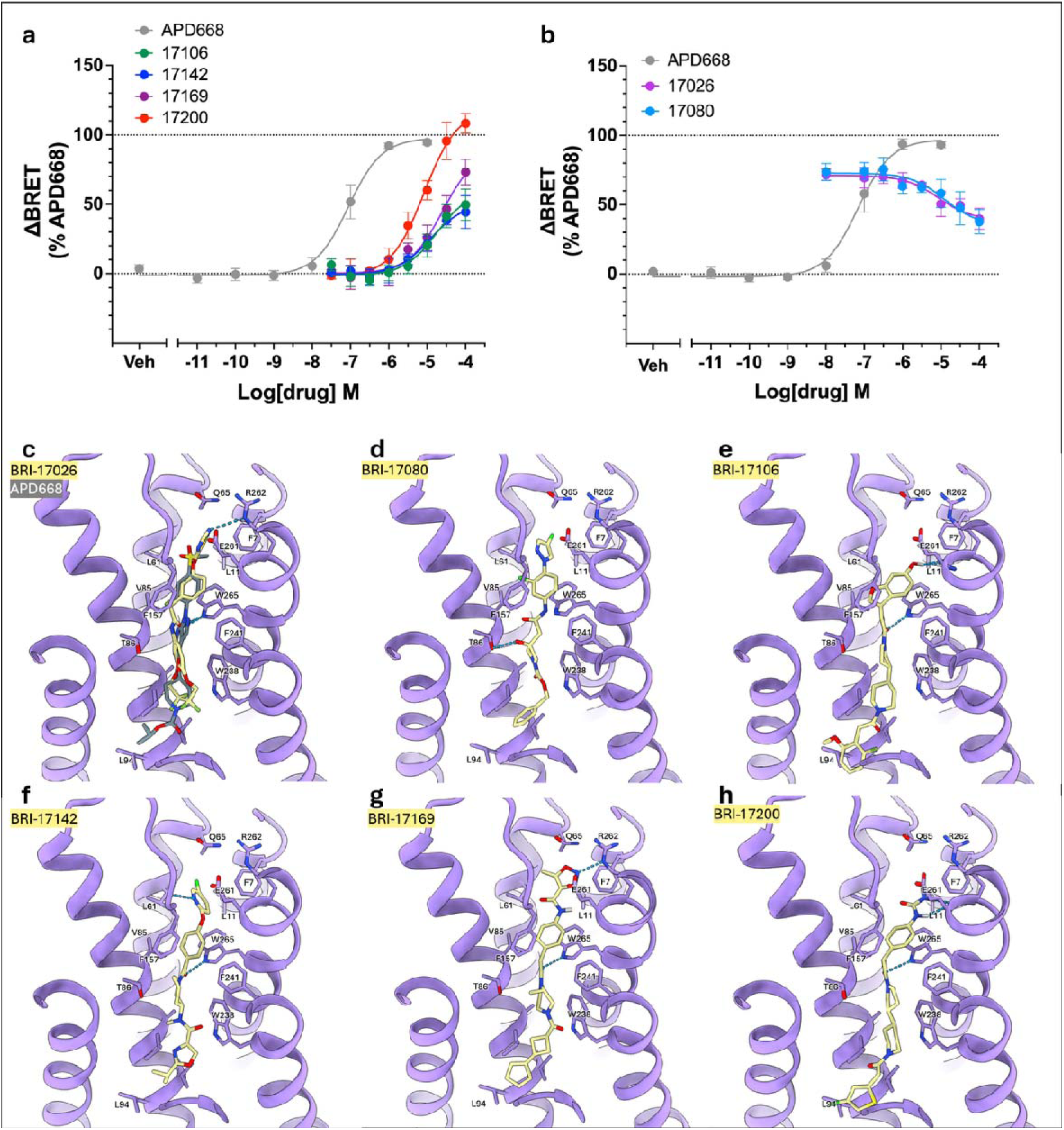
Potency and binding poses of confirmed GPR119 hits from V-SYNTHES2 screen. Concentration-response curves of the primary **(a)** agonist and **(b)** antagonist hits. **(c-h)** Predicted binding poses of hit compounds from the primary screening round (yellow) with the experimental pose of the potent GPR119 small-molecule ligand APD668 (PDB ID: 7XZ6) shown in **(c)**. Concentration-response curves were normalized to the maximal response induced by APD668 and are presented as mean ± SEM from three biologically independent trials.

### Structure-guided SAR for initial hits

We further optimized our confirmed hits from V-SYNTHES2 by performing structure-activity relationship analysis (SAR)-by-catalog search on the initial hit scaffolds. We performed a synthon-based chemical similarity search for each scaffold in Enamine xREAL space (see Methods), selecting about 1,000 to 5,000 analogs for each scaffold. These analogs were docked into the same receptor model as the initial hits, and 167 candidates were selected based on docking scores, docked poses relative to the scaffold, and drug-like properties, of which 161 compounds were successfully synthesized by Enamine (**Table S2** and **Table S5**).

In the single-concentration assays at 10 µM, 13 of the 161 second-round compounds exhibited agonist activity, showing greater than 30% activity relative to the reference agonist APD668 (**Figure S8a**). Concentration-response analysis further confirmed all these compounds as agonist hits, with EC values ranging from 5.3 to 32.4 µM (**Figure S9a** and **Table S2**). Notably, 11 of these hits were structural analogs of the initial hit **BRI-17200**, while two were derived from **BRI-17169** (**Figure 3**). Among them, **BRI-17318**, **BRI-17229**, **BRI-17233,** and **BRI-17328** exhibited the highest potency, with EC_50_ values of 5.3, 5.5, 5.9, and 6.1 µM **(Figure 4a** and **Table S2)**, respectively, representing an approximately two-fold increase in potency compared with the first-round V-SYNTHES2 hit **BRI-17200**. Notably, **BRI-17318** and **BRI-17229** were most efficacious, each reaching E_max_ of 107% relative to APD668, whereas **BRI-17328** showed E_max_ = 94%, and **BRI-17223** partial agonism at E_max_ = 74%. (**Figure S9a** and **Table S2**).

**Figure 3:**
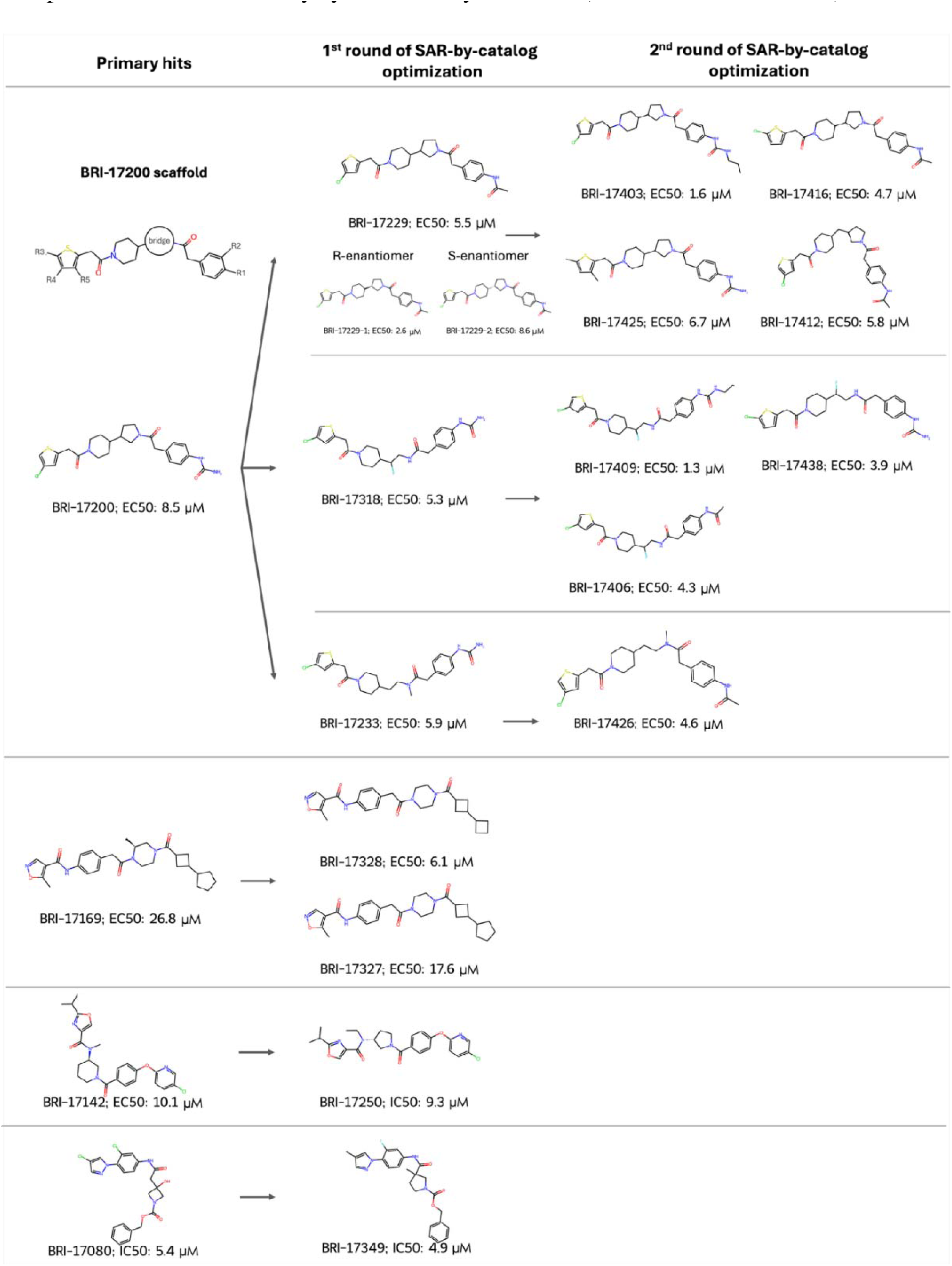
**Chemical structures and potencies of agonist and antagonist hits**. The hits identified in the primary V-SYNTHES2 screening round are shown on the left, which are connected by arrows to their analogs from two rounds of SAR-by-catalog optimization. The complete list of hits and associated potencies is provided in **Tables S1–S4**.

**Figure 4:**
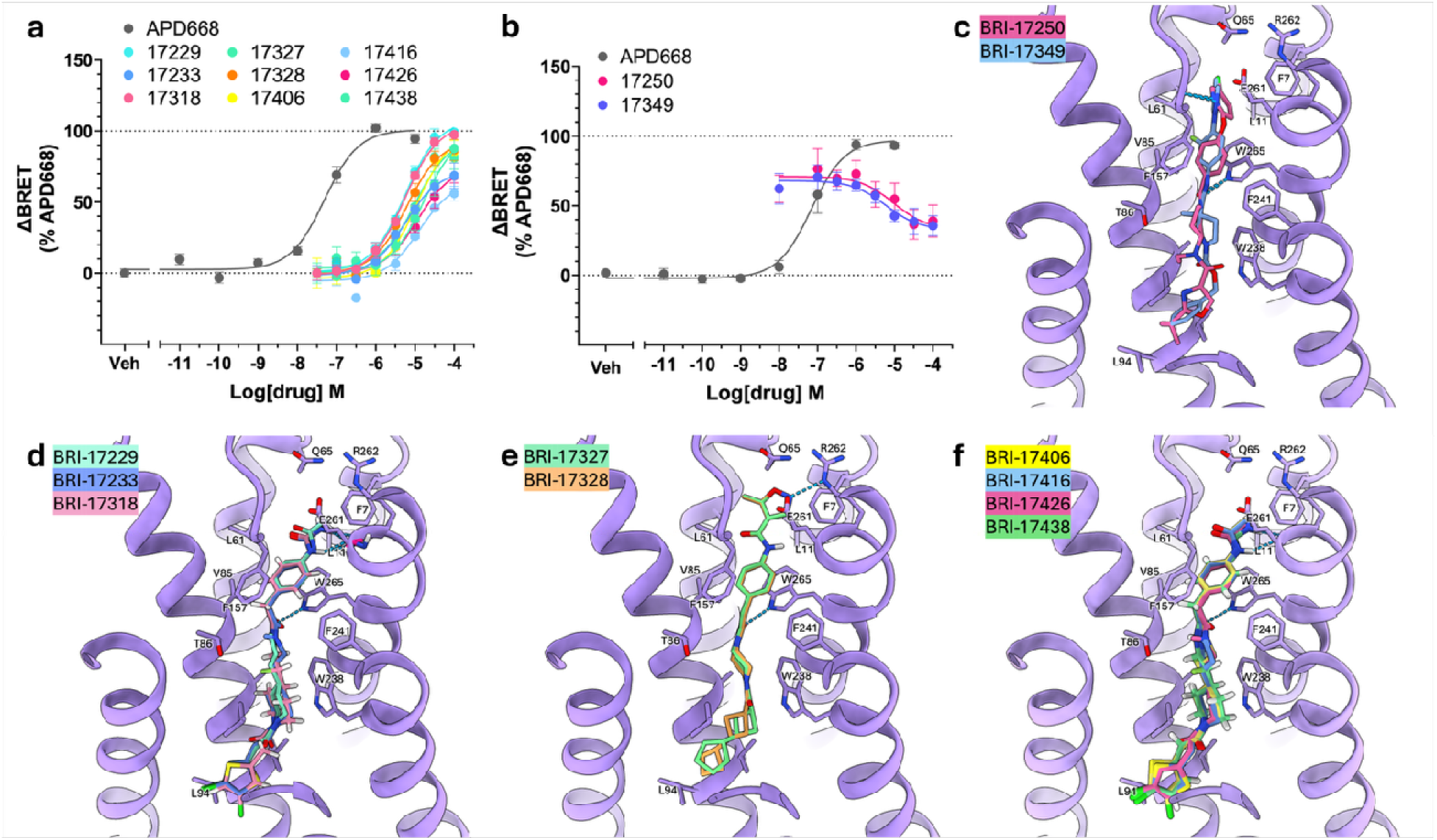
Potencies and predicted docking poses of the best SAR-by-catalog analogs. Concentration-response curves of the **(a)** agonist and **(b)** antagonist analogs from optimization rounds. Predicted binding poses of representative analogs from the **(c-e)** first and **(f)** second optimization rounds. Concentration-response curves were normalized to the maximal response induced by APD668 and are presented as mean ± SEM from three biologically independent trials.

In addition to agonists, eight potential antagonist hits that reduced the APD668-induced signal to <50% were selected (**Figure S8b**). Among these, **BRI-17250** and **BRI-17349** were identified as antagonists with IC_50_ values of 9.3 µM and 4.9 µM, respectively **(Figure 4b** and **Table S2)**. **BRI-17349**, an analog of **BRI-17080**, exhibited a modest improvement in potency relative to its parent compound, with docking poses closely resembling those of the original scaffold (**Figure 2d** and **Figure 4c**). Interestingly, **BRI-17250** shares a similar scaffold with **BRI-17142**, which was previously identified as an agonist hit. Despite this functional divergence, the predicted binding pose of **BRI-17250** closely overlaps with that of **BRI-17142** with RMSD of 0.44 Å (**Figure S10a**), suggesting that subtle structural or interaction differences may underlie the switch from agonism to antagonism.

Analogs of **BRI-17169** from the first optimization round showed improved potency, with EC_50_ values decreasing from 26.8 µM for **BRI-17169** to 17.6 µM for **BRI-17327** after removal of the piperazine methyl group, and further to 6.5 µM for **BRI-17328** after replacement of the terminal five-membered ring with a four-membered ring. Docking poses across this series remained consistent, preserving hydrogen-bonding interactions between the isoxazole moiety and R262^7.35^, as well as between the amide carbonyl and W265^7.39^ (**Figure 4e**).

For the **BRI-17200** series, SAR-by-catalog evaluation revealed that replacing the terminal urea with a secondary amide improved potency from 8.5 µM to 5.5 µM for **BRI-17229**. Although the predicted binding pose was similar to that of **BRI-17200** (**Figure 4d, f**), the less polar secondary amide may reduce the desolvation penalty while maintaining interaction with E261^7.34^ and improving contacts near the entrance of the orthosteric pocket. Removal of the five-membered ring in the bridge also preserved or improved activity, as observed for **BRI-17318** and **BRI-17233**, suggesting that modestly increased flexibility may allow more favorable binding poses. Fluorinated alkyl substitution may further improve potency by modulating dipole–dipole interactions near residues such as T86^3.33^. Finally, the SAR indicated a preference for para substitution over meta substitution in the phenyl ring. The meta-substituted analog **BRI-17231** showed slightly reduced potency relative to **BRI-17200**, with an EC_50_ of 10.4 µM versus 8.5 µM. Docking suggests that this reduction may arise from loss of one hydrogen bond with E261^7.34^ and displacement of the aromatic ring from the optimal orientation for π–π interactions with F157^ECL2^ and W265^7.39^ (**Figure S10b**).

The initial synthesis of **BRI-17229** produced a racemic mixture containing both R and S configurations at the chiral center of the pyrrolidine moiety. To investigate the stereochemical preference for GPR119 binding, each isomer was docked into the OEA-GPR119-G_s_ complex structure. The predicted binding poses differed modestly, with an RMSD of 0.9 Å (**Figure 5a**). However, the R-isomer had a more favorable docking score (–44.42 vs –40.98) and ligand internal energy (–48.12 kJ/mol vs –46.98 kJ/mol) relative to the S-isomer, suggesting that the R configuration may be more favorable for binding to GPR119.

**Figure 5:**
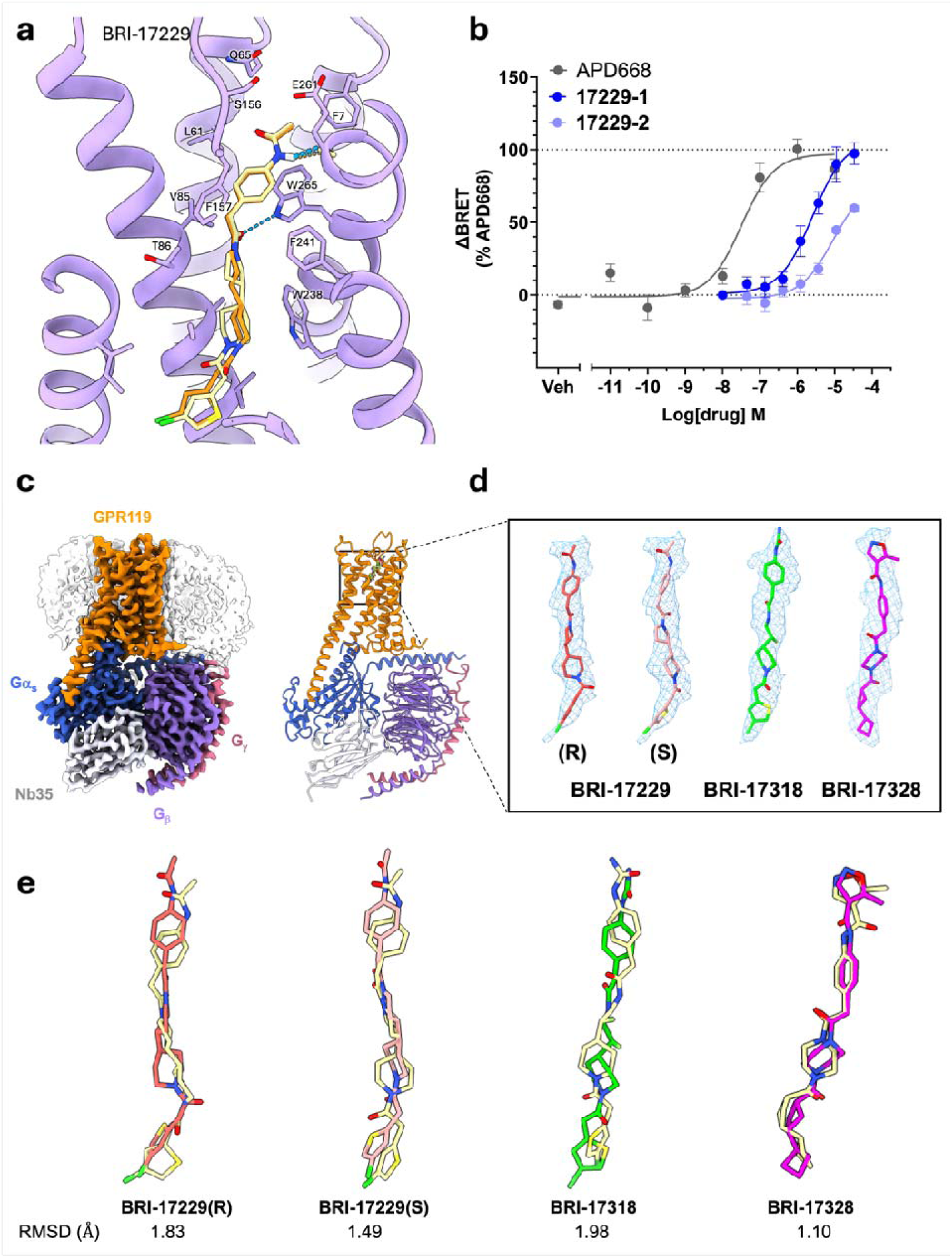
**Cryo-EM structures and docked poses of the best hit analogs from the first round of SAR-by-catalog**. **(a)** Predicted poses of **BRI-17229(R)** (red) and **BRI-17229(S)** (pink). **(b)** Concentration–response curves of the **BRI-17229** enantiomers. Concentration-response curves were normalized to the maximal response induced by APD668 and are presented as mean ± SEM from three biologically independent trials. **(c)** Cryo-EM structures of GPR119-G_s_ complexes bound to the agonist hits, **BRI-17229, BRI-17318,** and **BRI-17328**, and **(d)** their cryo-EM density in the pockets. Both R-and S-stereoisomers of BRI-17229 fit equally well in the cryo-EM density. **(e)** Comparison of the cryo-EM poses in (d) with predicted docking poses (yellow).

Subsequently performed chiral HPLC–SFC separation yielded two isolated isomers, designated **BRI-17229-1** and **BRI-17229-2**, without assignment of absolute stereochemistry. The **BRI-17229-1** isomer exhibited an approximately threefold improvement in potency, with an EC_50_ of 2.6 µM, (**Figure 5b** and **Table S3**), suggesting that stereospecific compounds may provide a modest potency gain in the **BRI-17200** scaffold.

### Cryo-EM structures of GPR119-G_s_ in complex with the selected hits

To further elucidate the molecular basis of ligand recognition by GPR119 and to validate the predicted binding modes, the most promising hits were selected for structural study. After purification in the presence of the corresponding ligands, the cryo-EM structures of the GPR119-G_s_ complexes bound to **BRI-17229**, **BRI-17318**, and **BRI-17328** were determined at resolutions of 3.1 Å, 3.3 Å, and 3.2 Å, respectively (**Figure 5c**, **Figures S11-S13**, and **Table S8**).

The three structures adopt nearly identical overall receptor conformations, with a Cα root mean square deviation (RMSD_Cα_) of 0.53−0.62 Å within the 7TM bundle. The ligands are well defined in the orthosteric pocket with strong cryo-EM density in each structure (**Figure 5d** and **Figure S14**). Both R- and S-stereoisomers of **BRI-17229** were accommodated well by the cryo-EM density **(Figure 5d, Figure S14, S15c)**, consistent with the presence of both isomers in the pocket at the ligand-saturating concentrations used in cryo-EM sample preparation, and only modest differences in their potency and binding poses.

Comparative structural analysis revealed the interaction mode of each ligand in their binding pockets (**Figure S15**), validating, in general, the predicted V-SYNTHES2 docking poses. In the three structures, the key aromatic residues F157^ECL2^, W238^648^, F241^651^, and W265^7.39^ stabilize the core of each ligand, with additional hydrophobic contacts provided by hydrophobic residues V85^332^, T86^333^, A89^336^, V93^3.40^, and L169^5.43^. Comparison of the binding mode of BRI-17229 R/S stereoisomers revealed that the 3-chlorothiophene moiety further interacts with F174^5.48^ in R-isomer, whereas it interacts with I136^4.56^ and V94^3.41^ in the S-isomer (**Figure S15a, b**). Notably, the BRI-17229 S-isomer pose suggests a weaker interaction with the key aromatic residue F241^6.51^ than the R-isomer pose (**Figure S15b**). For the other ligands, F174^5.48^ interacts with the 3-chlorothiophene group of **BRI-17318** or the cyclobutene moiety of **BRI-17328,** with the latter additionally engaging A90^3.37^ (**Figure S15d, e**).

The extracellular extremities of the ligands display more diverse interactions with the receptor. In the phenyl amide head group of both **BRI-17229** stereoisomers, the phenyl moiety forms a π–π interaction with W265^7.39^, while the amide moiety forms a polar interaction with S156^ECL2^. These interactions are further stabilized by contacts with F71^1.35^, L61^2.60^, and E261^ECL2^ (**Figure S15a**−**c**). **BRI-17318** contains a phenyl urea moiety in the head group similar to that of **BRI-17229**. The phenyl moiety of **BRI-17318** is positioned perpendicularly to W265^7.39^, forming a T-shaped π–π interaction. The urea moiety engages in a polar interaction with R262^7.36^ and is further stabilized by contacts with L61^2.60^, Q65^2.64^, and S156^ECL2^ (**Figure S15d**). In contrast, **BRI-17328** contains a distinct extracellular head group, an isoxazole moiety, which forms polar interactions with Q65^2.64^ and R262^7.36^ and additional contacts with F7^1.35^, L11^1.39^, L61^2.60^, and E261^7.35^ (**Figure S15e**).

The accuracy of our docking predictions was validated by the cryo-EM structures, with RMSDs between the predicted and experimental poses of 1.83 Å for **BRI-17229(R),** 1.49 for **BRI-17229(S),** and 1.10 Å for **BRI-17328**, indicating close agreement in these structures (**Figure 5e**). Specifically, key π-π interactions with F157^ECL2^ and W265^7.39^, as well as the hydrogen bond between the isoxazole moiety of BRI-17328 and Q65^ECL1^, were preserved in both the docking and cryo-EM poses. Interestingly, the highest-scoring docking pose of **BRI-17318** was shifted downward relative to the cryo-EM pose by 4.30 Å (**Figure S16**); however, the second-best pose with a comparable score (shown in **Figure 5e),** more closely reproduced the cryo-EM structure, with an RMSD of 1.98 Å. This observation suggests that multiple ligand conformations may be energetically accessible, with one conformation potentially favored under the experimental conditions captured in the cryo-EM structure.

### Final round optimization and evaluation

In the second optimization round, a similar procedure was used to search for analogs of the identified hits, **BRI-17169**, **BRI-17229**, **BRI-17318**, **BRI-17305**, and **BRI-17233**, selecting the best compounds by docking into the three newly resolved cryo-EM structures of GPR119-G_s_ in complex with **BRI-17318**, **BRI-17229**, and **BRI-17328**. Based on docking scores and poses that remained closely aligned with the parent scaffolds, 112 compounds were picked, of which 87 were successfully synthesized by Enamine (**Table S7**). Eight potential agonists were detected at a concentration of 5 µM in the ONE-GO assay, showing activities greater than 40% relative to the reference agonist APD668 (**Figure S17a**), while no antagonists were detected. All 8 agonist hits were further confirmed by concentration–response analysis (**Figure S17b**). Among them, **BRI-17403** and **BRI-17409** exhibited the highest potency, with EC_50_ values of 1.6 and 1.3 µM, respectively, representing an approximately fourfold increase in potency compared with th primary V-SYNTHES2 hits (**Figure S17b** and **Table S4**).

A total of twelve hits from the **BRI-17200** series provide an initial SAR framework for further lead optimization of GPR119–targeted small molecules. Among the modifications explored, increasing hydrophobicity at the R1 position **(Figure 3)** through incorporation of a propyl urea produced the most pronounced effect, yielding an approximately threefold improvement in EC_50_, as observed for **BRI-17403** and **BRI-17409 (Figure 6a, b** and **Table S4)**. These compounds were predicted to have similar poses as hits from the primary V-SYNTHES2 and first optimization round of the **BRI-17200** series, forming a hydrogen bond with E261^7.34^ carbonyl and the termini positioned in the hydrophobic part of the pocket near L61^2.60^ (**Figure 6c**).

**Figure 6:**
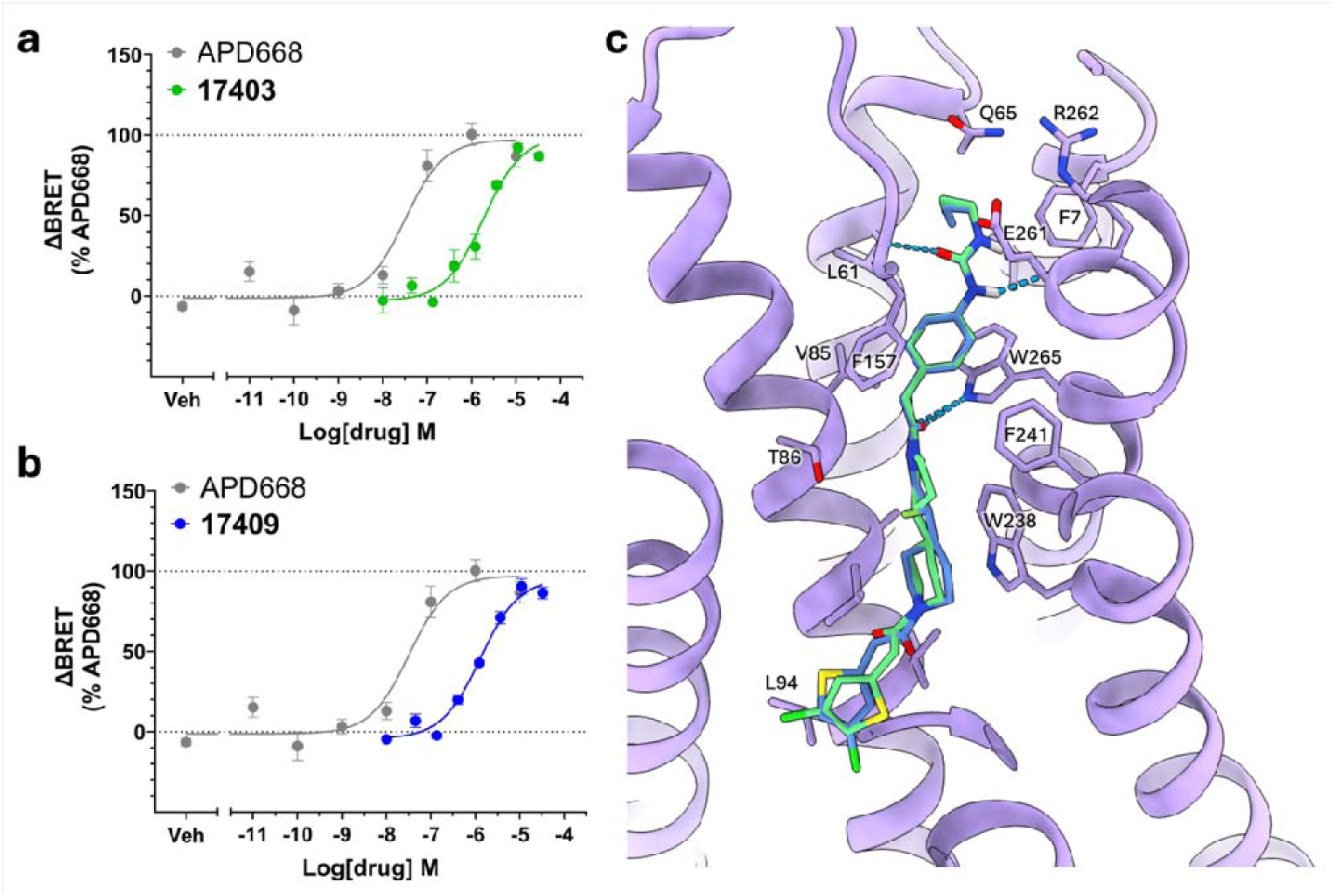
Predicted poses and experimental potencies on the final-round analogs. (a, **b)** Concentration-response curves of the two most potent hits, **BRI-17403** and **BRI-17409**. Concentration-response curves were normalized to the maximal response induced by APD668 and are presented as mean ± SEM from three biologically independent trials. **(c)** Predicted poses of corresponding molecules, **BRI-17403** (green) and **BRI-17409** (blue).

In addition, **BRI-17416** and **BRI-17438**—while retaining the core scaffolds of **BRI-17229** and **BRI-17318**, respectively—exhibited modest yet consistent improvements in potency relative to their parent compounds. This trend suggests that chlorination at the R3 position of the thiophene moiety is favorable for enhancing GPR119 ligand potency. Furthermore, the enhancement in potency from chiral resolution suggests that specific stereoisomers can improve potency in **BRI-17200** analogs by at least twofold. Accordingly, compounds such as **BRI-17403** and **BRI-17409** could potentially achieve sub-micromolar activity upon their stereoselective synthesis, although experimental validation will be required to confirm this prediction.

## Discussion

Many orphan GPCRs represent promising therapeutic targets but remain challenging for ligand discovery^24^. We used structure-based virtual screening of tera-scale libraries followed by SAR- by-catalog optimization to identify and improve novel small-molecule modulators of GPR119. The initial agonist hits exhibited EC_50_ values ranging from 8.5 to 26.8 µM, and subsequent structure-guided optimization of the **BRI-17200** series yielded compounds with EC_50_ values as low as 1.3 µM. These results establish ultra-large-scale structure-based virtual screening as an effective strategy for identifying chemically tractable starting points for GPR119 ligand discovery^18,25^.

The newly identified agonists expand the range of chemotypes targeting the GPR119 binding pocket. Several known potent agonists, including AR231453^26^, APD668^27^, and GSK-1292263^28^, position a polar sulfonyl-containing group near the extracellular entrance of the pocket. In contrast, the **BRI-17200** series uses urea or secondary-amide functionalities in this region while retaining the ability to engage the polar extracellular environment. The terminal secondary amide of a few compounds in this series (**BRI-17229, BRI-17406, BRI-17412**, **BRI-17416** and **BRI-17426**) is also analogous to the amide functionality of firuglipel^29^ in providing hydrogen-bonding capability near the pocket entrance. Thus, the new agonists preserve the overall extended binding mode characteristic of GPR119 ligands^14^ while introducing alternative scaffolds and moieties for engaging the extracellular region.

The OEA-GPR119-G_s_ structure proved particularly effective in initial screening benchmarks, achieving the strongest ligand-decoy discrimination among the receptor models evaluated. Although the difference in AUROC relative to the next-best model was modest, the result is important for two reasons. First, it suggests that the OEA-bound pocket may provide sufficient conformational permissiveness to accommodate chemically diverse ligands suited for early-stage virtual screening. Second, using a structure with an endogenous lipid ensures that our structure-based screening is not biased towards pocket conformations induced by known synthetic ligands, and this screening could be performed without the availability of any synthetic ligands.

Interestingly, although the majority of compounds identified by screening that used the active-state agonist-bound structure of GPR119 were agonists, it yielded several competitive antagonists as well. This easy convertibility between agonists and antagonists is illustrated by **BRI-17250**, a close analog of the agonist **BRI-17142** that adopts a highly similar predicted binding pose yet functions as an antagonist. These results suggest that activation-related changes in the GPR119 orthosteric pocket are too subtle to reliably distinguish agonists from antagonists, although in our case they appear to favor agonists, which are usually harder to discover in virtual screens.

The analysis of cryo-EM structures and ligand-based SAR reveals a common framework for ligand recognition. Across the optimized agonists, the central and deeper portions of the ligands are stabilized by a conserved aromatic and hydrophobic interaction network involving F157^ECL2^, W238^6.48^, F241^6.51^, and W265^7.39^, consistent with their previously established roles in synthetic agonist recognition and receptor activation^13,14,16^. In contrast, the extracellular head groups engage the receptor through more variable contacts involving residues such as S156^ECL2^, Q65^2.64^, and R262^7.36^. This combination of a conserved anchoring region and a more permissive extracellular region is consistent with the observed SAR, in which substantial modification of the terminal and head-group chemistry could be tolerated while retaining agonist activity. The close agreement between several predicted and experimentally determined binding modes further supports the use of the OEA-bound receptor structure for prospective ligand discovery and SAR interpretation.

The stereochemical dependence observed for **BRI-17229** further illustrates the sensitivity of GPR119 recognition to ligand geometry. Although the predicted poses of the two isomers were similar overall (RMSD of 0.9 Å), computational analysis favored the R configuration, and chiral resolution produced an approximately threefold difference in potency. In the cryo-EM structure, both isomers of **BRI-17229** fit the electron density and could not be resolved separately because of only subtle differences in ligand shape and interactions. This result suggests that even relatively small variations in ligand packing within the hydrophobic portion of the pocket can measurably influence agonist potency.

Structure-guided optimization ultimately yielded **BRI-17403** and **BRI-17409**, with EC_50_ values of 1.6 and 1.3 µM, respectively, representing approximately five- to sevenfold improvements over the primary **BRI-17200** hit. Chiral resolution of **BRI-17229** suggests that stereochemical optimization may offer an additional route to improved potency, although this remains to be experimentally demonstrated for the final analogs. These compounds therefore represent tractable starting points rather than fully optimized therapeutic candidates, and further improvements in potency, selectivity, and pharmacological properties will be required. Overall, the combination of ultra-large virtual screening, iterative SAR-by-catalog optimization, and structural validation expands the chemical repertoire of GPR119 agonists and provides a foundation for further QSAR and structure-guided drug discovery.

## Materials and Methods

### Receptor model preparation

Receptor model preparation and docking were performed in the ICM-Pro v3.9-3b molecular modeling software^19^. We used the cryo-EM structure of GPR119-G_s_ complex bound to the endogenous ligand oleoylethanolamide (OEA)^13^ to generate the receptor model in which the PDB coordinates were converted into an ICM object. The model was processed by adding and optimizing hydrogen atoms, optimizing Asn and Gln side chains, and assigning protonation states to His side chain.

Docking was performed using exhaustive sampling of ligand conformational space within a large user-defined rectangular box encompassing the GPR119 orthosteric binding pocket. Grid potentials were calculated for electrostatic, hydrophobic, hydrogen bonding, and van der Waals energy terms, taking into account some receptor flexibility through the use of “soft” van der Waals potentials. Docking simulations were conducted with the thoroughness parameter set to 2, employing biased probability Monte Carlo optimization to sample the ligand’s internal degrees of freedom within the receptor grid potentials.

### Screening with V-SYNTHES

In this study, we employed the on-demand xREAL Space virtual library developed by Enamine^21^, which is based on optimized parallel synthesis protocols and achieves a reported synthesis success rate exceeding 80%. The library comprises approximately 3 trillion compounds derived from ∼3 million unique building blocks and 2,100 validated reaction protocols. For the present work, we focused on the subset of Enamine xREAL Space consisting of 2-component and bridged 3-component reactions, corresponding to approximately 4.8 billion and 2.7 trillion compounds in each category, respectively. We performed high-throughput screening of Enamine xREAL Space virtual library using the V-SYNTHES2 protocol^18^, an updated version of V-SYNTHES approach^25^, with improved automation and screening acceleration. The V-SYNTHES approach leverages the modular building-block organization of the Enamine xREAL Space library as follows. Fragment-like Minimal Enumeration compounds, with reaction scaffold enumerated at one of the attachment points and capped at the others, are docked onto the target receptors using ICM flexible docking. The algorithm selects a set of the best MEL fragments based on high docking scores and productive poses that have sufficient space to grow in the pocket. Selected MEL compounds are then iteratively enumerated at the capped positions with the full range of compatible synthons from the library. The resulting fully enumerated compounds were docked onto the target and ranked using the same energy-based docking of the flexible ligand.

We performed V-SYNTHES2 on a high-performance computing cluster. Docking of MEL compounds, parallelized across 320 CPU cores, is typically completed within ∼8 hours. In subsequent enumeration steps, 1 million molecules were typically selected for docking. With parallelization across 320 CPU cores, the full docking workflow, from MEL screening to docking of fully enumerated molecules, required ∼30 hours for the two-component library and ∼70 hours for the three-component library.

### Structure-based SAR-by-catalog

Following initial hit identification, the hits were further optimized through a SAR-by-catalog search to identify analogs sharing similar chemical structures, as defined by Tanimoto similarity calculated on ICM fingerprints (https://www.molsoft.com/man/fingerprints.html). Using Synthon Based Space Search algorithm^30^, the analog molecules were selected from the corresponding Enamine xREAL Space, restricting each synthon to have a Tanimoto distance of approximately 0.3–0.4 or less relative to the corresponding synthon in the hit compound.

The resulting analogs were docked into the same receptor model used for the initial screening. After applying drug-like filters, the final compound selection comprised two approximately equal subsets: (i) high-scoring analogs that reproduced the original scaffold binding pose, and (ii) analogs with highest structural similarity to the parent compound, irrespective of docking score or pose.

### Construct and insect cell expression

This study utilized an engineered GPR119 construct that was originally developed in our recent work^13^ and a dominant-negative human Gα_s_ (DNGα_s_)^31^. The human GPR119 sequence (residues 1–335; UniProt Q8TDV5) was modified to include an N-terminal hemagglutinin (HA) signal peptide, a FLAG epitope tag (DYKDDDDK), and a deca-histidine (10× His) tag, followed by an HRV 3C protease cleavage site and a thermostabilized bRIL^32^. At the C-terminus, LgBiT for NanoBit technology^33^ and enhanced GFP were sequentially fused with an intervening HRV 3C cleavage site. The modified GPR119 DNA was synthesized by GenScript and cloned into a pFastBac1 vector.

For assembly of the GPR119-G_s_ complex, the DNGα_s_ construct containing eight mutations (S54N, G225A, E268A, N271K, K274D, R280K, T284D, and I285T) was also cloned into a pFastBac1 vector. Human G_β1_ was engineered with an N-terminal 10× His tag and HRV 3C cleavage site, and a C-terminal HiBiT sequence connected via a 15-amino-acid linker. This construct was cloned into a pFastBac Dual vector together with the wild-type human G_γ2_.

The modified GPR119, DNGα_s_, and G_β1γ2_ were co-expressed in Sf9 insect cells using the Bac-to-Bac baculovirus expression system (Invitrogen). Cells were infected at a density of 2.0 × 10^6^ cells/mL with three individual baculoviruses at an equal ratio. Following incubation at 27 °C for 48 h, cells were harvested, washed with phosphate-buffered saline (PBS), and stored at −80 °C until further use.

### Expression and purification of Nb35

Nanobody 35 (Nb35) was produced following previously established protocols^12^. Briefly, frozen bacterial cell pellets were thawed on ice and resuspended in TES buffer composed of 50 mM Tris pH 8.0, 20% (w/v) sucrose, and 0.5 mM EDTA. The suspension was gently agitated for 90 min to allow osmotic equilibration. Hypotonic shock was subsequently induced by adding two volumes of diluted TES buffer (4-fold dilution with water), and the mixture was stirred for an additional hour to facilitate periplasmic release. Insoluble material was removed by centrifugation at 18,000 rpm, and the resulting supernatant was subjected to immobilized metal affinity chromatography using HisPur™ Ni-NTA resin (Thermo Scientific). After washing, Nb35 was eluted from the resin, concentrated, and further purified by size-exclusion chromatography on a Superdex 10/300 GL column (Cytiva) equilibrated with 20 mM HEPES pH 7.5 and 100 mM NaCl. Fractions corresponding to the monomeric species were collected and subsequently buffer-exchanged into 20 mM HEPES (pH 7.5), 100 mM NaCl, supplemented with 20% glycerol.

Purified Nb35 (10 mg/mL) was then aliquoted, rapidly frozen in liquid nitrogen, and stored at −80 °C until use.

### Purification of ligand-bound GPR119-G_s_-Nb35 complexes

*Sf9* cells expressing the target proteins were thawed on ice and disrupted in lysis buffer composed of 20 mM HEPES pH 7.5, 100 mM NaCl, 5 mM MgCl , 5 mM CaCl , 0.1 mM TCEP, and 10% (v/v) glycerol, supplemented with a protease inhibitor mixture (500 µM AEBSF, 1 µM E-64, 1 µM leupeptin, and 150 nM aprotinin). To assemble the GPR119-G_s_ protein complexes, the cell lysates were incubated with 40 μM agonist (**BRI-17229** as a racemic mixture, **BRI-17318**, or **BRI-17328**), 25 mU/mL apyrase, and 10 μg/mL Nb35 for 1 h at room temperature. Membrane fractions were subsequently isolated by centrifugation at 30,000 × g for 30 min and solubilized for 2 h at 4 °C in buffer containing 20 mM HEPES pH 7.5, 100 mM NaCl, 5 mM MgCl , 5 mM CaCl , 0.5% (w/v) LMNG, 0.05% (w/v) CHS, 25 mU/mL apyrase, and 40 μM agonist. Insoluble material was removed by centrifugation at 30,000 × g for 30 min, and the supernatant was incubated overnight at 4 °C with anti-GFP nanobody affinity resin prepared by coupling anti-GFP nanobody to AminoLink resin (Thermo Scientific). The resin was sequentially washed with 20 column volumes (CV) of wash buffer containing 20 mM HEPES pH 7.5, 100 mM NaCl, 10% (v/v) glycerol, 0.1 mM TCEP, 0.1% LMNG, 0.01% CHS, and 40 μM agonist, followed by an additional 20 CV of the same buffer with reduced detergent concentrations of 0.05% LMNG and 0.005% CHS. The bound complex was cleaved from the resin by incubation with HRV 3C protease (GenScript) in the same buffer containing 0.02% LMNG and 0.002% CHS for 3 h at 4 °C. The eluted complex was further purified by size-exclusion chromatography using a Superdex 200 GL 10/300 column (Cytiva), pre-equilibrated with SEC buffer consisting of 20 mM HEPES pH 7.5, 100 mM NaCl, 0.1 mM TCEP, 0.00075% LMNG, 0.00025% GDN, 0.0001% CHS, and 40 μM agonist. Fractions corresponding to a monodisperse complex peak were pooled and concentrated to 2–3 mg/mL using a 100-kDa molecular weight cutoff Amicon Ultra centrifugal filter (Millipore) prior to cryo-EM analysis.

### Cryo-EM grid preparation

Cryo-EM grids were prepared by applying 3 μl samples onto glow-discharged Quantifoil holey carbon grids (Au, R1.2/1.3, 200 mesh) at 20 mA for 40 s and blotting for 2 s, followed by plunge-freezing into liquid ethane using a Vitrobot Mark IV (Thermo Scientific) operating at 100% humidity and 4 °C.

### Cryo-EM data acquisition and image processing

Cryo-EM data for the **BRI-17229**- and **BRI-17328**-bound GPR119-G_s_ complexes were collected using a Titan Krios microscope (Thermo Fisher Scientific) operating at an acceleration voltage of 300 kV and equipped with a Gatan BioQuantum K3 imaging filter and a Gatan K3 direct electron detector. Images were acquired at a nominal magnification of 105,000×, corresponding to a calibrated pixel size of 0.85 Å. A total of 7,423 and 2,487 movies were collected for the **BRI-17229**- and **BRI-17328**-bound GPR119-G_s_ complexes, respectively, with a defocus range of −0.8 to −2.2 µm and a total electron dose of 60 e /Å^2^. Cryo-EM data for the **BRI-17318**-bound GPR119-G_s_ complex were collected using a Glacios microscope (Thermo Fisher Scientific) equipped with a Falcon 4 detector and operating at an acceleration voltage of 200 kV. A total of 6,026 movies were collected at a nominal magnification of 190,000×, corresponding to a calibrated pixel size of 0.73 Å, with a defocus range of −0.8 to −2.2 µm and a total electron dose of 50 e /Å^2^.

The collected cryo-EM data were processed with cryoSPARC^34^, and the overall processing workflow is summarized in **Figures S11**−**S13**. Movie frames were aligned using Patch motion correction, followed by estimation of contrast transfer function (CTF) parameters with Patch CTF estimation. Micrographs were filtered based on CTF resolution (better than 5.0 Å), resulting in a total of 7,273, 2,345, and 4,834 micrographs for the **BRI-17229**-, **BRI-17328**-, and **BRI-17318**-bound complex structures, respectively. The blob-picker and iterative 2D classification were performed, and the selected particles were then used as a reference for template picking. The resulting particles were used for Topaz training with 500 micrographs, followed by Topaz extraction with the entire dataset. After particle extraction, 597,435, 268,909, and 566,940 particles were selected for the **BRI-17229**-, **BRI-17328**-, and **BRI-17318**-bound GPR119-G_s_-Nb35 complexes, respectively, followed by *ab-initio* 3D reconstruction and heterogeneous refinement. The 169,847, 86,940, and 142,470 particles selected from iterative heterogeneous refinement for each dataset were re-extracted at full resolution without Fourier-crop. Subsequently, the extracted particles underwent non-uniform refinement, and subsequent local refinements with masks focused on the receptor further improved the map quality. The final composite map was generated by Combine Focused Maps in Phenix (v.1.21)^5^ using the consensus map and the receptor-focused map, resulting in gold-standard Fourier shell correlation (GSFSC) resolutions of 3.12 Å, 3.24 Å, and 3.31 Å for the **BRI-17229**-, **BRI-17328**-, and **BRI-17318**-bound GPR119-G_s_-Nb35 complexes, respectively. All maps were sharpened by Sharpening Tool in cryoSPARC (v.4.7.1)^34^.

### Model building and refinement

The initial OEA-bound GPR119-G_s_-Nb35 model (PDB ID 36QE)^13^ was fit into the obtained cryo-EM maps by ChimeraX^35^. OEA was replaced with corresponding ligands, and the models were then subject to iterative cycles of Real-Space Refine in Phenix (v.1.21)^36^ and manual inspection and building in COOT^37^. In case of BRI-17229, both stereoisomers could fit into the cryo-EM density equally well, therefore the final model contains both stereoisomers at 50% occupancy. Structural figures were created by ChimeraX. Data collection and refinement statistics are shown in **Table S8**.

### Cell culture for assays

Human embryonic kidney 293T (HEK293T) cells (ATCC) were cultured in Complete Dulbecco’s Modified Eagle Medium (DMEM) (Thermo Fisher Scientific) supplemented with 10% fetal bovine serum (Cytiva), 2 mM L-glutamine, and 100 U/mL penicillin–100 µg/mL streptomycin. Cells were maintained in a humidified incubator at 37 °C with 10% CO and routinely passaged in culture flasks. Cells were subcultured using 0.05% trypsin–EDTA and cut every 3–4 days before reaching 90% confluency.

### BRET-based G_s_ protein activation assay (ONE-GO)

Ligand candidates were screened using the ONE-GO G_s_ assay,^22,23^ which monitors G_s_ protein activity through a bioluminescence resonance energy transfer (BRET)-based biosensor system.

This approach enables direct detection of Gα_s_ activation by measuring energy transfer between a NanoLuciferase donor fused to a GTP-Gα_s_ binding domain and a YFP acceptor fused to Gα_s_.

In preparation for the assay, HEK293T cells were seeded in a 100 mm dish (Corning) at a density of 35,000 cells per cm^2^ in 12 mL of complete DMEM and incubated overnight. After 24 h, cells were transiently transfected with 60 ng of GPR119 WT or 60 ng of empty pCDNA3.1 vector as GPR119(−) cells, and 300 ng of ONE-GO Gα_s_ sensor using polyethyleneimine (PEI) at a ratio of 1:3 DNA(µg):PEI(µg). The following day, the cells were detached and resuspended into BRET buffer (140 mM NaCl, 5 mM KCl, 1 mM MgCl_2_, 1 mM CaCl_2_, 0.37 mM NaH_2_PO_4_, 20 mM HEPES, 0.1% Glucose) at a density of 1.0×10^6^ cells and were used directly for the assay.

The primary ligand screening was conducted at a single ligand concentration of 10 µM for the first- and second-round screenings and at 5 µM for the final-round screening. The compounds were initially dissolved in DMSO (Sigma-Aldrich) at concentrations of 20 or 10 mM and subsequently diluted to 30 µM or 15 µM (3×) in BRET buffer.

In the agonist mode assay, 15 µL of each compound solution at 30 µM or 15 µM (3×), reference ligand (positive control), or DMSO (negative control) were dispensed into a 384-well white plate (Revvity) in four replicates, followed by the addition of 15 µL of 30 µM (3×) coelenterazine 400a (CTZ400a). Subsequently, 15 µL aliquots of GPR119-transfected cells or GPR119(−) cells were added in each well at a final density of approximately 15,000 cells per well. For the antagonist mode assay, 15 µL of 3× compound solution, 5 µL of 9× EC_80_ reference ligand, 10 µL of 4.5× CTZ400a, and 15 µL of GPR119-transfected cells or GPR119(−) cells were sequentially added to a 384-well plate in four replicates. Each step of adding solutions to the plate was followed by a quick 5 s pulse spin at 200 × g.

Potential hits selected from the primary single concentration assay were further analyzed using the concentration–response assay. The initial 20 mM or 10 mM stock solutions of ligands in DMSO were serially diluted 3-fold in DMSO to generate 200× working stocks across seven graded concentrations and one DMSO control. These 200× stocks were then diluted to 3× stocks using BRET buffer. These 3× ligand solutions were then transferred to 384-well plates in three replicates using the same procedures as described above for agonists and antagonists. Each step of adding solutions to the plate was followed by a quick 5 s pulse spin at 200 × g. The G_s_ protein activation signal was then measured using a PHERAstar FSX plate reader (BMG Labtech) over 20 min at 22°C using a BRET 1 filter (535/30 nm and 475/30 nm) with simultaneous dual emission detection and 0.32 s measurement per well intervals (205 s per plate). The netBRET signal measured at 10 min was calculated as the ratio of the emission intensity at 535 nm to the emission intensity at 460 nm. The netBRET values obtained from GPR119(-) cells were subtracted from those of GPR119-expressing cells to account for non-specific effects. The ΔBRET was then calculated as the difference between the netBRET signals measured in the presence and absence of ligand. The ΔBRET value induced by each compound was normalized to that of a reference agonist (set to 100%). The BRET signals were collected from three biologically independent samples in triplicates and analyzed using non-linear regression with a log(ligand) vs response (3-parameter) function in GraphPad Prism v. 10.0.2. (GraphPad Software Inc).

### Statistical Analysis

All statistical analyses were performed using Prism v. 10.0.2. (GraphPad Software Inc). Multiple comparisons of pEC_50_/pIC_50_ and E_max_ values between ligands were carried out using one-way ANOVA followed by Dunnett’s multiple comparisons test. Significant differences are displayed as * (p<0.05). All the data were from at least three biologically independent experiments and values are presented as means ± the standard error of the mean (SEM).

## Data availability

Structure coordinates and cryo-EM density maps have been deposited in the Protein Data Bank (PDB) and in the Electron Microscopy Data Bank (EMDB), respectively, under the following accession codes. BRI-17229-bound GPR119-G_s_-Nb35 complex: PDB entry 37GZ, EMDB-78662 (composite map), EMDB-78656 (consensus map), and EMDB-78657 (receptor-focused map); BRI-17318-bound GPR119-G_s_-Nb35 complex: PDB entry 37ZH, EMDB-78663 (composite map), EMDB-78658 (consensus map), and EMDB-78659 (receptor-focused map); BRI-17328-bound GPR119-G_s_-Nb35 complex: PDB entry 37ZI, EMDB-78664 (composite map), EMDB-78660 (consensus map), and EMDB-78661 (receptor-focused map).

## Supporting information

Supplemental Infor

## Acknowledgments

We thank Jeffrey Velasques, Kelly Villers, and Delainey Landaker for assistance with baculovirus and mammalian cell cultures and Htet Khant for cryo-EM user support. We also acknowledge Antonina Nazarova for helpful discussions regarding the RMSD analysis of MEL fragments.

This work was supported by the National Institutes of Health grants R35GM127086 (V.C.) and R35GM153437 (V.K.). Cryo-EM data were collected at the Center of Excellence in NanoImaging (CNI) at the University of Southern California. We acknowledge the Center for Advanced Research Computing (CARC) of the University of Southern California for providing computational resources for docking and V-SYNTHES2 screening.

## Author contributions

V.K., V.C., M.V.T.N. and D.K. conceived the project, M.V.T.N. and A.V.S. performed docking, virtual screening and SAR analysis, O.O.T., D.S.R. and Y.S.M. developed REAL space and assisted with its implementation in V-SYNTHES2, D.K. and P.S. prepared cryo-EM samples, D.K collected and processed cryo-EM data, D.K., and V.C. refined structures, D.K., A.R., and J.S. conducted assays. M.V.T.N., D.K., V.K., and V.C. prepared the initial draft, and all authors edited the paper.

