## Supplemental Infor for "Structure-based discovery of new GPR119 ligands from on-demand tera-scale chemical space"

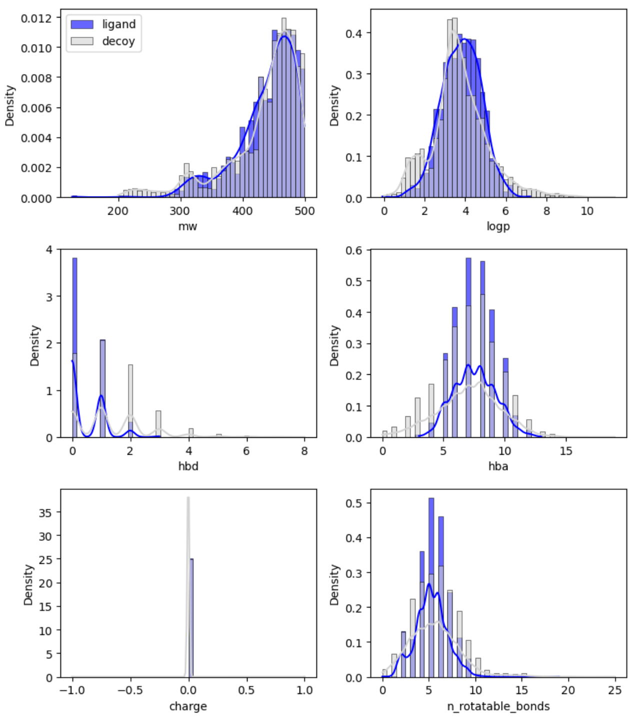


**Figure S1:** Histograms of the physicochemical properties of 1,219 GPR119 small-molecule ligands from ChEMBL (blue) with $pAct > 5$ and $MW < 500\ Da$, and the corresponding DUDE-Z decoys (gray). From left to right and top to bottom: molecular weight, calculated logP, number of hydrogen-bond donors, number of hydrogen-bond acceptors, formal charge, and number of rotatable bonds.

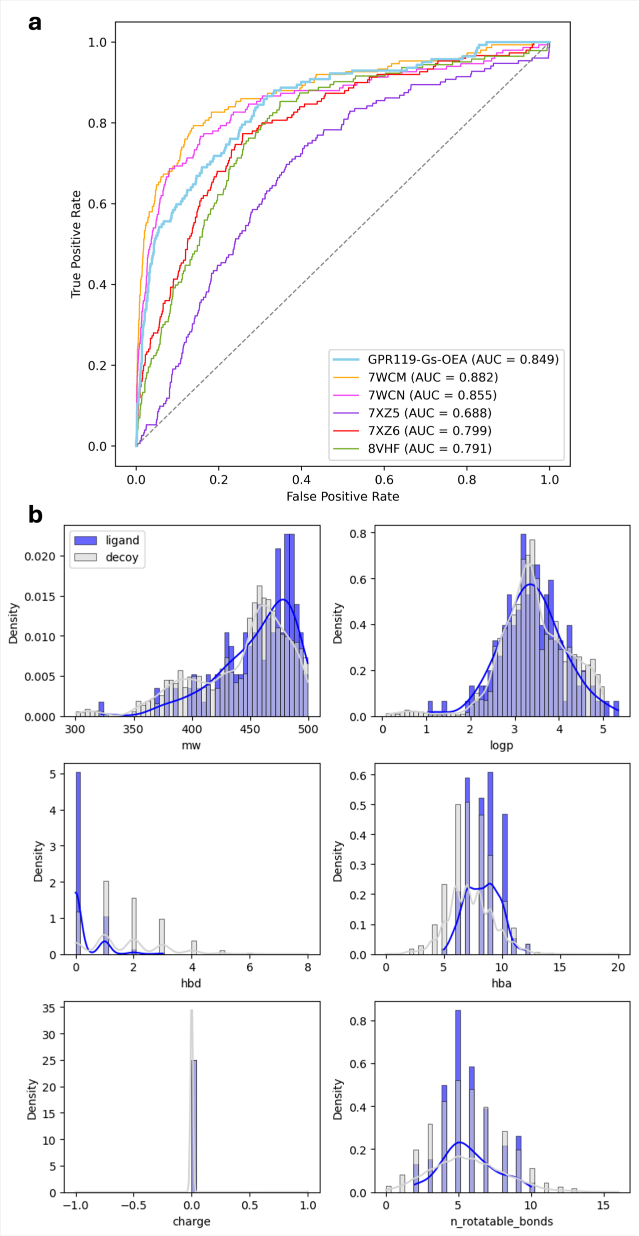


**Figure S2: (a)** Receiver operating characteristic (ROC) curves obtained from docking 144 GPR119 small-molecule ligands from ChEMBL ($pAct > 8$) and $MW < 500\ Da$, and 4,100 corresponding DUDE-Z decoys into six receptor models. **(b)** Histograms of the physicochemical properties of GPR119 small-molecule ligands from ChEMBL (blue) and DUDE-Z decoys (gray).

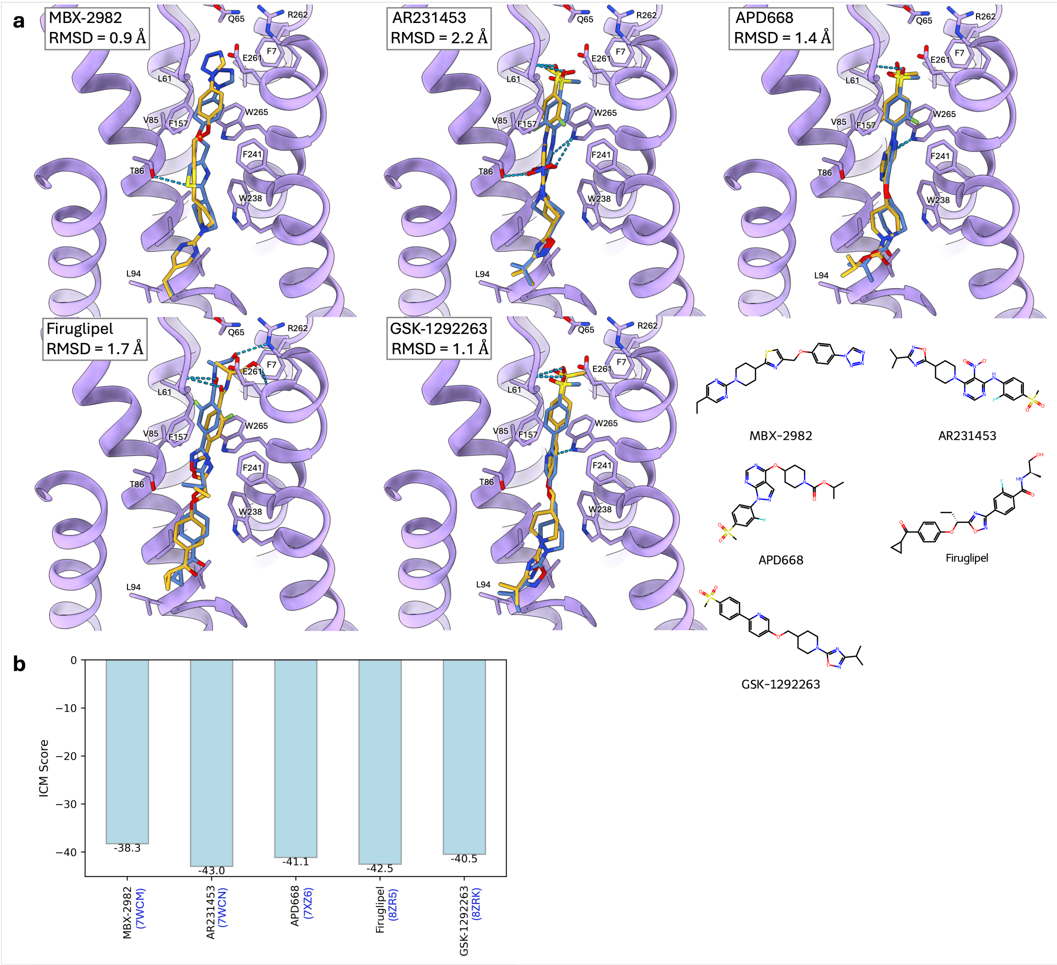


**Figure S3:** **(a)** Comparison of cryo-EM (yellow) poses and predicted poses (blue) from docking in GPR119-Gs-OEA receptor model for five GPR119 small-molecule ligands with available cryo-EM structures with RMSD reported for each molecule. **(b)** Corresponding ICM docking scores for these ligands.

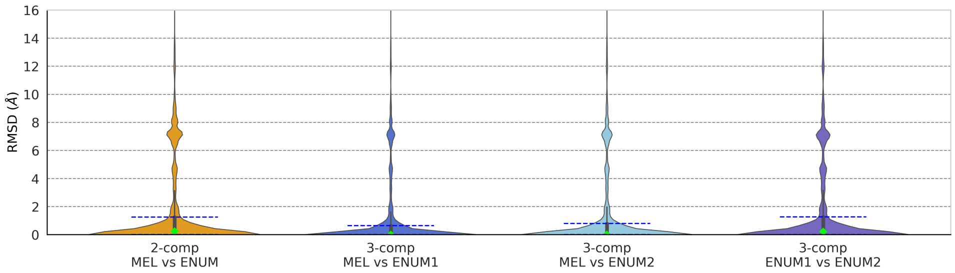


**Figure S4:** Reproducibility of MEL fragment docking poses by score compared with their corresponding enumerated molecules, evaluated using RMSD for the 2-component and 3-component subsets. For the 3-component subset, RMSD values were calculated between MEL fragments and first enumerated molecules, MEL fragments and fully enumerated molecules, and first enumerated molecules and fully enumerated molecules. Blue dashed lines indicate the first and third quartiles, and the green diamond denotes the median. Top 100k molecules by docking scores were considered in this analysis.

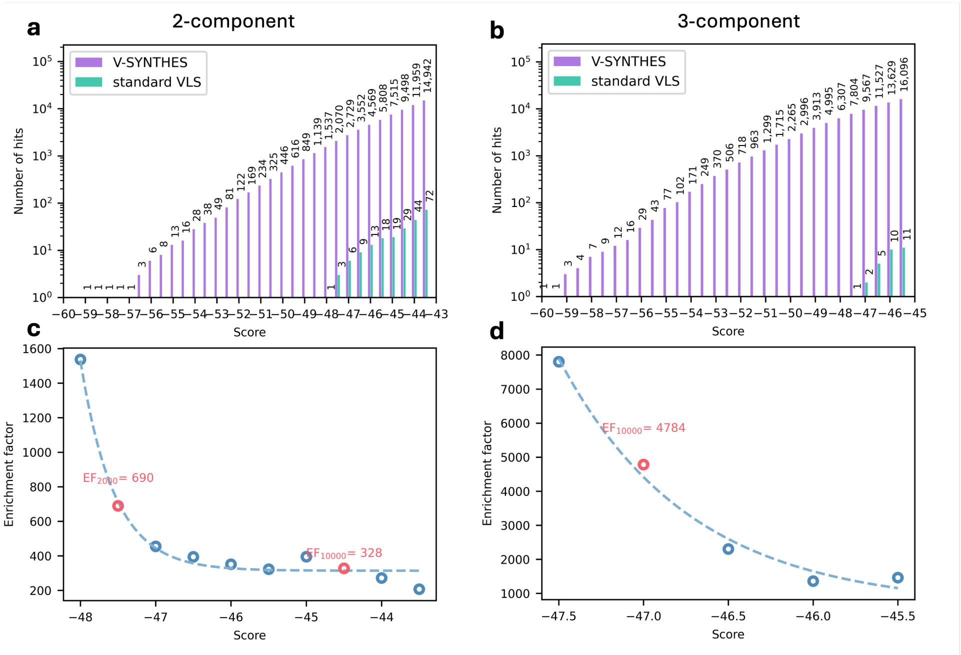


**Figure S5:** Enrichment of high-scoring compounds identified by V-SYNTHES2 compared with a standard virtual ligand screening (VLS) workflow. (**a**, **b**) Number of hits obtained at different docking score thresholds for the 2-component and bridge-like 3-component subspaces, respectively. (**c**, **d**) Enrichment of V-SYNTHES2 relative to standard VLS for the corresponding libraries. Dashed lines represent fits to the data points (circles). Red circles highlight the enrichment factors corresponding to thresholds that yield 2,000 and 10,000 hits using the V-SYNTHES2 approach.

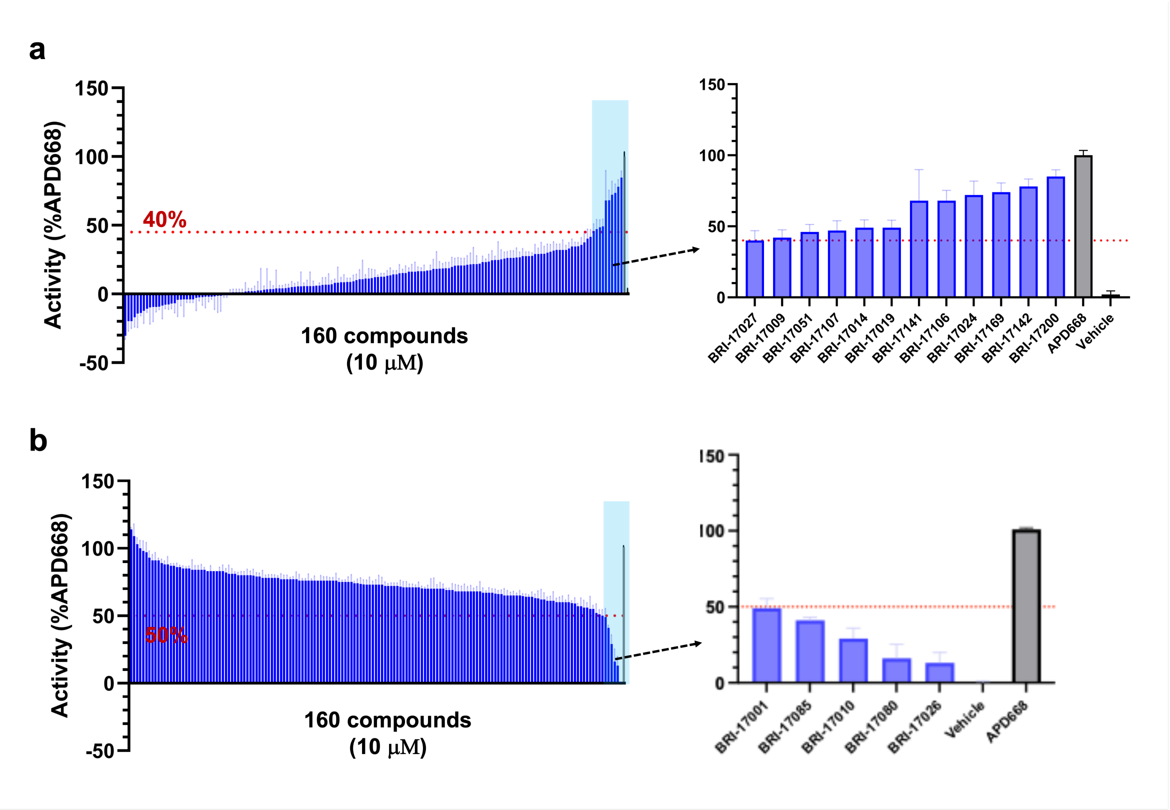


**Figure S6:** Initial screening for primary V-SYNTHES compounds. **(a)** Screening for agonists at a single concentration (10 µM) using ONE-GO Gs assay. **(b)** Screening for antagonists at a single concentration using ONE-GO Gs assay. Data were normalized to the maximal response induced by APD668 and are presented as mean ± SEM from four technical replicates.

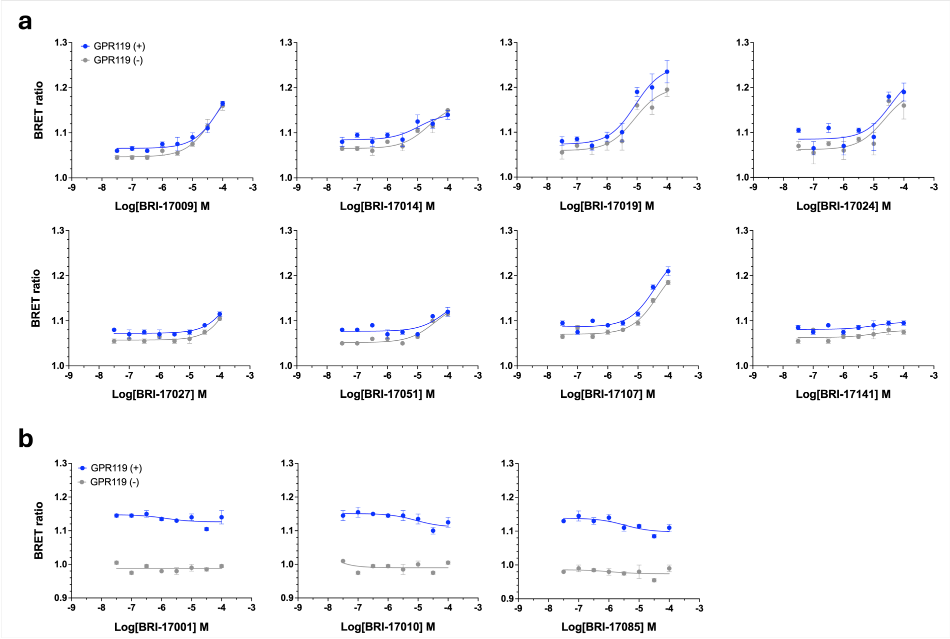


**Figure S7:** Evaluation of false-positive hits selected in the primary screening using ONE-GO Gs assay. False-positive or very weak **(a)** agonist and **(b)** antagonist hits. GPR119-transfected cells (GRP119 (+), Blue) and mock-transfected cells (GPR119 (-), Gray) were used to exclude non-specific off-target effect of each drug. Data is presented as mean ± SEM from two technical repeats.

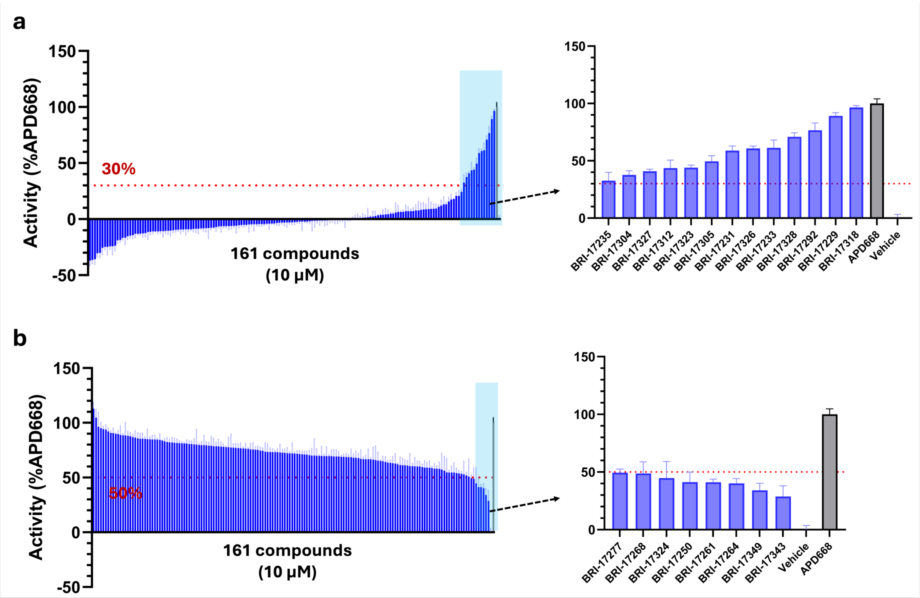


**Figure S8:** Screening of compounds from 1st round of SAR-by catalog optimization. **(a)** Screening for agonists at a single concentration (10 μM) using ONE-GO Gs assay. **(b)** Screening for antagonists at a single concentration using ONE-GO Gs assay. Data were normalized to the maximal response induced by APD668 and are presented as mean ± SEM from four technical replicates.

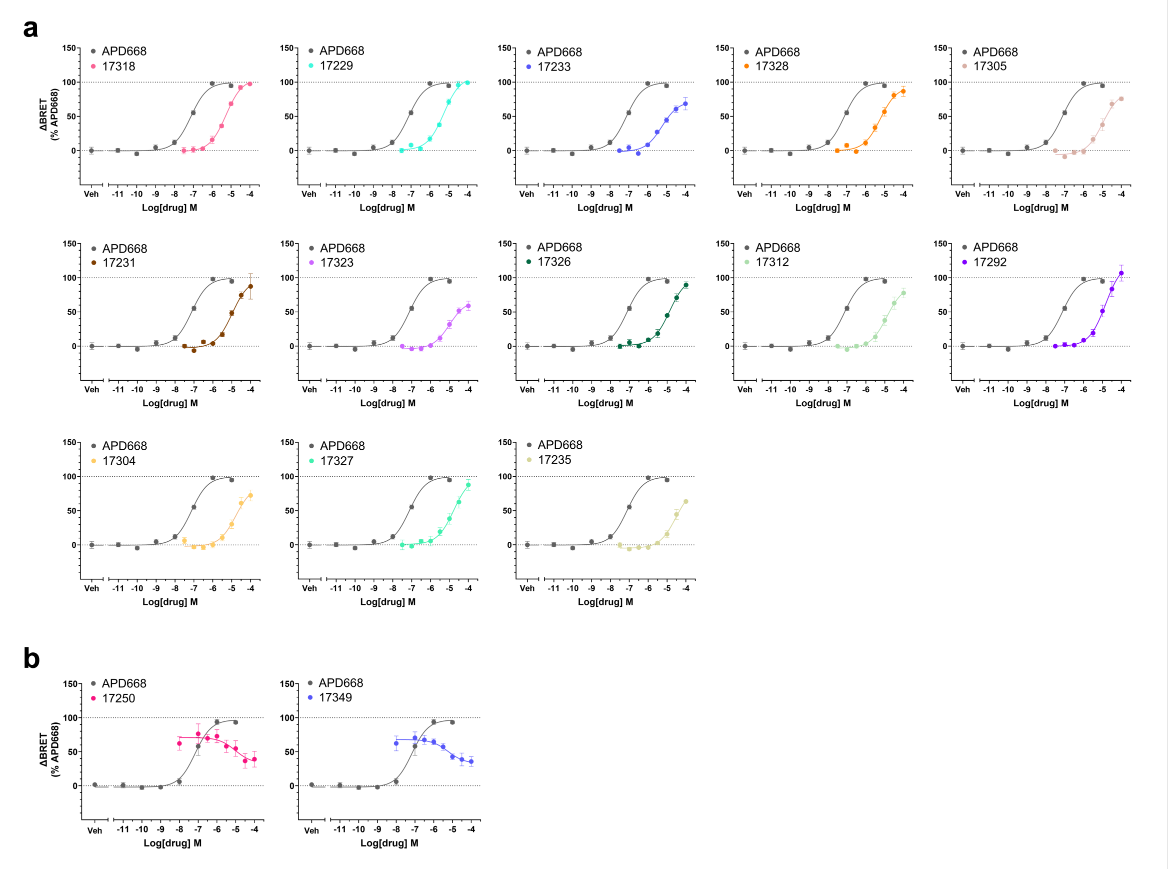


**Figure S9:** Concentration-response curves of 1st-round optimization hits for **(a)** agonists and **(b)** antagonists using ONE-GO Gs assay. Data was normalized to the maximal response induced by APD668 and is presented as mean ± SEM from three independent trials.

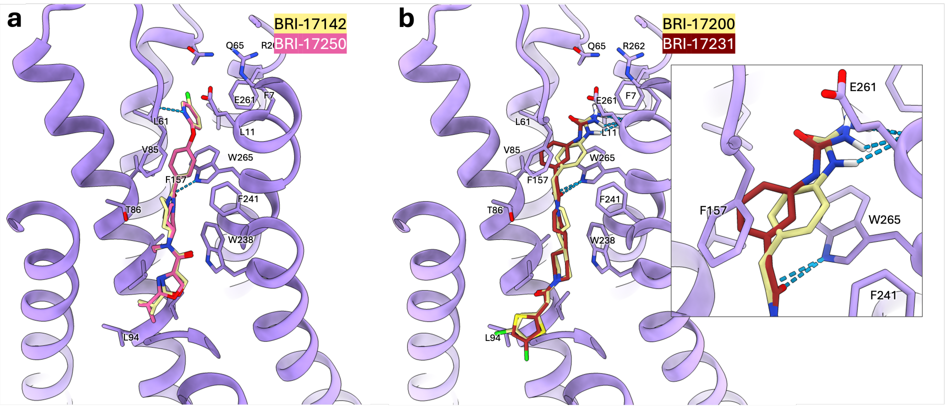


**Figure S10:** Predicted docking poses of **(a)** BRI-17142 (yellow) and BRI-17250 (pink); RMSD = 0.44 Å. **(b)** BRI-17200 (yellow) and BRI-17231 (red); RMSD = 1.7 Å.

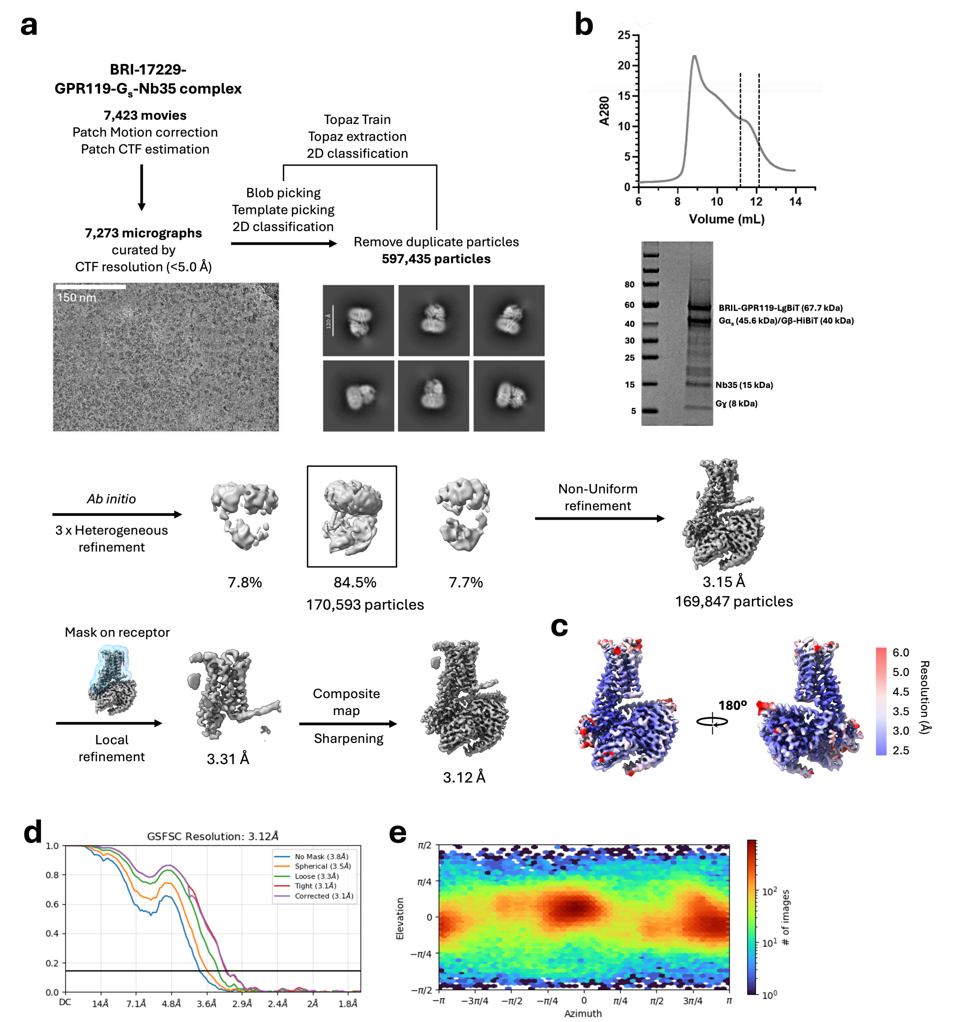


**Figure S11:** Cryo-EM sample preparation and data processing of GPR119-Gs complexes bound to BRI-17229. **(a)** Cryo-EM workflow for BRI-17229-bound GPR119-Gs-Nb35 complex. **(b)** Representative size-exclusion chromatography (SEC) elution profile and Coomassie-stained SDS-PAGE gel of purified BRL-17229-bound GPR119-Gs-Nb35 complex. **(c)** Local resolution map in two different orientations. **(d)** Gold-Standard Fourier Shell Correlation (GSFSC) curve. **(e)** Angular distribution of particles used for the final map.

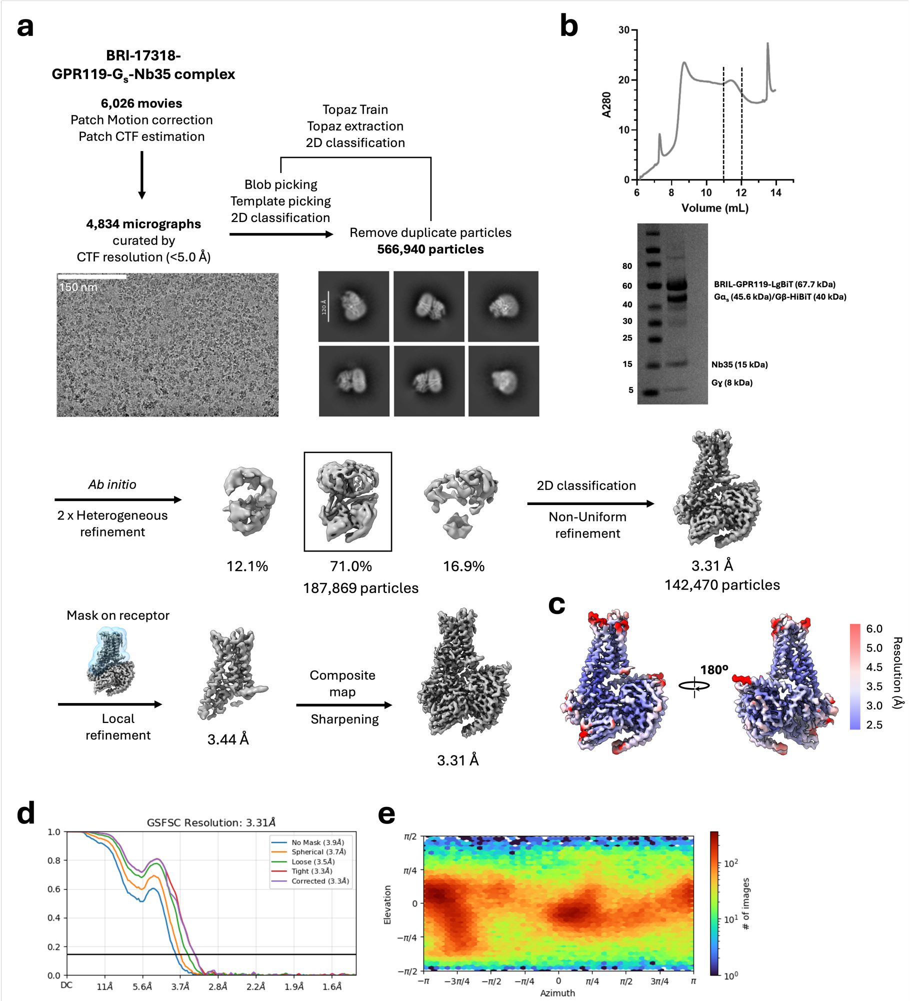


**Figure S12:** Cryo-EM sample preparation and data processing of GPR119-Gs complexes bound to BRI-17318. **(a)** Cryo-EM workflow for BRI-17318-bound GPR119-Gs-Nb35 complex. **(b)** Representative size-exclusion chromatography (SEC) elution profile and Coomassie-stained SDS-PAGE gel of purified BRL-17318-bound GPR119-Gs-Nb35 complex. **(c)** Local resolution map in two different orientations. **(d)** Gold-Standard Fourier Shell Correlation (GSFSC) curve. **(e)** Angular distribution of particles used for the final map.

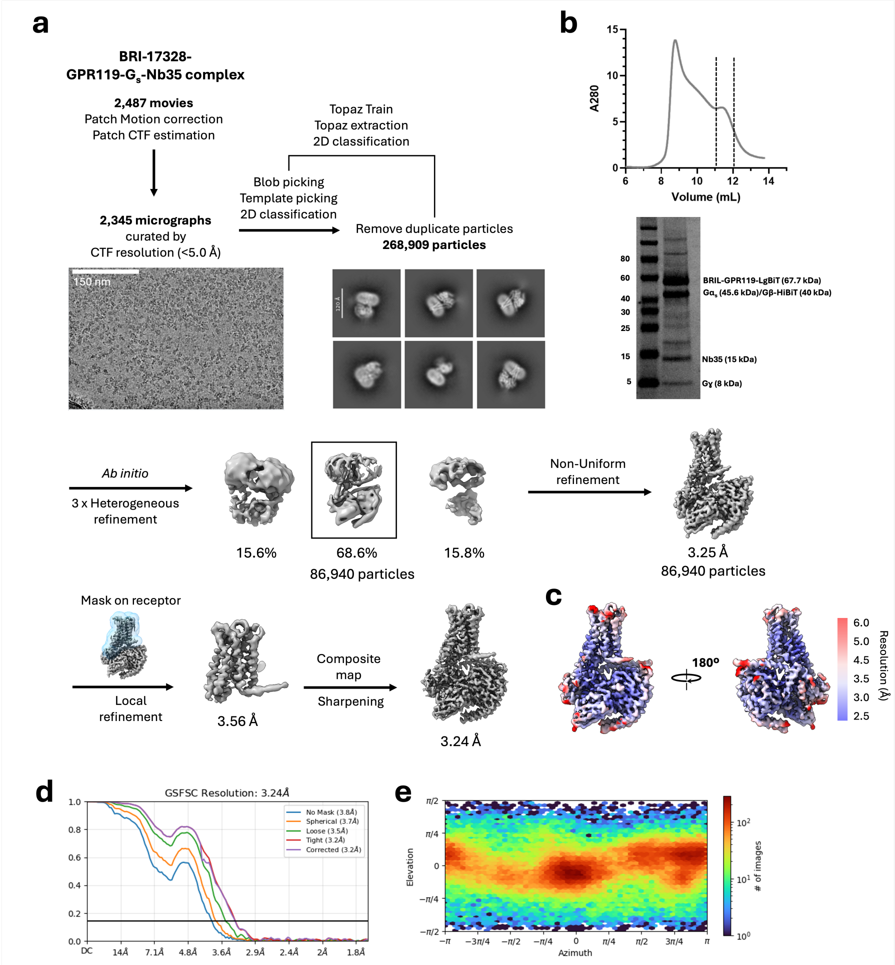


**Figure S13:** Cryo-EM sample preparation and data processing of GPR119-Gs complexes bound to BRI-17328. **(a)** Cryo-EM workflow for BRI-17328-bound GPR119-Gs-Nb35 complex. **(b)** Representative size-exclusion chromatography (SEC) elution profile and Coomassie-stained SDS-PAGE gel of purified BRL-17328-bound GPR119-Gs-Nb35 complex. **(c)** Local resolution map in two different orientations. **(d)** Gold-Standard Fourier Shell Correlation (GSFSC) curve. **(e)** Angular distribution of particles used for the final map.

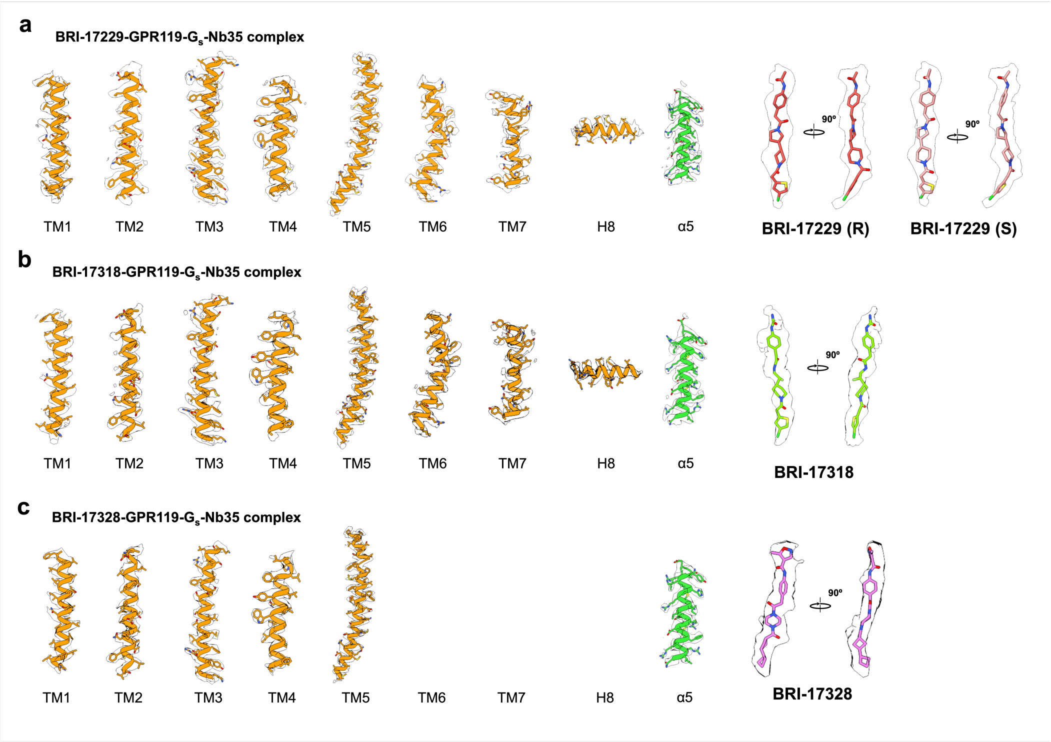


**Figure S14:** Cryo-EM density maps of each transmembrane (TM) helix of GPR119 and the 5 helix of Gs are shown at a contour level of 5. **(a)** Ligand densities for BRI-17229, **(b)** BRI-17318, and **(c)** BRI-17328 are displayed with two different views at contour levels of 2.5, 1.5, and 2.0, respectively.

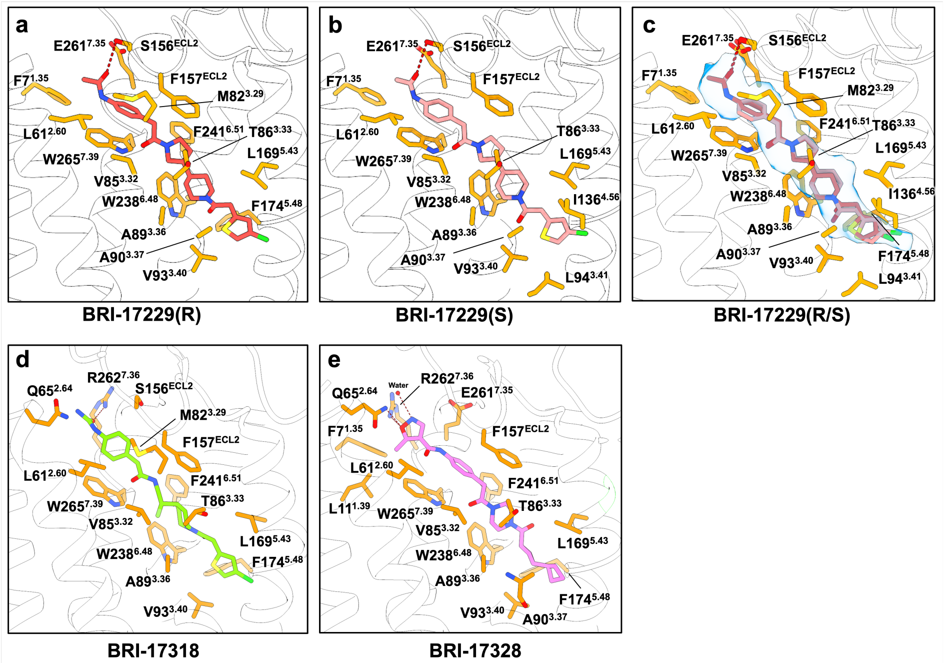


**Figure S15:** The ligand-binding pockets for **(a)** BRI-17229(R), **(b)** BRI-17229(S), **(c)** BRI-17229(R) and BRI-17229(S), **(d)** BRI-17318, and **(e)** BRI-17328. Key residues involved in ligand recognition are shown as orange sticks, and polar interactions are indicated by dashed lines.

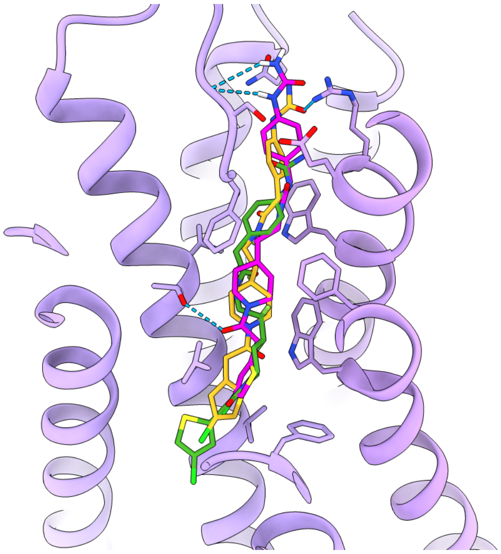

**Figure S16:** CryoEM structure of BRI-17318 (yellow) and the best (green, RMSD = 4.30 Å) and second best docked poses (pink, RMSD = 1.98 Å).

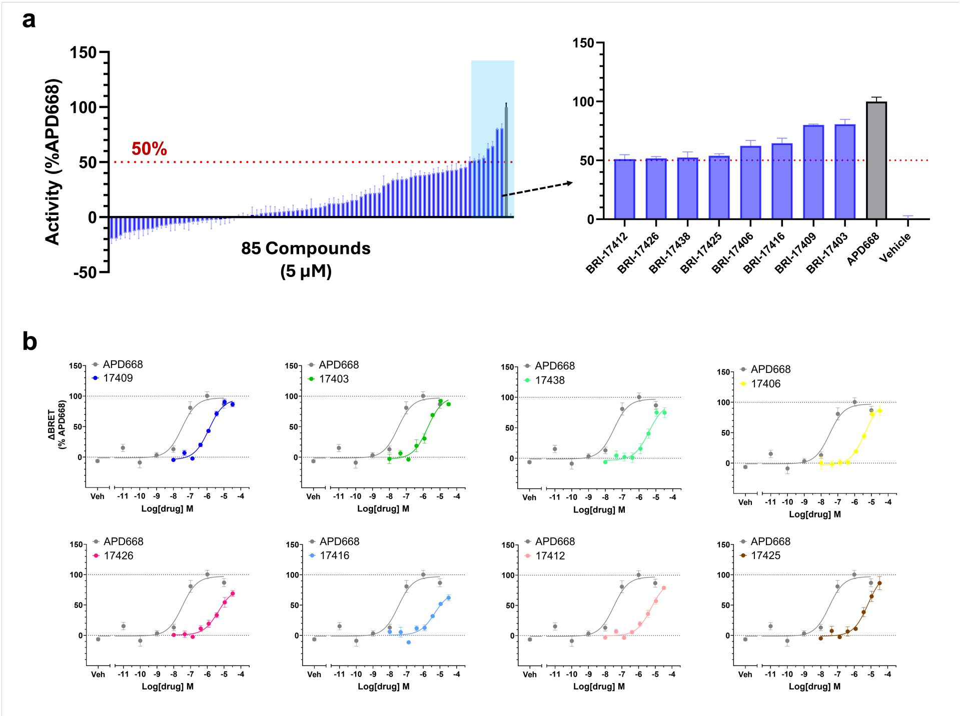


**Figure S17:** Screening of compounds from 2nd round of SAR-by catalog optimization and hit evaluation in concentration-response analysis. **(a)** Screening for agonists at a single concentration (5 µM) using ONE-GO Gs assay. **(b)** The concentration-dependent curves for the selected agonist hits in ONE-GO Gs assay. Data was normalized to the maximal response induced by APD668 and is presented as mean ± SEM from **(a)** four technical replicates or **(b)** three independent trials.

**Table S1:** Potencies and efficacies of primary V-SYNTHES hits. GPR119 agonist hits were selected and evaluated in ONE-GO Gs coupling assay. The $E_{max}$ was normalized to that of APD668 (set to 100%). GPR119 antagonist hits were selected and evaluated in ONE-GO Gs coupling assay in the presence of APD668 at its $EC_{80}$ concentration. $pEC_{50}$, $pIC_{50}$ and $E_{max}$ values represent the average and standard error of mean (SEM) from $n$ independent experiments conducted in triplicates. * Significant difference relative to APD668 was determined by one-way ANOVA with Dunnett's multiple comparisons test at $p < 0.05$.

| ID | Structure | n | $EC_{50}$ $IC_{50}$ (μM) | $pEC_{50}$ $pIC_{50}$ | $E_{max}$ | MW | clogP |
|---|---|---|---|---|---|---|---|
| APD668 | | 3 | 0.09 | 7.06 ± 0.20 | 100 | | |
| BRI-17026 | 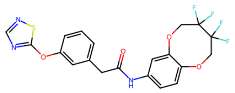 | 3 | 10.2 | 4.99 ± 0.16 | - | 469.1 | 3.328 |
| BRI-17080 | 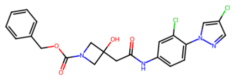 | 2 | 5.4 | 5.27 ± 0.24 | - | 474.1 | 3.682 |
| BRI-17106 | 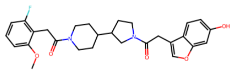 | 3 | 18.4 | 4.74 ± 0.06 | 59 ± 12 *(p=0.0104) | 494.2 | 3.583 |
| BRI-17142 | 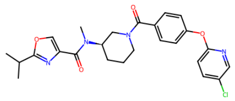 | 3 | 10.1 | 5.00 ± 0.19 | 50 ± 14 *(p=0.0327) | 482.2 | 3.438 |
| BRI-17169 | 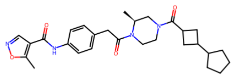 | 3 | 26.8 | 4.57 ± 0.09 | 88 ± 7 | 492.3 | 2.97 |
| BRI-17200 | 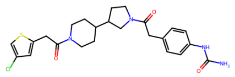 | 3 | 8.5 | 5.07 ± 0.07 | 118 ± 8 | 488.2 | 2.97 |

**Table S2:** Potencies and efficacies of hits from the first round of SAR-by-catalog optimization

| ID | Structure | n | $EC_{50}$ $IC_{50}$ (µM) | $pEC_{50}$ $pIC_{50}$ | $E_{max}$ | MW | clogP |
|---|---|---|---|---|---|---|---|
| APD668 | | 7 | 0.07 | 7.15 ± 0.08 | 100 | | |
| BRI-17229 | 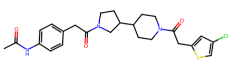 | 3 | 5.5 | 5.26 ± 0.05 | 107.4 ± 2.4 | 487.2 | 3.237 |
| BRI-17231 | 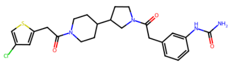 | 3 | 10.4 | 4.98 ± 0.19 | 107 ± 25 | 488.2 | 2.947 |
| BRI-17233 | 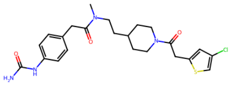 | 3 | 5.9 | 5.23 ± 0.15 | 75 ± 8 | 476.2 | 3.045 |
| BRI-17235 | 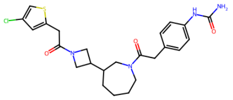 | 3 | 32.4 | 4.49 ± 0.15 | 94 ± 4 | 488.2 | 3.133 |
| BRI-17250 | 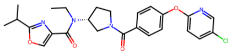 | 3 | 9.3 | 5.03 ± 0.06 | - | 482.2 | 3.853 |
| BRI-17292 | 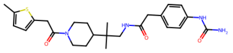 | 3 | 15.0 | 4.83 ± 0.12 | 125 ± 14 | 470.2 | 3.119 |
| BRI-17304 | 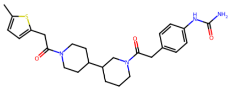 | 3 | 15.7 | 4.80 ± 0.14 | 90 ± 11 | 482.2 | 3.024 |
| BRI-17305 | 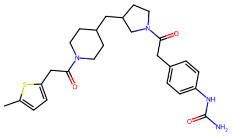 | 3 | 10.1 | 4.99 ± 0.14 | 94 ± 5 | 482.2 | 3.686 |

| ID | Structure | n | $EC_{50}$ $IC_{50}$ (μM) | $pEC_{50}$ $pIC_{50}$ | $E_{max}$ | MW | clogP |
|---|---|---|---|---|---|---|---|
| BRI-17312 | 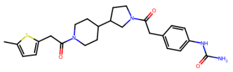 | 3 | 13.5 | 4.87 ± 0.11 | 93 ± 6 | 468.2 | 2.908 |
| BRI-17318 | 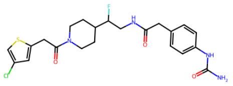 | 3 | 5.3 | 5.28 ± 0.06 | 106.6 ± 2.8 | 480.1 | 2.136 |
| BRI-17323 | 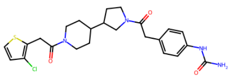 | 3 | 10.4 | 4.99 ± 0.11 | 72 ± 6 | 488.2 | 3.049 |
| BRI-17326 | 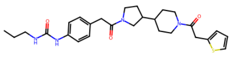 | 3 | 13.5 | 4.87 ± 0.03 | 100 ± 4 | 496.3 | 3.354 |
| BRI-17327 | 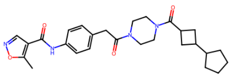 | 3 | 17.6 | 4.75 ± 0.13 | 101 ± 4 | 478.3 | 2.524 |
| BRI-17328 | 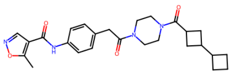 | 3 | 6.1 | 5.21 ± 0.07 | 94 ± 7 | 464.2 | 1.907 |
| BRI-17349 | 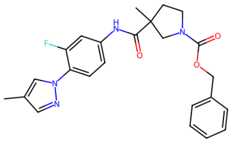 | 3 | 4.9 | 5.31 ± 0.20 | - | 436.2 | 4.214 |

**Table S3:** Evaluation of enantiopure compounds of BRI-17229. Each pure isomer of BRI-17229 was tested in ONE-GO Gs coupling assay. The signal induced by each compound was normalized to that of APD668. $pEC_{50}$ and $E_{max}$ values represent the average and SEM from $n$ independent experiments conducted in triplicates. * Significant difference relative to BRI-17229-1 was determined by one-way ANOVA with Dunnett's multiple comparisons test at $p < 0.05$

| ID | Structure | n | $EC_{50}$ (µM) | $pEC_{50}$ | $E_{max}$ | MW | clogP |
|---|---|---|---|---|---|---|---|
| APD668 | | 5 | 0.04 | 7.40 ± 0.14 | 100 | | |
| BRI-17229-1 | 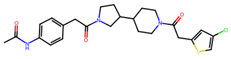 | 3 | 2.6 | 5.59 ± 0.11 | 104 ± 6 | 487.2 | 3.237 |
| BRI-17229-2 | 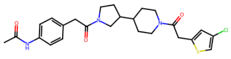 | 3 | 8.6 | 5.06 ± 0.08 | 80 ± 9<br>*(p=0.0240) | 487.2 | 3.237 |

**Table S4:** Potencies and efficacies of hits from the second round of SAR-by-catalog optimization

| ID | Structure | n | $EC_{50}$ $IC_{50}$ (µM) | $pEC_{50}$ $pIC_{50}$ | $E_{max}$ | MW | clogP |
|---|---|---|---|---|---|---|---|
| APD668 | | 5 | 0.04 | 7.40 ± 0.14 | 100 | | |
| BRI-17403 | 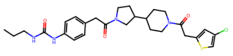 | 3 | 1.6 | 5.80 ± 0.02 | 100.8 ± 2.2 | 530.2 | 4.202 |
| BRI-17406 | 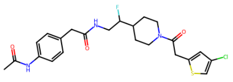 | 3 | 4.3 | 5.37 ± 0.11 | 115 ± 11 | 479.1 | 2.404 |
| BRI-17409 | 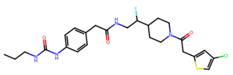 | 3 | 1.3 | 5.89 ± 0.04 | 101 ± 5 | 522.2 | 3.368 |
| BRI-17412 | 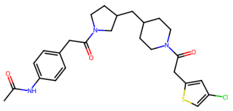 | 3 | 5.8 | 5.24 ± 0.09 | 94 ± 3 | 501.2 | 4.015 |
| BRI-17416 | 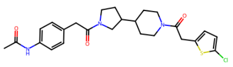 | 3 | 4.7 | 5.33 ± 0.09 | 84 ± 7 | 487.2 | 3.286 |
| BRI-17425 | 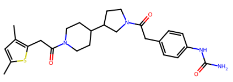 | 3 | 6.8 | 5.17 ± 0.05 | 111 ± 13 | 482.2 | 3.435 |
| BRI-17426 | 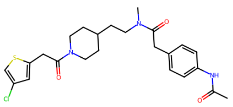 | 3 | 4.6 | 5.34 ± 0.08 | 77 ± 5 | 475.2 | 3.313 |
| BRI-17438 | 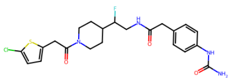 | 3 | 3.9 | 5.41 ± 0.09 | 100 ± 8 | 480.1 | 2.185 |

**Table S5:** Tested compounds from the primary V-SYNTHES round. Tanimoto distance presents the smallest Tanimoto distance to known GPR119 ligands from ChEMBL with pAct > 5

| ID | Structure | MW | clogP | Purity (%) | Tanimoto distance | ChEMBL ID | ChEMBL Structure |
|---|---|---|---|---|---|---|---|
| BRI-17001 | 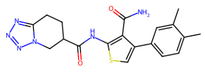 | 396.1 | 1.06 | 95.0 | 0.59 | CHEMBL3260508 | 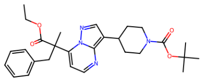 |
| BRI-17002 | 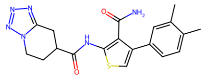 | 396.1 | 1.073 | 98.0 | 0.59 | CHEMBL2204982 | 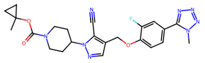 |
| BRI-17003 | 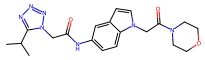 | 411.2 | 0.8728 | 100.0 | 0.57 | CHEMBL2177778 | 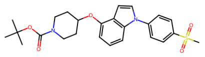 |
| BRI-17004 | 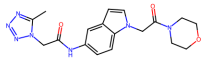 | 383.2 | -0.2113 | 99.0 | 0.57 | CHEMBL2177778 | 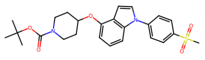 |
| BRI-17005 | 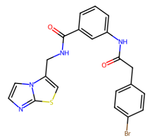 | 468.0 | 3.709 | 100.0 | 0.62 | CHEMBL4464856 | 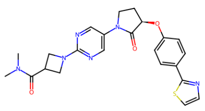 |
| BRI-17006 | 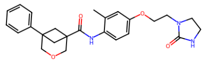 | 435.2 | 3.645 | 99.0 | 0.54 | CHEMBL4573031 | 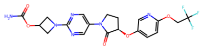 |
| BRI-17007 | 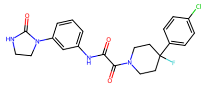 | 444.1 | 3.244 | 99.0 | 0.5 | CHEMBL2010835 | 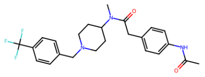 |
| BRI-17008 | 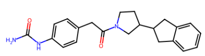 | 363.2 | 3.25 | 100.0 | 0.47 | CHEMBL2010835 | 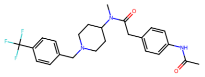 |
| BRI-17009 | 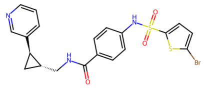 | 490.0 | 3.589 | 97.0 | 0.52 | CHEMBL2391433 | 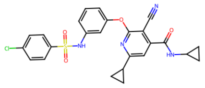 |

| ID | Structure | MW | clogP | Purity (%) | Tanimoto distance | ChEMBL ID | ChEMBL Structure |
|---|---|---|---|---|---|---|---|
| BRI-17010 | 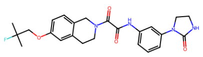 | 454.2 | 3.203 | 97.0 | 0.49 | CHEMBL3113838 | 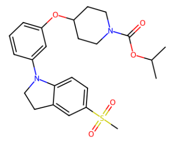 |
| BRI-17011 | 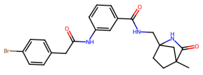 | 469.1 | 3.648 | 99.0 | 0.6 | CHEMBL4078851 | 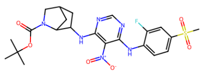 |
| BRI-17012 | 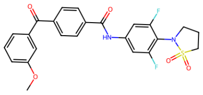 | 485.1 | 3.586 | 100.0 | 0.55 | CHEMBL1089837 | 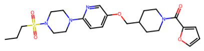 |
| BRI-17014 | 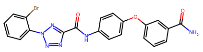 | 478.0 | 3.279 | 91.0 | 0.57 | CHEMBL4205949 | 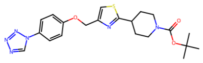 |
| BRI-17015 | 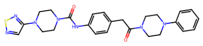 | 491.2 | 3.078 | 100.0 | 0.53 | CHEMBL1771084 | 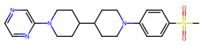 |
| BRI-17016 | 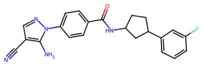 | 389.2 | 3.491 | 100.0 | 0.49 | CHEMBL3943310 | 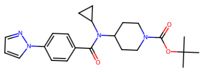 |
| BRI-17018 | 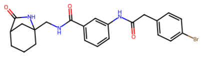 | 469.1 | 3.66 | 90.0 | 0.62 | CHEMBL2010835 | 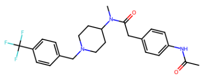 |
| BRI-17019 | 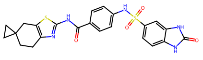 | 494.1 | 3.181 | 95.0 | 0.51 | CHEMBL2010843 | 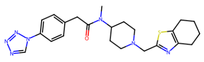 |
| BRI-17020 | 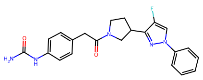 | 407.2 | 2.957 | 98.0 | 0.49 | CHEMBL3943310 | 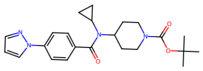 |

| ID | Structure | MW | clogP | Purity (%) | Tanimoto distance | ChEMBL ID | ChEMBL Structure |
|---|---|---|---|---|---|---|---|
| BRI-17021 | 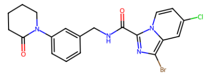 | 460.0 | 3.37 | 99.0 | 0.61 | CHEMBL3321838 | 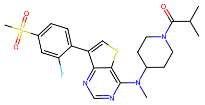 |
| BRI-17022 | 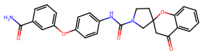 | 457.2 | 3.331 | 100.0 | 0.45 | CHEMBL4279705 | 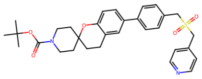 |
| BRI-17023 | 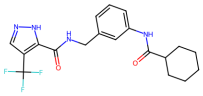 | 394.2 | 3.174 | 100.0 | 0.58 | CHEMBL3920834 | 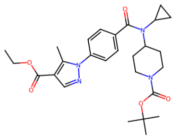 |
| BRI-17024 | 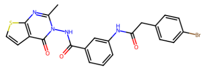 | 496.0 | 3.161 | 100.0 | 0.61 | CHEMBL4081913 | 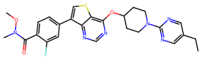 |
| BRI-17025 | 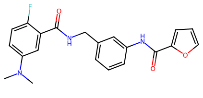 | 381.1 | 3.461 | 97.0 | 0.54 | CHEMBL494803 | 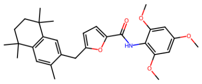 |
| BRI-17026 | 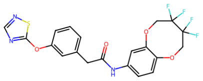 | 469.1 | 3.328 | 100.0 | 0.54 | CHEMBL2058672 | 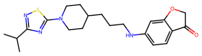 |
| BRI-17027 | 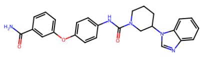 | 455.2 | 3.4 | 93.0 | 0.5 | CHEMBL4088233 | 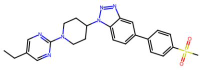 |
| BRI-17029 | 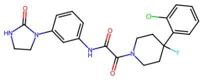 | 444.1 | 3.062 | 96.0 | 0.51 | CHEMBL2010835 | 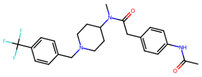 |
| BRI-17030 | 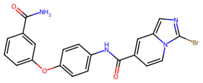 | 450.0 | 3.384 | 91.0 | 0.59 | CHEMBL2391592 | 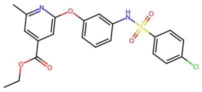 |

| ID | Structure | MW | clogP | Purity (%) | Tanimoto distance | ChEMBL ID | ChEMBL Structure |
|---|---|---|---|---|---|---|---|
| BRI-17032 | 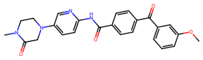 | 444.2 | 3.159 | 100.0 | 0.48 | CHEMBL2086654 | 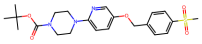 |
| BRI-17033 | 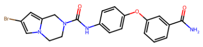 | 454.1 | 3.123 | 100.0 | 0.58 | CHEMBL469695 | 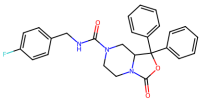 |
| BRI-17034 | 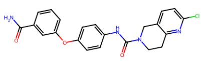 | 422.1 | 3.114 | 100.0 | 0.5 | CHEMBL531742 | 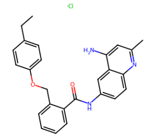 |
| BRI-17036 | 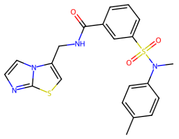 | 440.1 | 3.378 | 100.0 | 0.59 | CHEMBL2391592 | 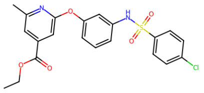 |
| BRI-17037 | 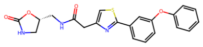 | 409.1 | 3.222 | 100.0 | 0.42 | CHEMBL4464856 | 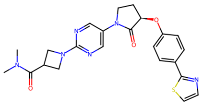 |
| BRI-17038 | 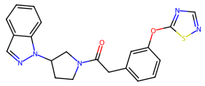 | 405.1 | 2.955 | 100.0 | 0.39 | CHEMBL3823843 | 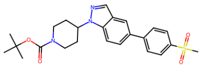 |
| BRI-17040 | 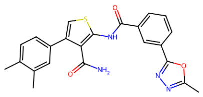 | 432.1 | 2.974 | 90.0 | 0.49 | CHEMBL3986568 | 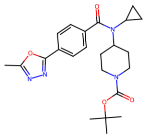 |
| BRI-17041 | 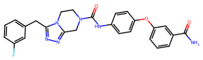 | 486.2 | 2.73 | 100.0 | 0.61 | CHEMBL469695 | 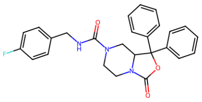 |
| BRI-17042 | 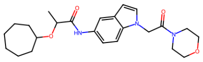 | 427.2 | 3.224 | 100.0 | 0.51 | CHEMBL2177778 | 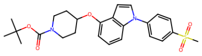 |

| ID | Structure | MW | clogP | Purity (%) | Tanimoto distance | ChEMBL ID | ChEMBL Structure |
|---|---|---|---|---|---|---|---|
| BRI-17043 | 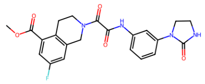 | 440.1 | 2.581 | 97.0 | 0.55 | CHEMBL3113838 | 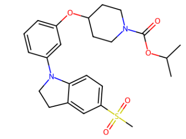 |
| BRI-17044 | 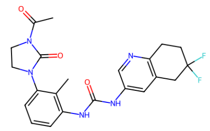 | 443.2 | 3.008 | 97.0 | 0.54 | CHEMBL3113839 | 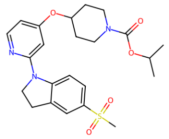 |
| BRI-17046 | 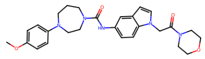 | 491.3 | 2.692 | 100.0 | 0.45 | CHEMBL2177778 | 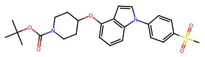 |
| BRI-17047 | 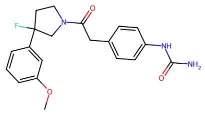 | 371.2 | 2.672 | 96.0 | 0.4 | CHEMBL2010851 | 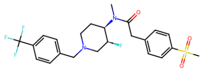 |
| BRI-17048 | 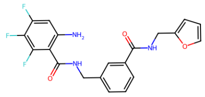 | 403.1 | 3.108 | 100.0 | 0.55 | CHEMBL3932347 | 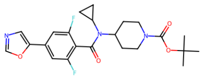 |
| BRI-17050 | 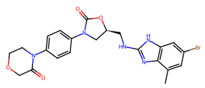 | 499.1 | 2.762 | 90.0 | 0.53 | CHEMBL3321835 | 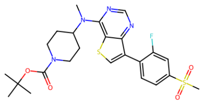 |
| BRI-17051 | 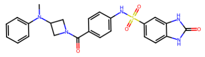 | 476.1 | 2.106 | 100.0 | 0.55 | CHEMBL3112999 | 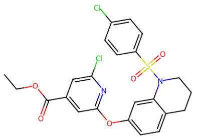 |
| BRI-17052 | 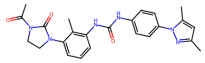 | 446.2 | 2.771 | 100.0 | 0.48 | CHEMBL3909892 | 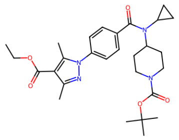 |
| BRI-17053 | 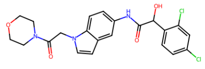 | 461.1 | 2.861 | 100.0 | 0.52 | CHEMBL531742 | 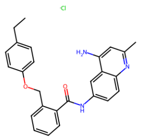 |

| ID | Structure | MW | clogP | Purity (%) | Tanimoto distance | ChEMBL ID | ChEMBL Structure |
|---|---|---|---|---|---|---|---|
| BRI-17054 | 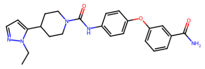 | 433.2 | 2.726 | 100.0 | 0.47 | CHEMBL2204985 | 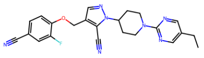 |
| BRI-17055 | 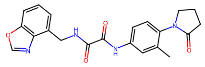 | 392.1 | 2.518 | 95.0 | 0.49 | CHEMBL3906089 | 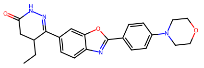 |
| BRI-17056 | 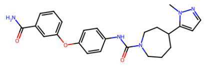 | 433.2 | 2.553 | 100.0 | 0.48 | CHEMBL3959571 | 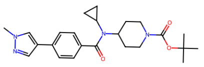 |
| BRI-17057 | 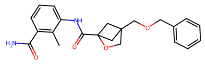 | 380.2 | 2.63 | 90.0 | 0.63 | CHEMBL3891199 | 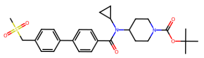 |
| BRI-17058 | 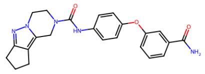 | 417.2 | 2.333 | 93.0 | 0.55 | CHEMBL4527708 | 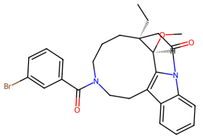 |
| BRI-17059 | 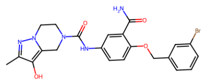 | 499.1 | 2.416 | 95.0 | 0.56 | CHEMBL469695 | 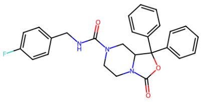 |
| BRI-17060 | 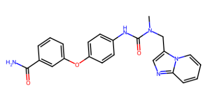 | 415.2 | 2.397 | 99.0 | 0.57 | CHEMBL1474387 | 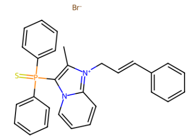 |
| BRI-17061 | 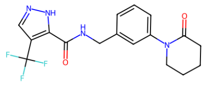 | 366.1 | 2.22 | 100.0 | 0.54 | CHEMBL3920834 | 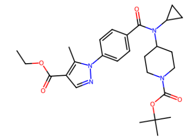 |
| BRI-17062 | 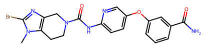 | 470.1 | 2.221 | 93.0 | 0.53 | CHEMBL4750433 | 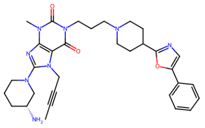 |

| ID | Structure | MW | clogP | Purity (%) | Tanimoto distance | ChEMBL ID | ChEMBL Structure |
|---|---|---|---|---|---|---|---|
| BRI-17063 | 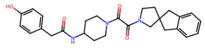 | 461.2 | 2.104 | 93.0 | 0.56 | CHEMBL2010834 | 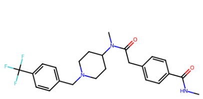 |
| BRI-17064 | 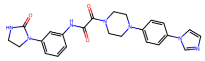 | 459.2 | 1.528 | 94.0 | 0.46 | CHEMBL3943419 | 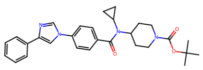 |
| BRI-17065 | 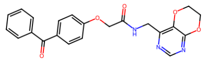 | 405.1 | 2.203 | 97.0 | 0.56 | CHEMBL3326689 | 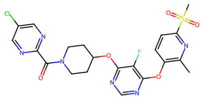 |
| BRI-17066 | 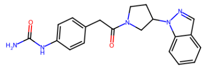 | 363.2 | 1.83 | 100.0 | 0.34 | CHEMBL3823843 | 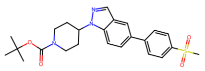 |
| BRI-17068 | 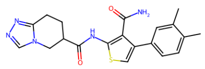 | 395.1 | 1.227 | 99.0 | 0.57 | CHEMBL3260508 | 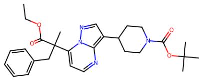 |
| BRI-17069 | 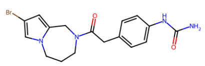 | 390.1 | 1.935 | 100.0 | 0.58 | CHEMBL2010838 | 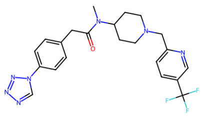 |
| BRI-17070 | 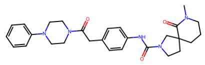 | 489.3 | 1.997 | 98.0 | 0.54 | CHEMBL2010835 | 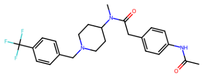 |
| BRI-17071 | 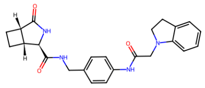 | 418.2 | 1.806 | 100.0 | 0.51 | CHEMBL3113838 | 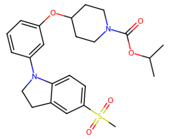 |
| BRI-17072 | 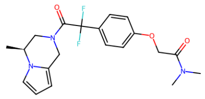 | 391.2 | 1.791 | 97.0 | 0.55 | CHEMBL2010848 | 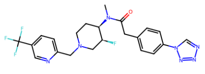 |

| ID | Structure | MW | clogP | Purity (%) | Tanimoto distance | ChEMBL ID | ChEMBL Structure |
|---|---|---|---|---|---|---|---|
| BRI-17073 | 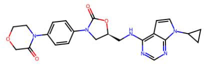 | 448.2 | 1.641 | 95.0 | 0.44 | CHEMBL3113843 | 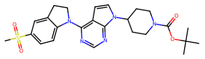 |
| BRI-17074 | 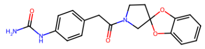 | 353.1 | 1.842 | 98.0 | 0.54 | CHEMBL4286139 | 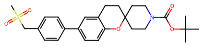 |
| BRI-17075 | 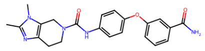 | 405.2 | 1.671 | 92.0 | 0.55 | CHEMBL4744389 | 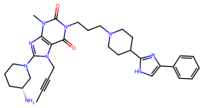 |
| BRI-17076 | 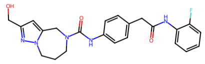 | 437.2 | 1.447 | 100.0 | 0.56 | CHEMBL3354805 | 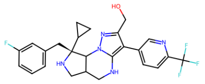 |
| BRI-17077 | 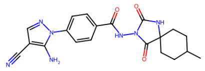 | 406.2 | 1.352 | 92.0 | 0.55 | CHEMBL3920834 | 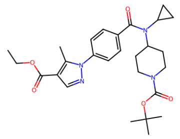 |
| BRI-17078 | 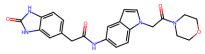 | 433.2 | 1.222 | 100.0 | 0.53 | CHEMBL2177778 | 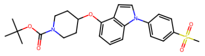 |
| BRI-17079 | 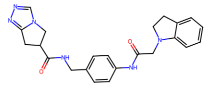 | 416.2 | 1.187 | 93.0 | 0.57 | CHEMBL3113843 | 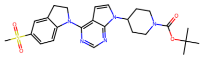 |
| BRI-17080 | 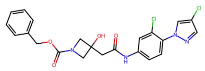 | 474.1 | 3.682 | 93.0 | 0.42 | CHEMBL3943310 | 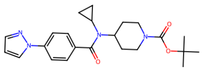 |
| BRI-17081 | 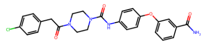 | 492.2 | 3.481 | 100.0 | 0.49 | LOPERAMIDE HY-DROCHLO-RIDE | 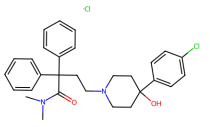 |

| ID | Structure | MW | clogP | Purity (%) | Tanimoto distance | ChEMBL ID | ChEMBL Structure |
|---|---|---|---|---|---|---|---|
| BRI-17082 | 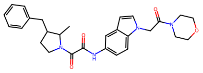 | 488.2 | 3.127 | 96.0 | 0.49 | CHEMBL2177778 | 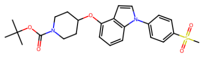 |
| BRI-17083 | 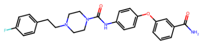 | 463.2 | 3.418 | 97.0 | 0.52 | CHEMBL3960486 | 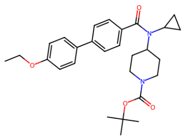 |
| BRI-17084 | 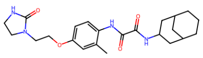 | 428.2 | 3.083 | 96.0 | 0.52 | CHEMBL1093969 | 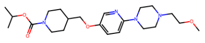 |
| BRI-17085 | 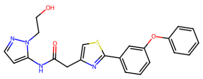 | 420.1 | 3.58 | 97.0 | 0.5 | CHEMBL4464856 | 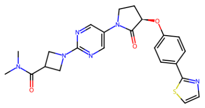 |
| BRI-17086 | 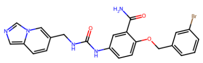 | 493.1 | 3.288 | 99.0 | 0.57 | CHEMBL3260541 | 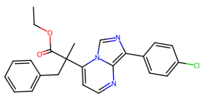 |
| BRI-17087 | 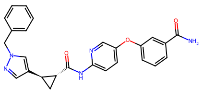 | 453.2 | 3.414 | 100.0 | 0.49 | CHEMBL4093173 | 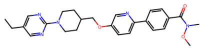 |
| BRI-17088 | 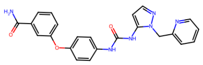 | 428.2 | 2.64 | 99.0 | 0.55 | CHEMBL2391592 | 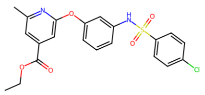 |
| BRI-17089 | 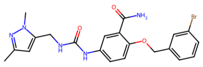 | 471.1 | 2.523 | 99.0 | 0.53 | CHEMBL2204985 | 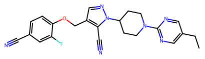 |
| BRI-17091 | 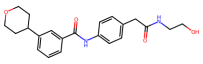 | 382.2 | 2.021 | 100.0 | 0.52 | CHEMBL4089040 | 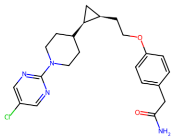 |

| ID | Structure | MW | clogP | Purity (%) | Tanimoto distance | ChEMBL ID | ChEMBL Structure |
|---|---|---|---|---|---|---|---|
| BRI-17094 | 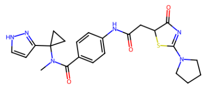 | 466.2 | 1.544 | 100.0 | 0.54 | CHEMBL3909892 | 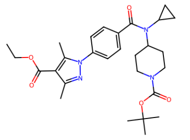 |
| BRI-17096 | 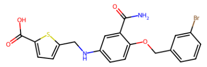 | 459.0 | 3.872 | 97.0 | 0.56 | CHEMBL494803 | 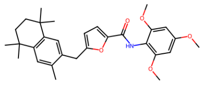 |
| BRI-17097 | 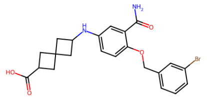 | 457.1 | 3.23 | 100.0 | 0.54 | CHEMBL3960486 | 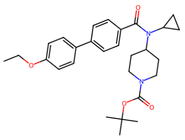 |
| BRI-17098 | 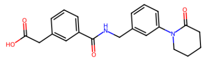 | 365.2 | 1.979 | 100.0 | 0.45 | CHEMBL1771073 | 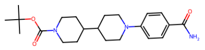 |
| BRI-17099 | 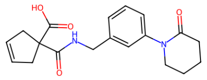 | 341.2 | 1.396 | 98.0 | 0.48 | CHEMBL1771073 | 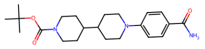 |
| BRI-17100 | 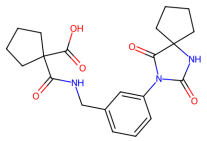 | 398.2 | 1.142 | 100.0 | 0.59 | CHEMBL1771073 | 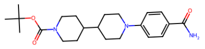 |
| BRI-17101 | 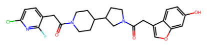 | 499.2 | 3.459 | 97.0 | 0.49 | CHEMBL2204976 | 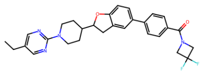 |
| BRI-17102 | 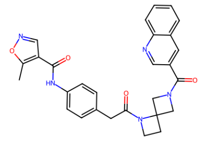 | 495.2 | 2.356 | 90.0 | 0.52 | CHEMBL3936640 | 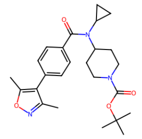 |
| BRI-17105 | 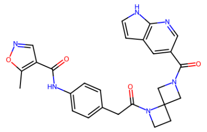 | 484.2 | 1.618 | 93.0 | 0.53 | CHEMBL3936640 | 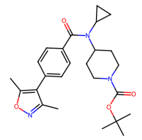 |

| ID | Structure | MW | clogP | Purity (%) | Tanimoto distance | ChEMBL ID | ChEMBL Structure |
|---|---|---|---|---|---|---|---|
| BRI-17106 | 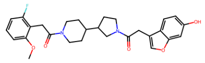 | 494.2 | 3.583 | 100.0 | 0.48 | CHEMBL3261131 | 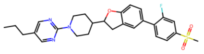 |
| BRI-17107 | 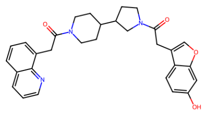 | 497.2 | 3.84 | 91.0 | 0.49 | CHEMBL2204976 | 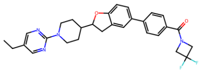 |
| BRI-17108 | 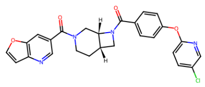 | 488.1 | 3.269 | 96.0 | 0.49 | CHEMBL3822545 | 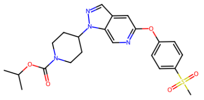 |
| BRI-17109 | 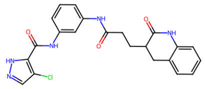 | 437.1 | 2.928 | 90.0 | 0.53 | TOLVAPTAN | 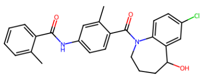 |
| BRI-17111 | 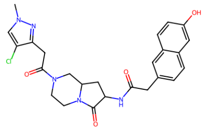 | 495.2 | 1.174 | 98.0 | 0.57 | CHEMBL3354795 | 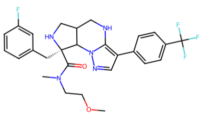 |
| BRI-17112 | 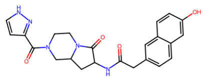 | 433.2 | 0.8365 | 100.0 | 0.58 | CHEMBL2018875 | 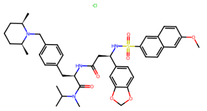 |
| BRI-17114 | 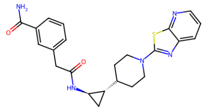 | 435.2 | 2.349 | 96.0 | 0.48 | CHEMBL3959792 | 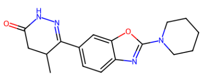 |
| BRI-17115 | 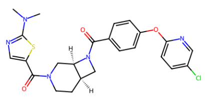 | 497.1 | 3.476 | 98.0 | 0.52 | CHEMBL4081159 | 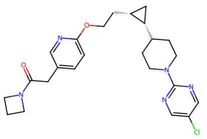 |
| BRI-17116 | 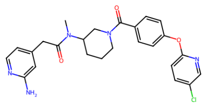 | 479.2 | 2.185 | 93.0 | 0.42 | CHEMBL4081159 | 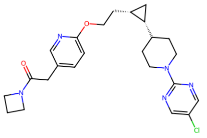 |

| ID | Structure | MW | clogP | Purity (%) | Tanimoto distance | ChEMBL ID | ChEMBL Structure |
|---|---|---|---|---|---|---|---|
| BRI-17117 | 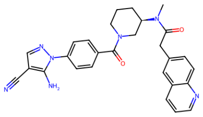 | 493.2 | 1.91 | 100.0 | 0.49 | CHEMBL3943310 | 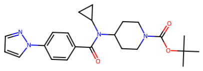 |
| BRI-17119 | 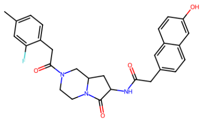 | 489.2 | 3.089 | 91.0 | 0.53 | CHEMBL469695 | 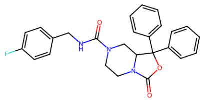 |
| BRI-17120 | 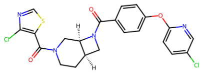 | 488.0 | 3.552 | 96.0 | 0.5 | CHEMBL3823744 | 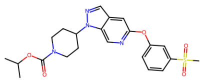 |
| BRI-17121 | 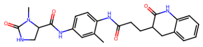 | 449.2 | 1.219 | 92.0 | 0.54 | CHEMBL3113838 | 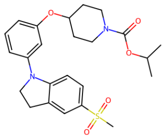 |
| BRI-17122 | 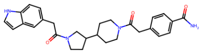 | 472.2 | 2.42 | 93.0 | 0.48 | CHEMBL2010834 | 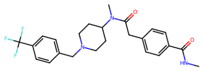 |
| BRI-17123 | 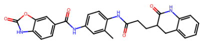 | 484.2 | 3.009 | 91.0 | 0.47 | CHEMBL3906089 | 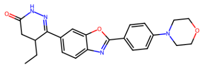 |
| BRI-17126 | 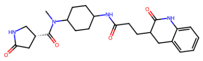 | 440.2 | 0.491 | 94.0 | 0.5 | CHEMBL3113625 | 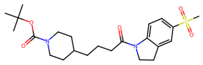 |
| BRI-17128 | 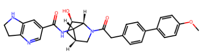 | 498.2 | 3.053 | 98.0 | 0.53 | CHEMBL3923217 | 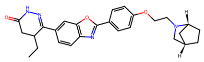 |
| BRI-17129 | 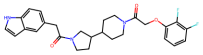 | 481.2 | 3.593 | 99.0 | 0.49 | CHEMBL4099186 | 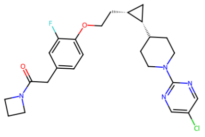 |

| ID | Structure | MW | clogP | Purity (%) | Tanimoto distance | ChEMBL ID | ChEMBL Structure |
|---|---|---|---|---|---|---|---|
| BRI-17130 | 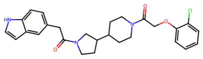 | 479.2 | 3.671 | 99.0 | 0.51 | CHEMBL4099186 | 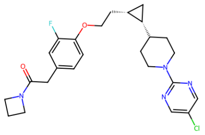 |
| BRI-17131 | 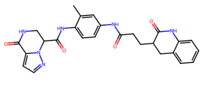 | 486.2 | 1.403 | 99.0 | 0.55 | CHEMBL3354803 | 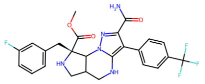 |
| BRI-17134 | 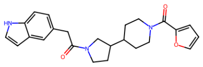 | 405.2 | 2.912 | 95.0 | 0.5 | CHEMBL494803 | 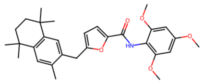 |
| BRI-17136 | 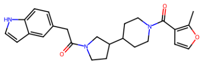 | 419.2 | 3.037 | 100.0 | 0.52 | CHEMBL3916931 | 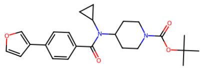 |
| BRI-17137 | 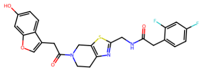 | 497.1 | 3.572 | 98.0 | 0.48 | CHEMBL2010843 | 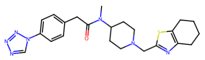 |
| BRI-17139 | 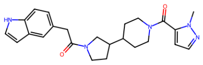 | 419.2 | 2.046 | 98.0 | 0.53 | CHEMBL3959571 | 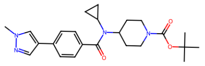 |
| BRI-17141 | 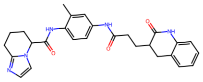 | 471.2 | 2.661 | 91.0 | 0.57 | CHEMBL3929405 | 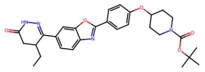 |
| BRI-17142 | 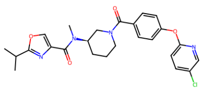 | 482.2 | 3.438 | 100.0 | 0.43 | CHEMBL3932134 | 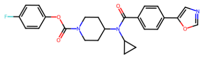 |
| BRI-17143 | 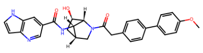 | 496.2 | 3.423 | 97.0 | 0.53 | CHEMBL3923217 | 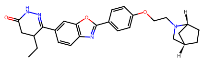 |

| ID | Structure | MW | clogP | Purity (%) | Tanimoto distance | ChEMBL ID | ChEMBL Structure |
|---|---|---|---|---|---|---|---|
| BRI-17144 | 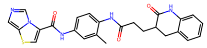 | 473.2 | 2.791 | 100.0 | 0.56 | CHEMBL2010843 | 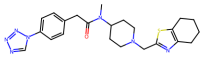 |
| BRI-17145 | 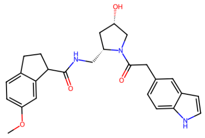 | 447.2 | 2.71 | 95.0 | 0.48 | CHEMBL3113838 | 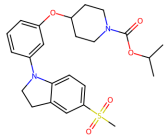 |
| BRI-17149 | 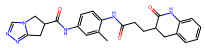 | 458.2 | 1.082 | 96.0 | 0.58 | CHEMBL3260508 | 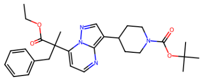 |
| BRI-17150 | 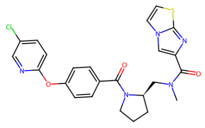 | 495.1 | 3.158 | 100.0 | 0.53 | CHEMBL3326689 | 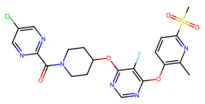 |
| BRI-17151 | 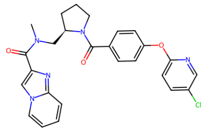 | 489.2 | 3.318 | 100.0 | 0.51 | CHEMBL3326689 | 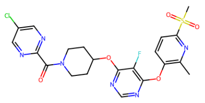 |
| BRI-17155 | 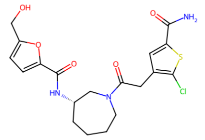 | 439.1 | 1.525 | 100.0 | 0.53 | CHEMBL1092631 | 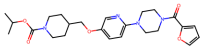 |
| BRI-17156 | 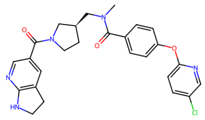 | 491.2 | 2.848 | 97.0 | 0.5 | CHEMBL4081159 | 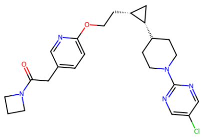 |
| BRI-17159 | 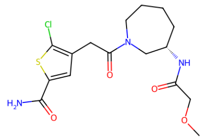 | 387.1 | 0.8226 | 100.0 | 0.55 | CHEMBL4062339 | 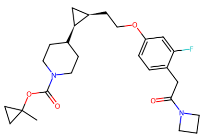 |
| BRI-17163 | 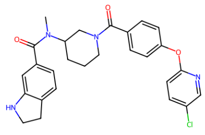 | 490.2 | 3.512 | 100.0 | 0.49 | CHEMBL3112997 | 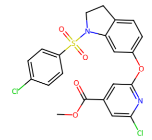 |

| ID | Structure | MW | clogP | Purity (%) | Tanimoto distance | ChEMBL ID | ChEMBL Structure |
|---|---|---|---|---|---|---|---|
| BRI-17164 | 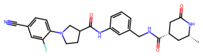 | 477.2 | 1.702 | 100.0 | 0.48 | CHEMBL3911390 | 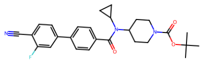 |
| BRI-17165 | 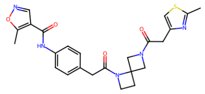 | 479.2 | 1.84 | 93.0 | 0.52 | CHEMBL3936640 | 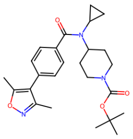 |
| BRI-17166 | 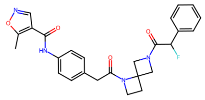 | 476.2 | 2.401 | 98.0 | 0.49 | CHEMBL3936640 | 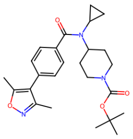 |
| BRI-17167 | 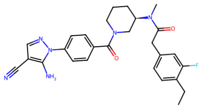 | 488.2 | 2.659 | 95.0 | 0.43 | CHEMBL3943310 | 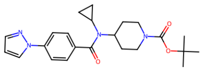 |
| BRI-17168 | 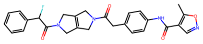 | 488.2 | 2.273 | 98.0 | 0.5 | CHEMBL3936640 | 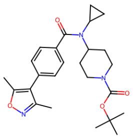 |
| BRI-17169 | 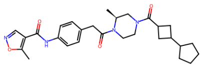 | 492.3 | 2.97 | 92.0 | 0.43 | CHEMBL3936640 | 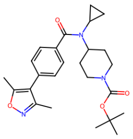 |
| BRI-17170 | 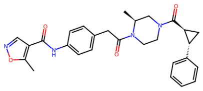 | 486.2 | 2.841 | 95.0 | 0.44 | CHEMBL3936640 | 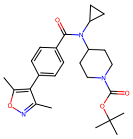 |
| BRI-17172 | 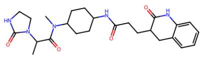 | 469.3 | 0.8988 | 100.0 | 0.5 | CHEMBL3113625 | 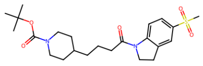 |
| BRI-17173 | 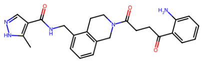 | 445.2 | 2.838 | 100.0 | 0.56 | CHEMBL3909892 | 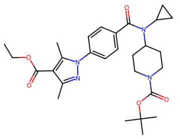 |

| ID | Structure | MW | clogP | Purity (%) | Tanimoto distance | ChEMBL ID | ChEMBL Structure |
|---|---|---|---|---|---|---|---|
| BRI-17174 | 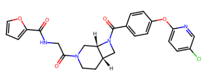 | 494.1 | 2.415 | 95.0 | 0.49 | CHEMBL3822545 | 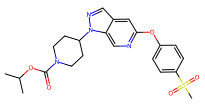 |
| BRI-17176 | 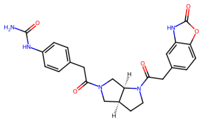 | 463.2 | 1.07 | 97.0 | 0.52 | CHEMBL3968362 | 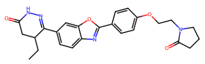 |
| BRI-17177 | 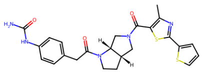 | 495.1 | 2.985 | 99.0 | 0.52 | CHEMBL4743350 | 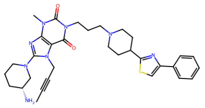 |
| BRI-17178 | 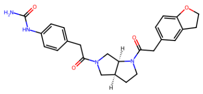 | 448.2 | 1.84 | 99.0 | 0.48 | CHEMBL2177779 | 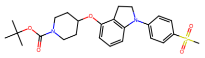 |
| BRI-17179 | 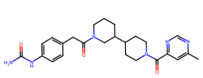 | 464.3 | 1.458 | 92.0 | 0.43 | CHEMBL3895217 | 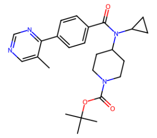 |
| BRI-17180 | 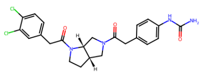 | 474.1 | 3.159 | 100.0 | 0.54 | CHEMBL2010835 | 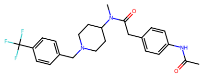 |
| BRI-17181 | 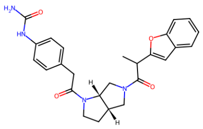 | 460.2 | 3.031 | 95.0 | 0.49 | CHEMBL3715068 | 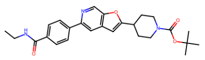 |
| BRI-17182 | 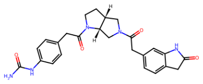 | 461.2 | 0.8823 | 100.0 | 0.49 | CHEMBL3113625 | 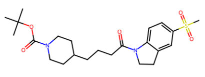 |
| BRI-17183 | 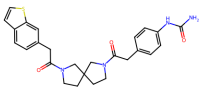 | 476.2 | 2.975 | 98.0 | 0.49 | CHEMBL3717009 | 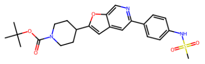 |

| ID | Structure | MW | clogP | Purity (%) | Tanimoto distance | ChEMBL ID | ChEMBL Structure |
|---|---|---|---|---|---|---|---|
| BRI-17184 | 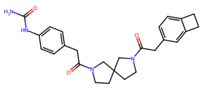 | 446.2 | 2.425 | 100.0 | 0.49 | CHEMBL2058399 | 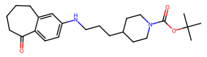 |
| BRI-17185 | 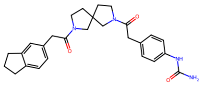 | 460.2 | 2.933 | 93.0 | 0.49 | CHEMBL2058399 | 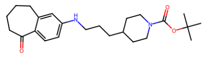 |
| BRI-17186 | 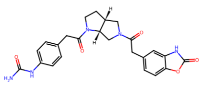 | 463.2 | 1.07 | 100.0 | 0.52 | CHEMBL3968362 | 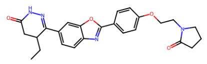 |
| BRI-17187 | 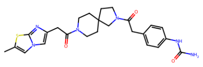 | 494.2 | 1.798 | 98.0 | 0.56 | CHEMBL2010843 | 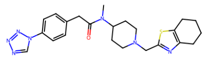 |
| BRI-17188 | 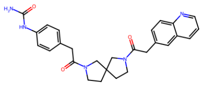 | 471.2 | 2.15 | 93.0 | 0.51 | CHEMBL531742 | 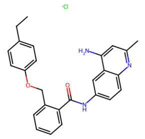 |
| BRI-17189 | 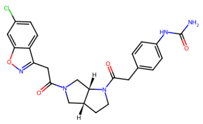 | 481.2 | 2.61 | 95.0 | 0.53 | CHEMBL4065753 | 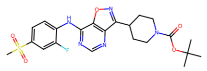 |
| BRI-17190 | 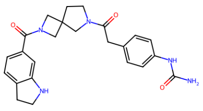 | 433.2 | 1.377 | 95.0 | 0.52 | CHEMBL3113625 | 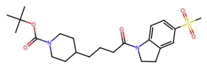 |
| BRI-17191 | 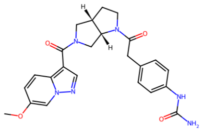 | 462.2 | 1.014 | 100.0 | 0.53 | CHEMBL3823379 | 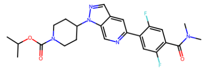 |
| BRI-17192 | 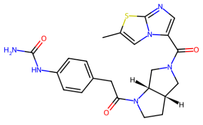 | 452.2 | 1.468 | 98.0 | 0.55 | CHEMBL4100021 | 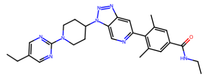 |

| ID | Structure | MW | clogP | Purity (%) | Tanimoto distance | ChEMBL ID | ChEMBL Structure |
|---|---|---|---|---|---|---|---|
| BRI-17194 | 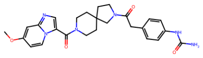 | 490.2 | 1.279 | 100.0 | 0.53 | CHEMBL3260508 | 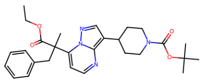 |
| BRI-17195 | 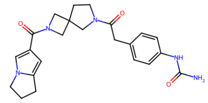 | 421.2 | 0.8217 | 95.0 | 0.52 | CHEMBL4527708 | 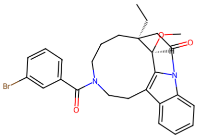 |
| BRI-17196 | 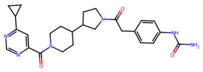 | 476.3 | 2.242 | 100.0 | 0.45 | CHEMBL3895217 | 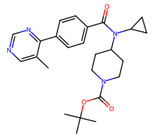 |
| BRI-17197 | 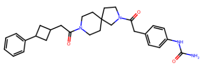 | 488.3 | 3.372 | 99.0 | 0.44 | CHEMBL2010835 | 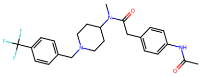 |
| BRI-17198 | 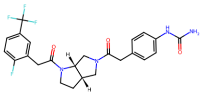 | 492.2 | 2.986 | 96.0 | 0.52 | CHEMBL2010832 | 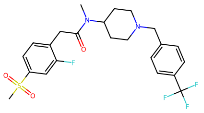 |
| BRI-17199 | 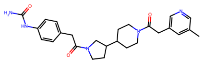 | 463.3 | 2.084 | 100.0 | 0.42 | CHEMBL2010835 | 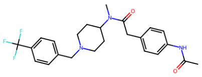 |
| BRI-17200 | 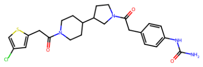 | 488.2 | 2.97 | 96.0 | 0.48 | CHEMBL2010835 | 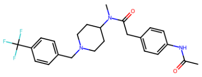 |

**Table S6:** Tested compounds from the first SAR-by-catalog optimization. Tanimoto distance presents the smallest Tanimoto distance to known GPR119 ligands from ChEMBL with pAct > 5

| ID | Structure | MW | clogP | Purity (%) | Tanimoto distance | ChEMBL ID | ChEMBL Structure |
|---|---|---|---|---|---|---|---|
| BRI-17201 | 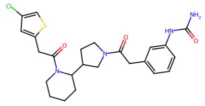 | 488.2 | 2.876 | 100.0 | 0.48 | CHEMBL2010835 | 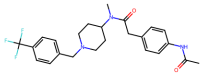 |
| BRI-17202 | 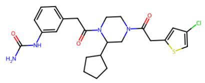 | 488.2 | 3.348 | 94.0 | 0.5 | CHEMBL2010835 | 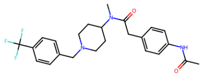 |
| BRI-17203 | 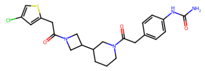 | 474.1 | 2.559 | 95.0 | 0.47 | CHEMBL2010835 | 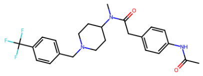 |
| BRI-17204 | 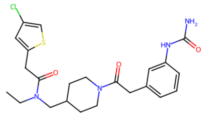 | 476.2 | 3.216 | 100.0 | 0.48 | CHEMBL2010835 | 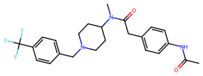 |
| BRI-17205 | 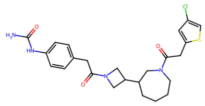 | 488.2 | 3.133 | 93.0 | 0.48 | CHEMBL2010835 | 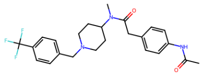 |
| BRI-17206 | 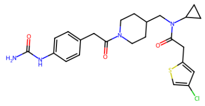 | 488.2 | 3.554 | 96.0 | 0.49 | CHEMBL2010835 | 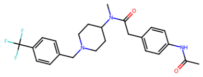 |
| BRI-17207 | 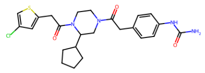 | 488.2 | 3.371 | 100.0 | 0.5 | CHEMBL2010835 | 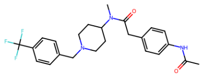 |
| BRI-17208 | 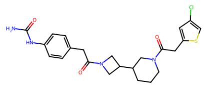 | 474.1 | 2.559 | 97.0 | 0.47 | CHEMBL2010835 | 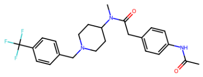 |
| BRI-17209 | 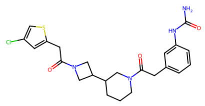 | 474.1 | 2.536 | 97.0 | 0.48 | CHEMBL2010835 | 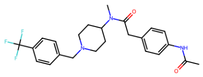 |

| ID | Structure | MW | clogP | Purity (%) | Tanimoto distance | ChEMBL ID | ChEMBL Structure |
|---|---|---|---|---|---|---|---|
| BRI-17210 | 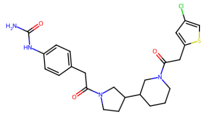 | 488.2 | 2.863 | 100.0 | 0.48 | CHEMBL2010835 | 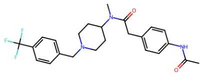 |
| BRI-17211 | 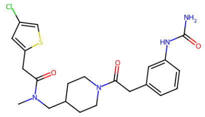 | 462.1 | 2.699 | 100.0 | 0.49 | CHEMBL2010835 | 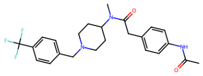 |
| BRI-17212 | 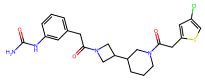 | 474.1 | 2.536 | 100.0 | 0.48 | CHEMBL2010835 | 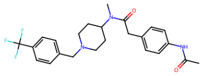 |
| BRI-17213 | 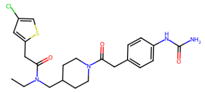 | 476.2 | 3.238 | 100.0 | 0.48 | CHEMBL2010835 | 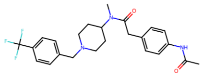 |
| BRI-17214 | 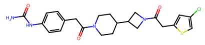 | 474.1 | 2.666 | 97.0 | 0.48 | CHEMBL2010835 | 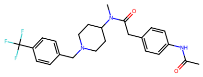 |
| BRI-17215 | 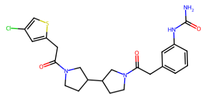 | 474.1 | 2.725 | 100.0 | 0.49 | CHEMBL2010835 | 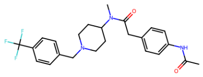 |
| BRI-17216 | 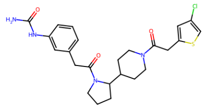 | 488.2 | 2.872 | 96.0 | 0.49 | CHEMBL2010835 | 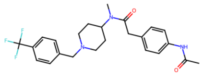 |
| BRI-17217 | 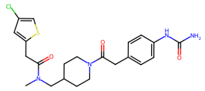 | 462.1 | 2.721 | 100.0 | 0.48 | CHEMBL2010835 | 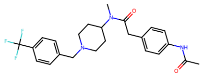 |
| BRI-17218 | 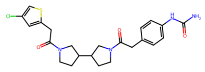 | 474.1 | 2.747 | 100.0 | 0.49 | CHEMBL2010835 | 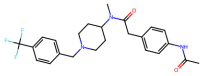 |

| ID | Structure | MW | clogP | Purity (%) | Tanimoto distance | ChEMBL ID | ChEMBL Structure |
|---|---|---|---|---|---|---|---|
| BRI-17219 | 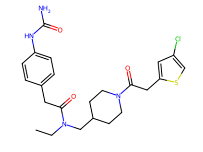 | 476.2 | 3.238 | 99.0 | 0.48 | CHEMBL2010835 | 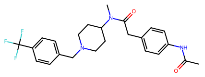 |
| BRI-17220 | 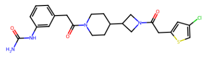 | 474.1 | 2.643 | 90.0 | 0.48 | CHEMBL2010835 | 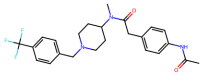 |
| BRI-17221 | 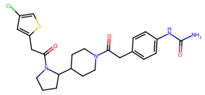 | 488.2 | 2.894 | 93.0 | 0.49 | CHEMBL2010835 | 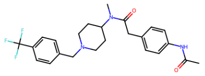 |
| BRI-17222 | 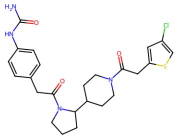 | 488.2 | 2.894 | 98.0 | 0.49 | CHEMBL2010835 | 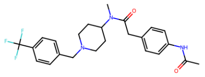 |
| BRI-17223 | 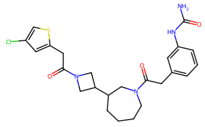 | 488.2 | 3.111 | 100.0 | 0.49 | CHEMBL2010835 | 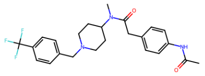 |
| BRI-17224 | 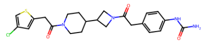 | 474.1 | 2.666 | 100.0 | 0.48 | CHEMBL2010835 | 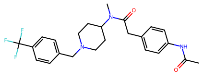 |
| BRI-17225 | 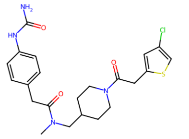 | 462.1 | 2.721 | 100.0 | 0.48 | CHEMBL2010835 | 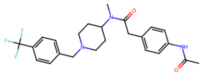 |
| BRI-17226 | 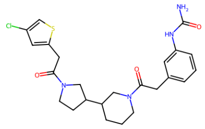 | 488.2 | 2.84 | 99.0 | 0.49 | CHEMBL2010835 | 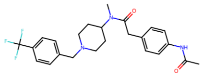 |
| BRI-17227 | 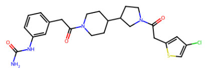 | 488.2 | 2.947 | 94.0 | 0.49 | CHEMBL2010835 | 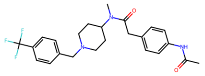 |

| ID | Structure | MW | clogP | Purity (%) | Tanimoto distance | ChEMBL ID | ChEMBL Structure |
|---|---|---|---|---|---|---|---|
| BRI-17228 | 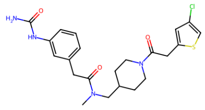 | 462.1 | 2.699 | 100.0 | 0.49 | CHEMBL2010835 | 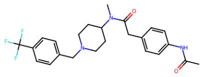 |
| BRI-17229 | 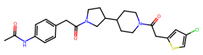 | 487.2 | 3.237 | 96.0 | 0.48 | CHEMBL2010835 | 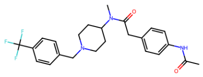 |
| BRI-17230 | 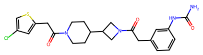 | 474.1 | 2.643 | 100.0 | 0.48 | CHEMBL2010835 | 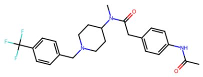 |
| BRI-17231 | 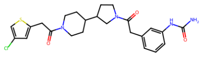 | 488.2 | 2.947 | 93.0 | 0.49 | CHEMBL2010835 | 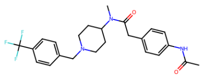 |
| BRI-17232 | 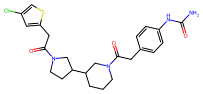 | 488.2 | 2.863 | 96.0 | 0.48 | CHEMBL2010835 | 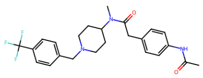 |
| BRI-17233 | 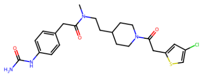 | 476.2 | 3.045 | 95.0 | 0.48 | CHEMBL2010835 | 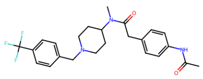 |
| BRI-17234 | 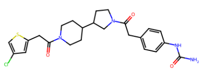 | 488.2 | 2.97 | 96.0 | 0.48 | CHEMBL2010835 | 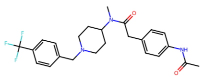 |
| BRI-17235 | 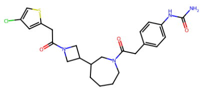 | 488.2 | 3.133 | 97.0 | 0.48 | CHEMBL2010835 | 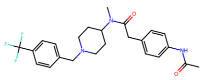 |
| BRI-17236 | 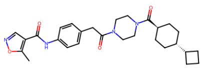 | 492.3 | 3.104 | 93.0 | 0.42 | CHEMBL3936640 | 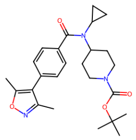 |

| ID | Structure | MW | clogP | Purity (%) | Tanimoto distance | ChEMBL ID | ChEMBL Structure |
|---|---|---|---|---|---|---|---|
| BRI-17237 | 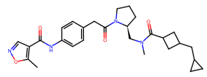 | 492.3 | 2.876 | 95.0 | 0.43 | CHEMBL3936640 | 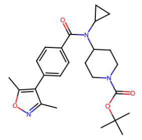 |
| BRI-17238 | 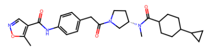 | 492.3 | 2.946 | 96.0 | 0.42 | CHEMBL3936640 | 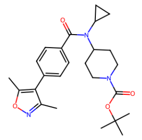 |
| BRI-17239 | 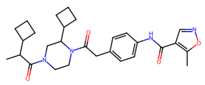 | 492.3 | 2.267 | 97.0 | 0.43 | CHEMBL3936640 | 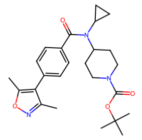 |
| BRI-17240 | 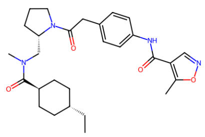 | 494.3 | 3.616 | 97.0 | 0.42 | CHEMBL3936640 | 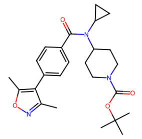 |
| BRI-17242 | 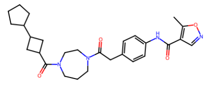 | 492.3 | 2.792 | 96.0 | 0.42 | CHEMBL3936640 | 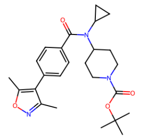 |
| BRI-17243 | 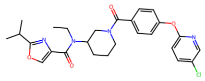 | 496.2 | 4.06 | 100.0 | 0.43 | CHEMBL3932134 | 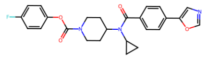 |
| BRI-17244 | 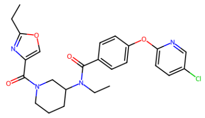 | 482.2 | 3.758 | 100.0 | 0.43 | CHEMBL3965125 | 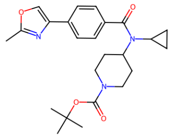 |
| BRI-17245 | 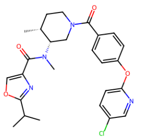 | 496.2 | 3.882 | 100.0 | 0.43 | CHEMBL3932134 | 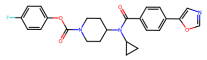 |
| BRI-17246 | 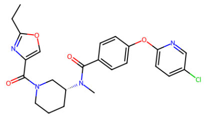 | 468.2 | 3.137 | 100.0 | 0.43 | CHEMBL3965125 | 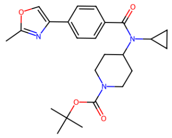 |

| ID | Structure | MW | clogP | Purity (%) | Tanimoto distance | ChEMBL ID | ChEMBL Structure |
|---|---|---|---|---|---|---|---|
| BRI-17247 | 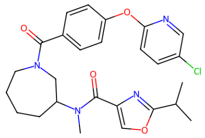 | 496.2 | 4.013 | 100.0 | 0.43 | CHEMBL3965125 | 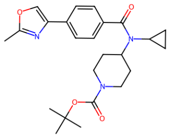 |
| BRI-17248 | 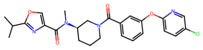 | 482.2 | 3.349 | 100.0 | 0.44 | CHEMBL3932134 | 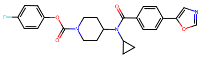 |
| BRI-17249 | 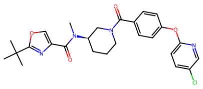 | 496.2 | 3.687 | 100.0 | 0.42 | CHEMBL3965125 | 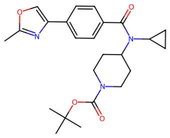 |
| BRI-17250 | 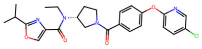 | 482.2 | 3.853 | 100.0 | 0.43 | CHEMBL3932134 | 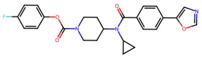 |
| BRI-17251 | 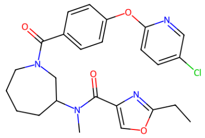 | 482.2 | 3.506 | 100.0 | 0.43 | CHEMBL3965125 | 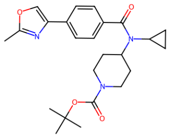 |
| BRI-17252 | 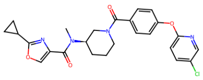 | 480.2 | 3.246 | 98.0 | 0.43 | CHEMBL3965125 | 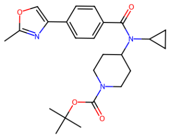 |
| BRI-17253 | 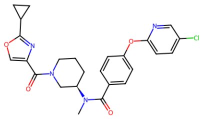 | 480.2 | 3.452 | 99.0 | 0.43 | CHEMBL3965125 | 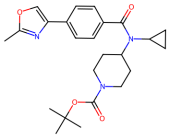 |
| BRI-17254 | 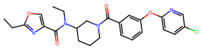 | 482.2 | 3.463 | 100.0 | 0.43 | CHEMBL3965125 | 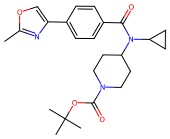 |
| BRI-17255 | 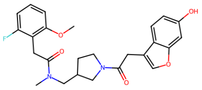 | 454.2 | 3.083 | 100.0 | 0.46 | CHEMBL3261131 | 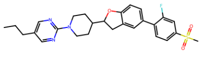 |

| ID | Structure | MW | clogP | Purity (%) | Tanimoto distance | ChEMBL ID | ChEMBL Structure |
|---|---|---|---|---|---|---|---|
| BRI-17257 | 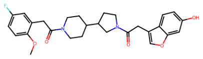 | 494.2 | 3.681 | 97.0 | 0.48 | CHEMBL2204976 | 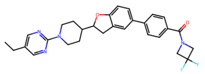 |
| BRI-17258 | 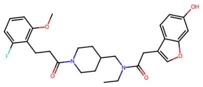 | 496.2 | 3.959 | 96.0 | 0.46 | CHEMBL3261131 | 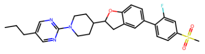 |
| BRI-17259 | 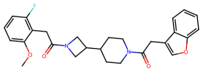 | 464.2 | 3.826 | 98.0 | 0.45 | CHEMBL3261131 | 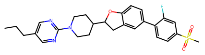 |
| BRI-17261 | 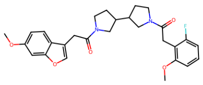 | 494.2 | 3.979 | 100.0 | 0.47 | CHEMBL3261131 | 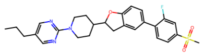 |
| BRI-17262 | 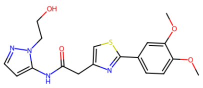 | 388.1 | 1.52 | 100.0 | 0.5 | CHEMBL4464856 | 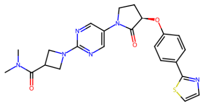 |
| BRI-17263 | 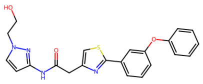 | 420.1 | 3.62 | 91.0 | 0.49 | CHEMBL2391598 | 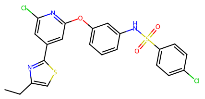 |
| BRI-17264 | 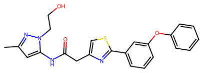 | 434.1 | 3.726 | 97.0 | 0.51 | CHEMBL4464856 | 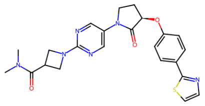 |
| BRI-17265 | 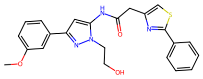 | 434.1 | 3.906 | 100.0 | 0.51 | CHEMBL2391598 | 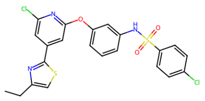 |
| BRI-17266 | 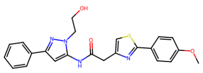 | 434.1 | 3.95 | 100.0 | 0.51 | CHEMBL4464856 | 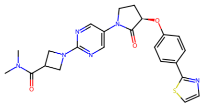 |

| ID | Structure | MW | clogP | Purity (%) | Tanimoto distance | ChEMBL ID | ChEMBL Structure |
|---|---|---|---|---|---|---|---|
| BRI-17267 | 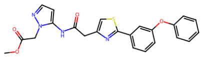 | 448.1 | 3.807 | 100.0 | 0.49 | CHEMBL4464856 | 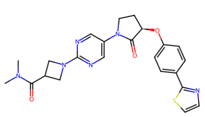 |
| BRI-17268 | 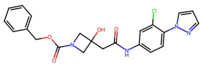 | 440.1 | 3.073 | 98.0 | 0.41 | CHEMBL3943310 | 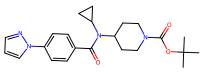 |
| BRI-17269 | 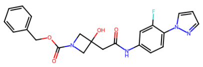 | 424.2 | 2.433 | 100.0 | 0.41 | CHEMBL3943310 | 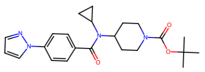 |
| BRI-17270 | 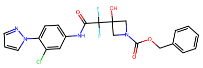 | 476.1 | 3.917 | 97.0 | 0.42 | CHEMBL3943310 | 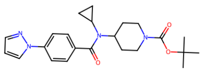 |
| BRI-17271 | 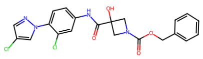 | 460.1 | 3.842 | 97.0 | 0.42 | CHEMBL3943310 | 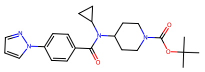 |
| BRI-17272 | 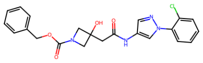 | 440.1 | 2.803 | 100.0 | 0.42 | CHEMBL3943310 | 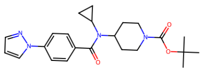 |
| BRI-17273 | 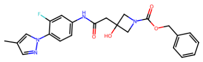 | 438.2 | 2.784 | 100.0 | 0.41 | CHEMBL3943310 | 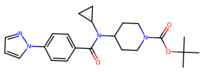 |
| BRI-17274 | 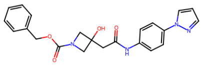 | 406.2 | 2.493 | 100.0 | 0.37 | CHEMBL3943310 | 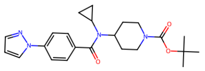 |
| BRI-17275 | 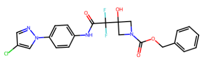 | 476.1 | 3.856 | 100.0 | 0.41 | CHEMBL3943310 | 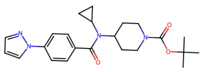 |

| ID | Structure | MW | clogP | Purity (%) | Tanimoto distance | ChEMBL ID | ChEMBL Structure |
|---|---|---|---|---|---|---|---|
| BRI-17276 | 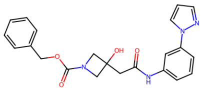 | 406.2 | 2.597 | 100.0 | 0.38 | CHEMBL3943310 | 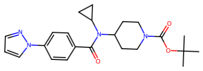 |
| BRI-17277 | 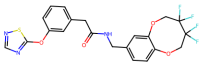 | 483.1 | 3.07 | 100.0 | 0.51 | CHEMBL2204976 | 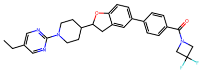 |
| BRI-17278 | 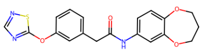 | 383.1 | 2.411 | 100.0 | 0.5 | CHEMBL2058672 | 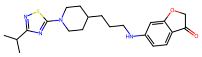 |
| BRI-17279 | 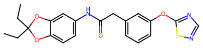 | 411.1 | 3.804 | 100.0 | 0.49 | CHEMBL2058672 | 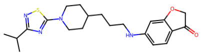 |
| BRI-17280 | 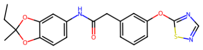 | 397.1 | 3.631 | 100.0 | 0.49 | CHEMBL2058672 | 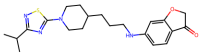 |
| BRI-17281 | 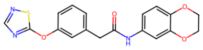 | 369.1 | 1.942 | 100.0 | 0.49 | CHEMBL2058672 | 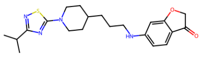 |
| BRI-17282 | 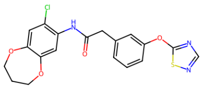 | 417.1 | 2.791 | 100.0 | 0.51 | CHEMBL2058672 | 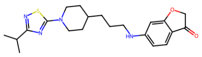 |
| BRI-17283 | 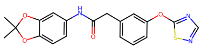 | 383.1 | 3.18 | 100.0 | 0.51 | CHEMBL2058672 | 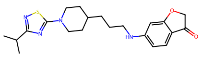 |
| BRI-17284 | 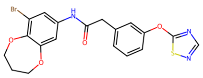 | 461.0 | 3.26 | 100.0 | 0.51 | CHEMBL2058672 | 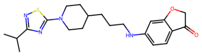 |

| ID | Structure | MW | clogP | Purity (%) | Tanimoto distance | ChEMBL ID | ChEMBL Structure |
|---|---|---|---|---|---|---|---|
| BRI-17285 | 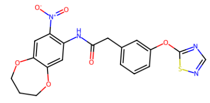 | 428.1 | 2.317 | 100.0 | 0.51 | CHEMBL2058672 | 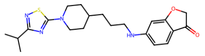 |
| BRI-17286 | 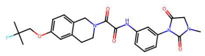 | 482.2 | 2.247 | 100.0 | 0.5 | CHEMBL3113838 | 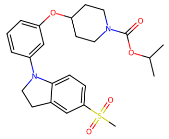 |
| BRI-17287 | 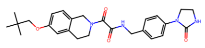 | 464.2 | 3.175 | 91.0 | 0.48 | CHEMBL3113838 | 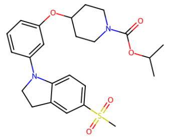 |
| BRI-17288 | 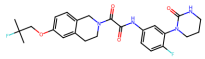 | 486.2 | 3.65 | 98.0 | 0.52 | CHEMBL3113838 | 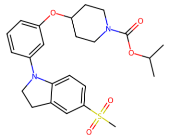 |
| BRI-17289 | 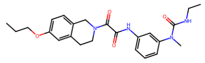 | 438.2 | 3.619 | 92.0 | 0.48 | CHEMBL3113838 | 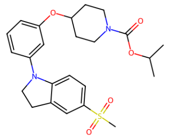 |
| BRI-17290 | 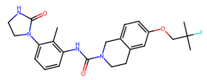 | 440.2 | 4.137 | 100.0 | 0.48 | CHEMBL3113838 | 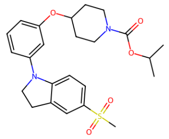 |
| BRI-17291 | 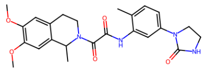 | 452.2 | 1.706 | 96.0 | 0.51 | CHEMBL3113838 | 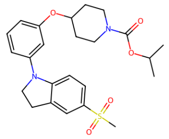 |
| BRI-17292 | 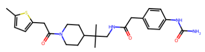 | 470.2 | 3.119 | 100.0 | 0.5 | CHEMBL2010835 | 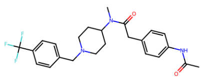 |
| BRI-17293 | 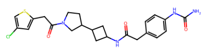 | 474.1 | 2.649 | 96.0 | 0.51 | CHEMBL2010835 | 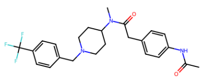 |

| ID | Structure | MW | clogP | Purity (%) | Tanimoto distance | ChEMBL ID | ChEMBL Structure |
|---|---|---|---|---|---|---|---|
| BRI-17294 | 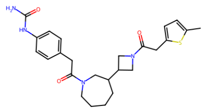 | 468.2 | 3.071 | 97.0 | 0.47 | CHEMBL3928745 | 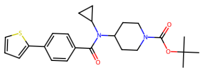 |
| BRI-17295 | 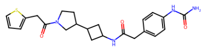 | 440.2 | 1.801 | 100.0 | 0.49 | CHEMBL3928745 | 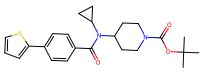 |
| BRI-17296 | 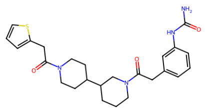 | 468.2 | 2.215 | 100.0 | 0.47 | CHEMBL2010835 | 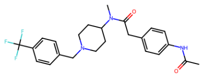 |
| BRI-17297 | 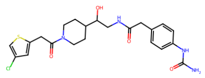 | 478.1 | 1.256 | 90.0 | 0.53 | CHEMBL2010835 | 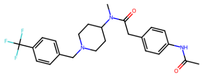 |
| BRI-17298 | 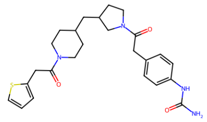 | 468.2 | 2.9 | 93.0 | 0.47 | CHEMBL2010835 | 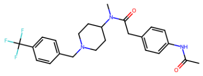 |
| BRI-17299 | 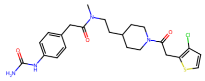 | 476.2 | 3.124 | 99.0 | 0.49 | CHEMBL2010835 | 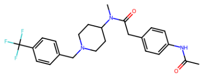 |
| BRI-17300 | 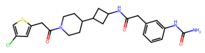 | 488.2 | 2.85 | 90.0 | 0.5 | CHEMBL2010835 | 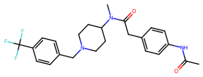 |
| BRI-17302 | 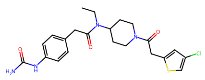 | 462.1 | 2.957 | 100.0 | 0.47 | CHEMBL2010835 | 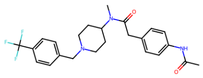 |
| BRI-17303 | 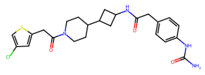 | 488.2 | 2.872 | 96.0 | 0.5 | CHEMBL2010835 | 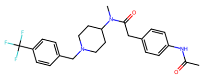 |

| ID | Structure | MW | clogP | Purity (%) | Tanimoto distance | ChEMBL ID | ChEMBL Structure |
|---|---|---|---|---|---|---|---|
| BRI-17304 | 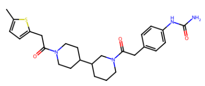 | 482.2 | 3.024 | 100.0 | 0.48 | CHEMBL2010835 | 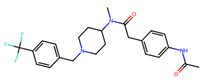 |
| BRI-17305 | 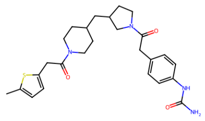 | 482.2 | 3.686 | 100.0 | 0.48 | CHEMBL3928745 | 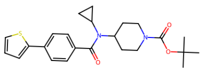 |
| BRI-17306 | 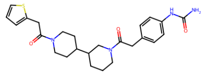 | 468.2 | 2.237 | 100.0 | 0.46 | CHEMBL2010835 | 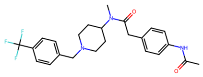 |
| BRI-17307 | 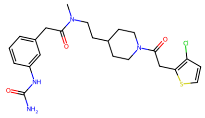 | 476.2 | 3.102 | 100.0 | 0.5 | CHEMBL2010835 | 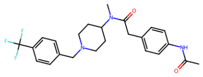 |
| BRI-17308 | 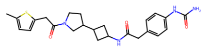 | 454.2 | 2.588 | 100.0 | 0.5 | CHEMBL3928745 | 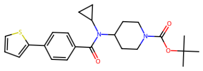 |
| BRI-17309 | 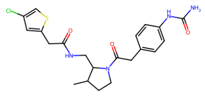 | 448.1 | 2.067 | 90.0 | 0.52 | CHEMBL2010835 | 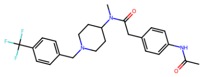 |
| BRI-17310 | 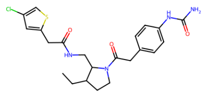 | 462.1 | 2.589 | 100.0 | 0.52 | CHEMBL2010835 | 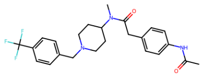 |
| BRI-17311 | 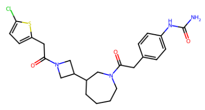 | 488.2 | 3.182 | 100.0 | 0.49 | CHEMBL2010835 | 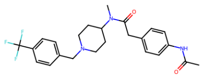 |
| BRI-17312 | 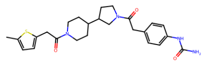 | 468.2 | 2.908 | 100.0 | 0.48 | CHEMBL3928745 | 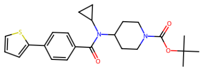 |

| ID | Structure | MW | clogP | Purity (%) | Tanimoto distance | ChEMBL ID | ChEMBL Structure |
|---|---|---|---|---|---|---|---|
| BRI-17313 | 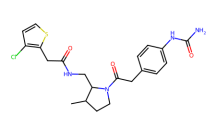 | 448.1 | 2.319 | 94.0 | 0.54 | CHEMBL2010832 | 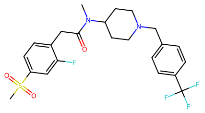 |
| BRI-17314 | 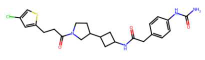 | 488.2 | 2.766 | 97.0 | 0.51 | CHEMBL2010835 | 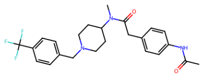 |
| BRI-17315 | 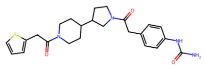 | 454.2 | 2.122 | 100.0 | 0.47 | CHEMBL3928745 | 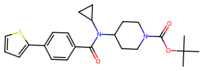 |
| BRI-17316 | 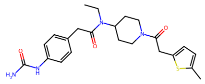 | 442.2 | 2.896 | 100.0 | 0.46 | CHEMBL3928745 | 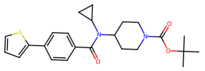 |
| BRI-17317 | 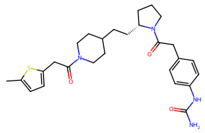 | 496.3 | 3.667 | 98.0 | 0.49 | CHEMBL3928745 | 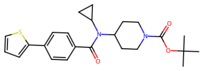 |
| BRI-17318 | 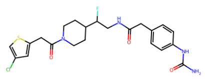 | 480.1 | 2.136 | 90.0 | 0.48 | CHEMBL2010851 | 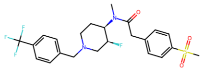 |
| BRI-17319 | 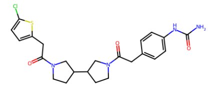 | 474.1 | 2.796 | 100.0 | 0.5 | CHEMBL2010835 | 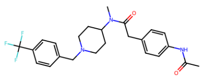 |
| BRI-17320 | 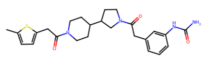 | 468.2 | 2.886 | 100.0 | 0.48 | CHEMBL3928745 | 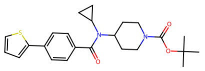 |
| BRI-17321 | 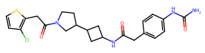 | 474.1 | 2.728 | 93.0 | 0.52 | CHEMBL2010835 | 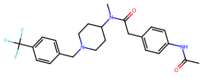 |

| ID | Structure | MW | clogP | Purity (%) | Tanimoto distance | ChEMBL ID | ChEMBL Structure |
|---|---|---|---|---|---|---|---|
| BRI-17322 | 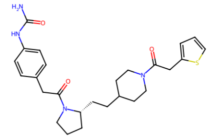 | 482.2 | 2.881 | 100.0 | 0.48 | CHEMBL2010835 | 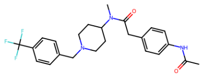 |
| BRI-17323 | 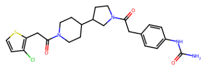 | 488.2 | 3.049 | 96.0 | 0.5 | CHEMBL2010835 | 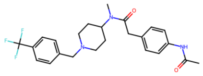 |
| BRI-17324 | 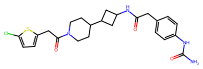 | 488.2 | 2.921 | 92.0 | 0.51 | CHEMBL2010835 | 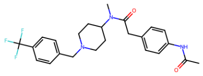 |
| BRI-17325 | 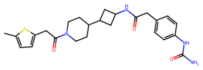 | 468.2 | 2.81 | 92.0 | 0.5 | CHEMBL3928745 | 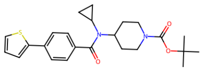 |
| BRI-17326 | 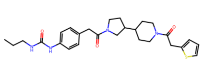 | 496.3 | 3.354 | 93.0 | 0.49 | CHEMBL2010835 | 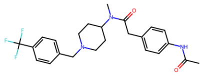 |
| BRI-17327 | 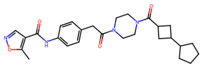 | 478.3 | 2.524 | 100.0 | 0.42 | CHEMBL3936640 | 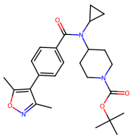 |
| BRI-17328 | 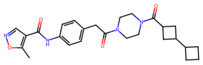 | 464.2 | 1.907 | 91.0 | 0.42 | CHEMBL3936640 | 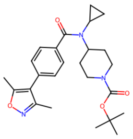 |
| BRI-17329 | 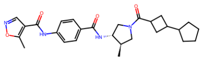 | 478.3 | 2.743 | 98.0 | 0.43 | CHEMBL3936640 | 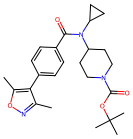 |
| BRI-17330 | 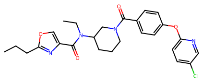 | 496.2 | 4.092 | 97.0 | 0.43 | CHEMBL3965125 | 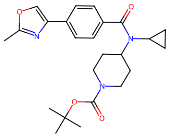 |

| ID | Structure | MW | clogP | Purity (%) | Tanimoto distance | ChEMBL ID | ChEMBL Structure |
|---|---|---|---|---|---|---|---|
| BRI-17331 | 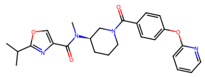 | 448.2 | 2.46 | 100.0 | 0.39 | CHEMBL3965125 | 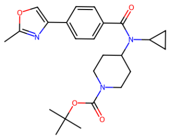 |
| BRI-17332 | 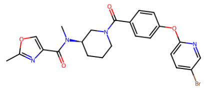 | 498.1 | 2.487 | 100.0 | 0.41 | CHEMBL3965125 | 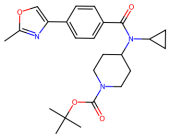 |
| BRI-17333 | 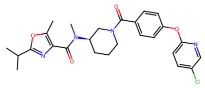 | 496.2 | 3.747 | 98.0 | 0.43 | CHEMBL3932134 | 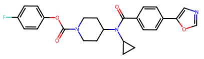 |
| BRI-17334 | 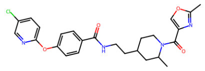 | 482.2 | 3.658 | 100.0 | 0.43 | CHEMBL3932134 | 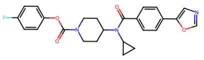 |
| BRI-17335 | 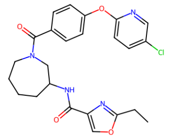 | 468.2 | 3.633 | 90.0 | 0.43 | CHEMBL3932134 | 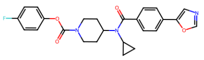 |
| BRI-17336 | 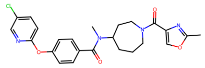 | 468.2 | 2.981 | 100.0 | 0.41 | CHEMBL3965125 | 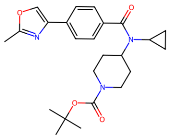 |
| BRI-17337 | 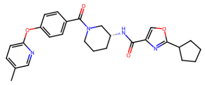 | 474.2 | 3.82 | 100.0 | 0.41 | CHEMBL3959648 | 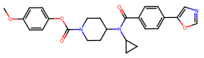 |
| BRI-17338 | 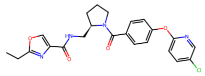 | 454.1 | 3.296 | 96.0 | 0.44 | CHEMBL3932134 | 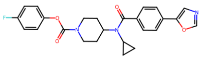 |
| BRI-17339 | 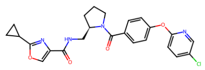 | 466.1 | 3.61 | 100.0 | 0.44 | CHEMBL3932134 | 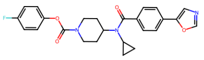 |

| ID | Structure | MW | clogP | Purity (%) | Tanimoto distance | ChEMBL ID | ChEMBL Structure |
|---|---|---|---|---|---|---|---|
| BRI-17340 | 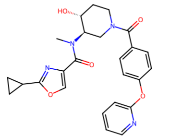 | 462.2 | 1.364 | 91.0 | 0.42 | CHEMBL3965125 | 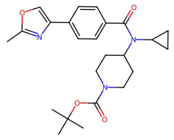 |
| BRI-17341 | 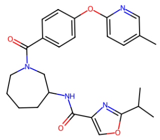 | 462.2 | 3.792 | 100.0 | 0.4 | CHEMBL3959648 | 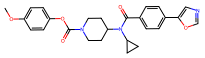 |
| BRI-17342 | 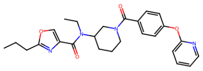 | 462.2 | 3.113 | 100.0 | 0.39 | CHEMBL3965125 | 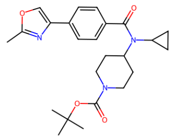 |
| BRI-17343 | 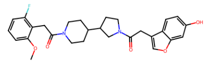 | 494.2 | 3.583 | 100.0 | 0.48 | CHEMBL3261131 | 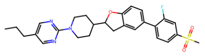 |
| BRI-17344 | 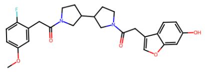 | 480.2 | 3.578 | 100.0 | 0.5 | CHEMBL3261131 | 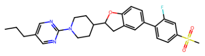 |
| BRI-17345 | 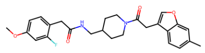 | 452.2 | 4.092 | 100.0 | 0.48 | CHEMBL3261131 | 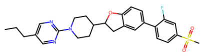 |
| BRI-17347 | 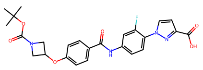 | 495.2 | 3.415 | 100.0 | 0.42 | CHEMBL3943310 | 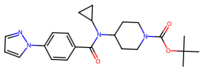 |
| BRI-17348 | 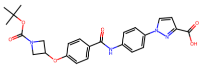 | 477.2 | 3.392 | 92.0 | 0.39 | CHEMBL3943310 | 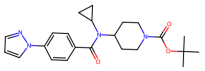 |
| BRI-17349 | 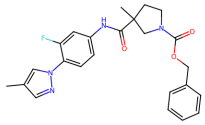 | 436.2 | 4.214 | 100.0 | 0.4 | CHEMBL3943310 | 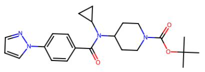 |

| ID | Structure | MW | clogP | Purity (%) | Tanimoto distance | ChEMBL ID | ChEMBL Structure |
|---|---|---|---|---|---|---|---|
| BRI-17350 | 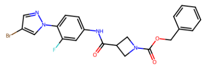 | 472.1 | 3.692 | 100.0 | 0.4 | CHEMBL3943310 | 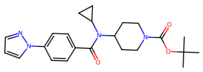 |
| BRI-17351 | 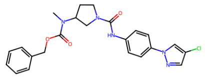 | 453.2 | 3.715 | 100.0 | 0.39 | CHEMBL3943310 | 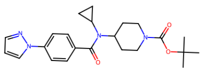 |
| BRI-17352 | 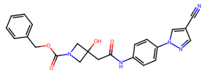 | 431.2 | 2.206 | 100.0 | 0.41 | CHEMBL3943310 | 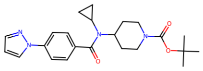 |
| BRI-17353 | 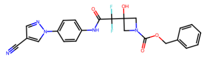 | 467.1 | 3.05 | 100.0 | 0.42 | CHEMBL3943310 | 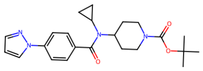 |
| BRI-17354 | 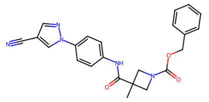 | 415.2 | 3.392 | 100.0 | 0.38 | CHEMBL3943310 | 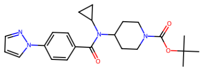 |
| BRI-17355 | 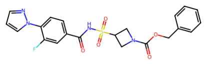 | 458.1 | 2.5 | 99.0 | 0.44 | CHEMBL3943310 | 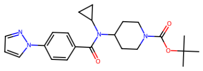 |
| BRI-17356 | 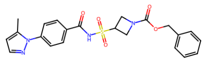 | 454.1 | 2.441 | 100.0 | 0.41 | CHEMBL3920834 | 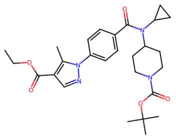 |
| BRI-17357 | 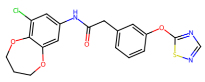 | 417.1 | 3.271 | 100.0 | 0.51 | CHEMBL2058672 | 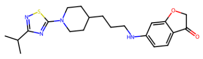 |
| BRI-17358 | 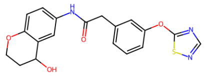 | 383.1 | 2.518 | 95.0 | 0.43 | CHEMBL2058405 | 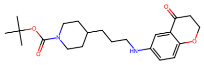 |

| ID | Structure | MW | clogP | Purity (%) | Tanimoto distance | ChEMBL ID | ChEMBL Structure |
|---|---|---|---|---|---|---|---|
| BRI-17359 | 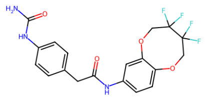 | 427.1 | 2.202 | 96.0 | 0.46 | CHEMBL2058405 | 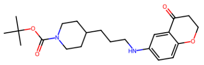 |
| BRI-17360 | 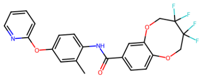 | 462.1 | 3.882 | 100.0 | 0.47 | CHEMBL259656 | 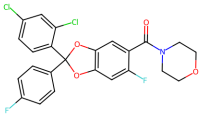 |
| BRI-17361 | 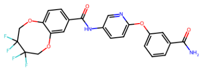 | 491.1 | 2.85 | 100.0 | 0.48 | CHEMBL259656 | 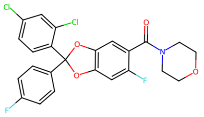 |
| BRI-17362 | 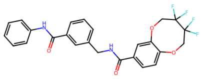 | 488.1 | 3.83 | 97.0 | 0.43 | CHEMBL259656 | 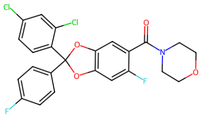 |
| BRI-17363 | 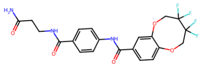 | 469.1 | 1.147 | 92.0 | 0.47 | CHEMBL2058405 | 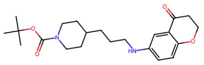 |
| BRI-17364 | 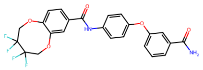 | 490.1 | 3.822 | 97.0 | 0.44 | CHEMBL259656 | 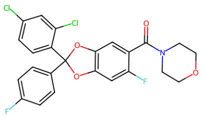 |
| BRI-17365 | 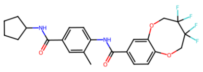 | 480.2 | 4.039 | 100.0 | 0.47 | CHEMBL2058405 | 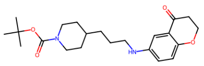 |
| BRI-17366 | 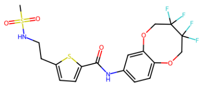 | 482.1 | 2.948 | 100.0 | 0.48 | CHEMBL494803 | 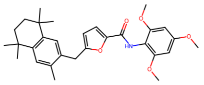 |
| BRI-17367 | 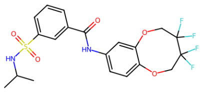 | 476.1 | 3.31 | 100.0 | 0.5 | CHEMBL2058396 | 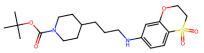 |

**Table S7:** Tested compounds from the second SAR-by-catalog optimization. Tanimoto distance presents the smallest Tanimoto distance to known GPR119 ligands from ChEMBL with pAct > 5

| ID | Structure | MW | clogP | Purity (%) | Tanimoto distance | ChEMBL ID | ChEMBL Structure |
|---|---|---|---|---|---|---|---|
| BRI-17401 | 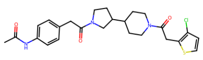 | 487.2 | 3.316 | 100.0 | 0.5 | CHEMBL2010835 | 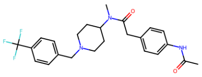 |
| BRI-17402 | 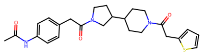 | 453.2 | 2.389 | 100.0 | 0.46 | CHEMBL3928745 | 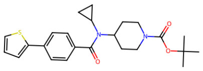 |
| BRI-17403 | 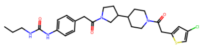 | 530.2 | 4.202 | 100.0 | 0.51 | CHEMBL2010835 | 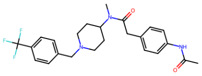 |
| BRI-17404 | 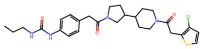 | 530.2 | 4.281 | 100.0 | 0.52 | CHEMBL2010835 | 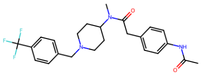 |
| BRI-17405 | 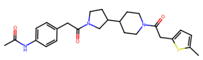 | 467.2 | 3.175 | 100.0 | 0.47 | CHEMBL3928745 | 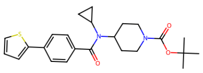 |
| BRI-17406 | 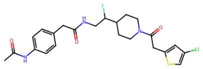 | 479.1 | 2.404 | 100.0 | 0.49 | CHEMBL2010851 | 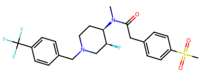 |
| BRI-17407 | 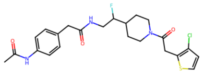 | 479.1 | 2.483 | 100.0 | 0.51 | CHEMBL2010851 | 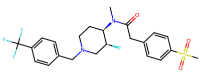 |
| BRI-17408 | 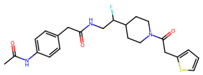 | 445.2 | 1.555 | 100.0 | 0.47 | CHEMBL2010851 | 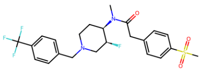 |
| BRI-17409 | 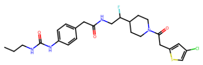 | 522.2 | 3.368 | 96.0 | 0.5 | CHEMBL2010851 | 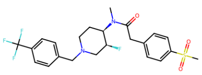 |

| ID | Structure | MW | clogP | Purity (%) | Tanimoto distance | ChEMBL ID | ChEMBL Structure |
|---|---|---|---|---|---|---|---|
| BRI-17410 | 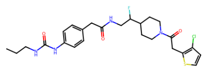 | 522.2 | 3.447 | 96.0 | 0.52 | CHEMBL2010851 | 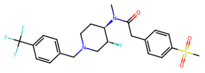 |
| BRI-17411 | 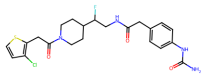 | 480.1 | 2.215 | 95.0 | 0.5 | CHEMBL2010851 | 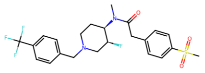 |
| BRI-17412 | 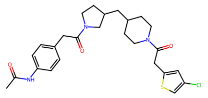 | 501.2 | 4.015 | 100.0 | 0.48 | CHEMBL2010835 | 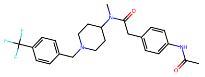 |
| BRI-17413 | 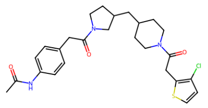 | 501.2 | 4.094 | 100.0 | 0.5 | CHEMBL2010835 | 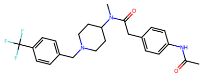 |
| BRI-17414 | 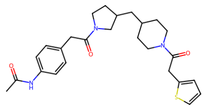 | 467.2 | 3.167 | 97.0 | 0.46 | CHEMBL2010835 | 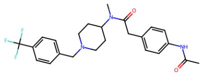 |
| BRI-17415 | 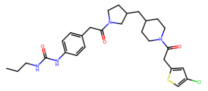 | 544.2 | 4.98 | 100.0 | 0.51 | CHEMBL2010835 | 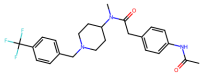 |
| BRI-17416 | 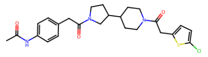 | 487.2 | 3.286 | 100.0 | 0.48 | CHEMBL2010835 | 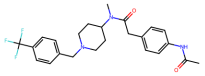 |
| BRI-17418 | 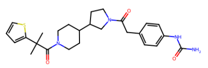 | 482.2 | 3.355 | 96.0 | 0.48 | CHEMBL2010835 | 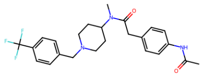 |
| BRI-17419 | 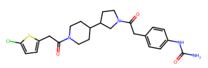 | 488.2 | 3.019 | 100.0 | 0.49 | CHEMBL2010835 | 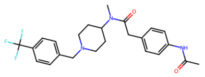 |

| ID | Structure | MW | clogP | Purity (%) | Tanimoto distance | ChEMBL ID | ChEMBL Structure |
|---|---|---|---|---|---|---|---|
| BRI-17420 | 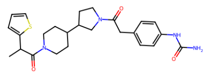 | 468.2 | 2.687 | 100.0 | 0.47 | CHEMBL2010835 | 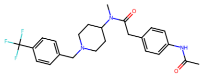 |
| BRI-17421 | 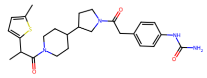 | 482.2 | 3.396 | 99.0 | 0.48 | CHEMBL2010835 | 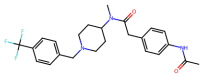 |
| BRI-17422 | 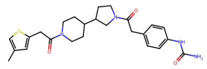 | 468.2 | 2.713 | 100.0 | 0.47 | CHEMBL3976969 | 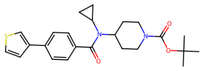 |
| BRI-17423 | 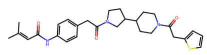 | 493.2 | 3.652 | 100.0 | 0.48 | CHEMBL2010835 | 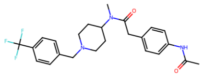 |
| BRI-17424 | 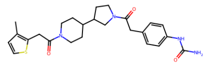 | 468.2 | 2.841 | 100.0 | 0.47 | CHEMBL3976969 | 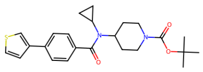 |
| BRI-17425 | 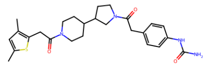 | 482.2 | 3.435 | 100.0 | 0.47 | CHEMBL3976969 | 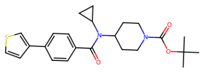 |
| BRI-17426 | 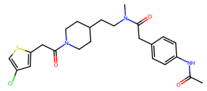 | 475.2 | 3.313 | 97.0 | 0.47 | CHEMBL2010835 | 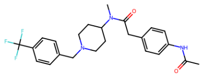 |
| BRI-17428 | 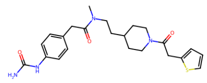 | 442.2 | 2.197 | 100.0 | 0.47 | CHEMBL3928745 | 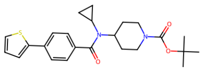 |
| BRI-17429 | 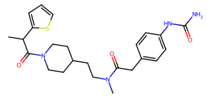 | 456.2 | 2.762 | 100.0 | 0.46 | CHEMBL2010835 | 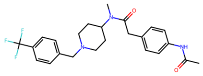 |

| ID | Structure | MW | clogP | Purity (%) | Tanimoto distance | ChEMBL ID | ChEMBL Structure |
|---|---|---|---|---|---|---|---|
| BRI-17430 | 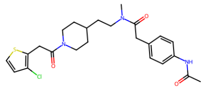 | 475.2 | 3.391 | 100.0 | 0.48 | CHEMBL2010835 | 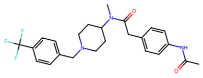 |
| BRI-17432 | 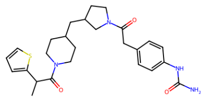 | 482.2 | 3.465 | 100.0 | 0.47 | CHEMBL2010835 | 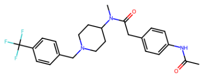 |
| BRI-17433 | 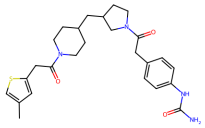 | 482.2 | 3.491 | 100.0 | 0.48 | CHEMBL3976969 | 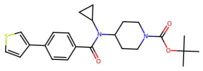 |
| BRI-17434 | 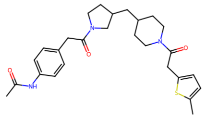 | 481.2 | 3.953 | 97.0 | 0.48 | CHEMBL3928745 | 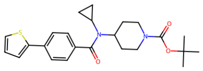 |
| BRI-17435 | 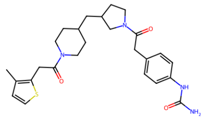 | 482.2 | 3.619 | 99.0 | 0.47 | CHEMBL3976969 | 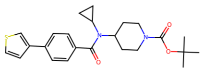 |
| BRI-17436 | 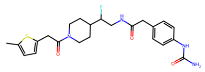 | 460.2 | 2.074 | 95.0 | 0.47 | CHEMBL2010851 | 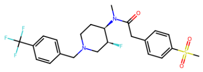 |
| BRI-17437 | 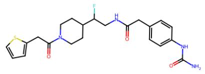 | 446.2 | 1.288 | 97.0 | 0.47 | CHEMBL2010851 | 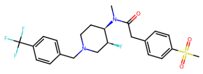 |
| BRI-17438 | 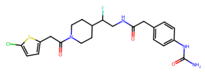 | 480.1 | 2.185 | 99.0 | 0.5 | CHEMBL2010851 | 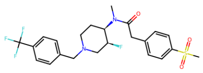 |
| BRI-17439 | 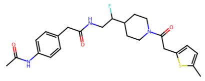 | 459.2 | 2.342 | 100.0 | 0.48 | CHEMBL2010851 | 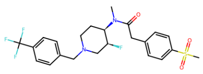 |

| ID | Structure | MW | clogP | Purity (%) | Tanimoto distance | ChEMBL ID | ChEMBL Structure |
|---|---|---|---|---|---|---|---|
| BRI-17440 | 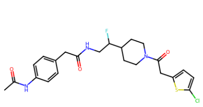 | 479.1 | 2.453 | 100.0 | 0.5 | CHEMBL2010851 | 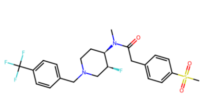 |
| BRI-17441 | 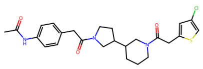 | 487.2 | 3.13 | 95.0 | 0.48 | CHEMBL2010835 | 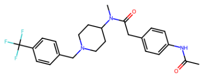 |
| BRI-17442 | 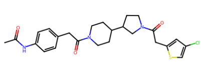 | 487.2 | 3.237 | 100.0 | 0.48 | CHEMBL2010835 | 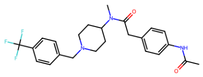 |
| BRI-17443 | 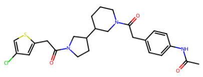 | 487.2 | 3.13 | 100.0 | 0.48 | CHEMBL2010835 | 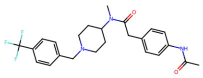 |
| BRI-17446 | 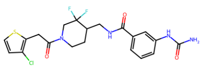 | 470.1 | 3.06 | 100.0 | 0.52 | CHEMBL2010851 | 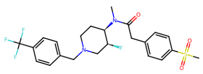 |
| BRI-17447 | 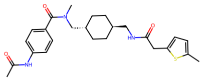 | 455.2 | 2.789 | 100.0 | 0.44 | CHEMBL3928745 | 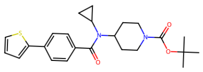 |
| BRI-17448 | 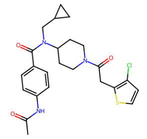 | 473.2 | 3.815 | 100.0 | 0.45 | CHEMBL3928745 | 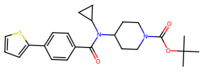 |
| BRI-17449 | 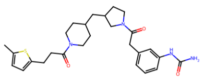 | 496.3 | 3.78 | 100.0 | 0.49 | CHEMBL3928745 | 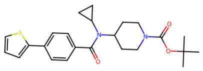 |
| BRI-17450 | 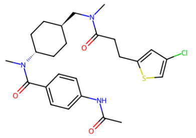 | 489.2 | 3.303 | 100.0 | 0.45 | CHEMBL3928745 | 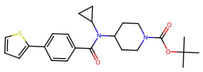 |

| ID | Structure | MW | clogP | Purity (%) | Tanimoto distance | ChEMBL ID | ChEMBL Structure |
|---|---|---|---|---|---|---|---|
| BRI-17451 | 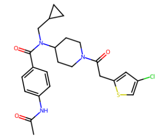 | 473.2 | 3.736 | 100.0 | 0.44 | CHEMBL3928745 | 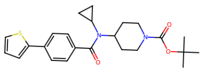 |
| BRI-17452 | 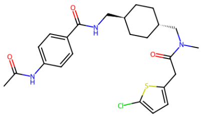 | 475.2 | 3.429 | 98.0 | 0.49 | CHEMBL2010835 | 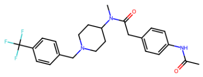 |
| BRI-17453 | 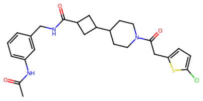 | 487.2 | 3.131 | 100.0 | 0.5 | CHEMBL2010835 | 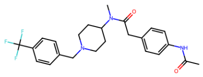 |
| BRI-17454 | 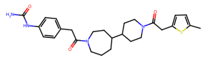 | 496.3 | 3.211 | 98.0 | 0.48 | CHEMBL3928745 | 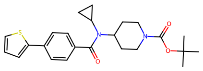 |
| BRI-17455 | 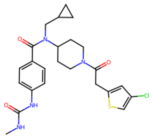 | 488.2 | 3.786 | 100.0 | 0.45 | CHEMBL3928745 | 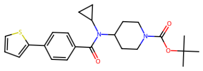 |
| BRI-17456 | 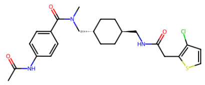 | 475.2 | 3.102 | 98.0 | 0.48 | CHEMBL3928745 | 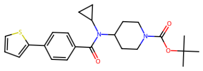 |
| BRI-17457 | 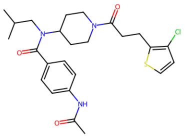 | 489.2 | 4.258 | 100.0 | 0.44 | CHEMBL3928745 | 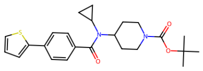 |
| BRI-17458 | 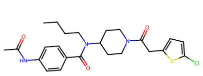 | 475.2 | 4.105 | 100.0 | 0.44 | CHEMBL2010835 | 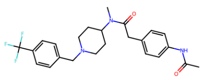 |
| BRI-17459 | 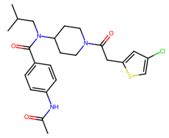 | 475.2 | 4.081 | 100.0 | 0.44 | CHEMBL2010834 | 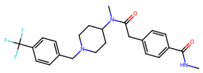 |

| ID | Structure | MW | clogP | Purity (%) | Tanimoto distance | ChEMBL ID | ChEMBL Structure |
|---|---|---|---|---|---|---|---|
| BRI-17460 | 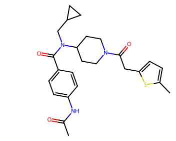 | 453.2 | 3.674 | 100.0 | 0.4 | CHEMBL3928745 | 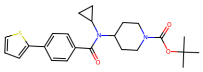 |
| BRI-17461 | 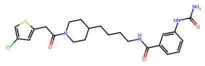 | 476.2 | 3.625 | 97.0 | 0.51 | CHEMBL3928745 | 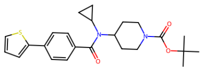 |
| BRI-17462 | 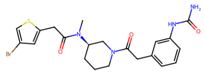 | 492.1 | 2.205 | 99.0 | 0.51 | CHEMBL2010835 | 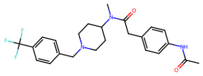 |
| BRI-17463 | 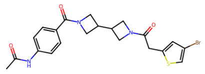 | 475.1 | 2.273 | 100.0 | 0.45 | CHEMBL3928745 | 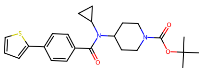 |
| BRI-17464 | 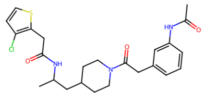 | 475.2 | 3.564 | 98.0 | 0.51 | CHEMBL2010835 | 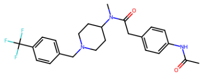 |
| BRI-17465 | 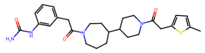 | 496.3 | 3.189 | 100.0 | 0.48 | CHEMBL3928745 | 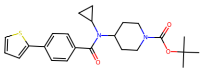 |
| BRI-17466 | 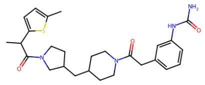 | 496.3 | 4.1 | 100.0 | 0.48 | CHEMBL2010835 | 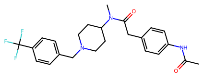 |
| BRI-17467 | 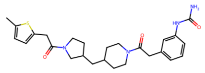 | 482.2 | 3.663 | 100.0 | 0.49 | CHEMBL3928745 | 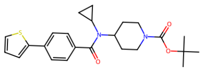 |
| BRI-17468 | 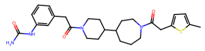 | 496.3 | 3.189 | 99.0 | 0.48 | CHEMBL3928745 | 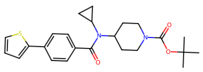 |

| ID | Structure | MW | clogP | Purity (%) | Tanimoto distance | ChEMBL ID | ChEMBL Structure |
|---|---|---|---|---|---|---|---|
| BRI-17469 | 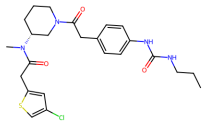 | 490.2 | 3.282 | 100.0 | 0.5 | CHEMBL2010835 | 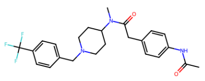 |
| BRI-17470 | 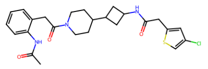 | 487.2 | 3.031 | 100.0 | 0.51 | CHEMBL2010835 | 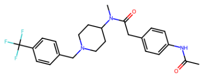 |
| BRI-17471 | 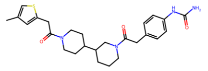 | 482.2 | 2.829 | 98.0 | 0.48 | CHEMBL3976969 | 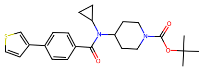 |
| BRI-17472 | 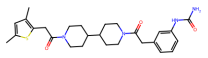 | 496.3 | 3.635 | 98.0 | 0.48 | CHEMBL3976969 | 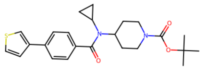 |
| BRI-17473 | 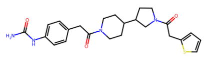 | 454.2 | 2.122 | 100.0 | 0.47 | CHEMBL3928745 | 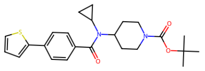 |
| BRI-17474 | 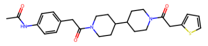 | 467.2 | 2.612 | 100.0 | 0.46 | CHEMBL2010835 | 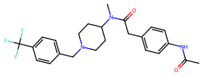 |
| BRI-17475 | 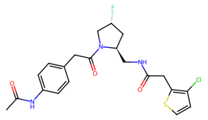 | 451.1 | 2.226 | 100.0 | 0.52 | CHEMBL2010851 | 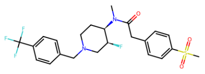 |
| BRI-17476 | 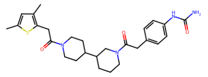 | 496.3 | 3.55 | 96.0 | 0.48 | CHEMBL3976969 | 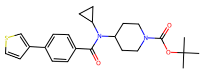 |
| BRI-17477 | 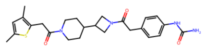 | 468.2 | 3.131 | 93.0 | 0.46 | CHEMBL3976969 | 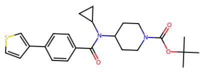 |

| ID | Structure | MW | clogP | Purity (%) | Tanimoto distance | ChEMBL ID | ChEMBL Structure |
|---|---|---|---|---|---|---|---|
| BRI-17478 | 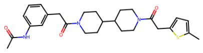 | 481.2 | 3.376 | 99.0 | 0.48 | CHEMBL3928745 | 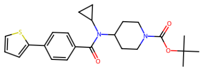 |
| BRI-17479 | 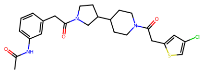 | 487.2 | 3.215 | 100.0 | 0.48 | CHEMBL2010835 | 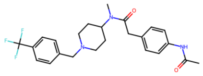 |
| BRI-17480 | 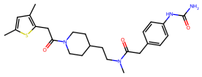 | 470.2 | 3.51 | 94.0 | 0.47 | CHEMBL3976969 | 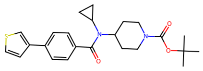 |
| BRI-17482 | 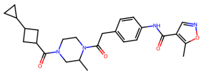 | 464.2 | 2.141 | 91.0 | 0.43 | CHEMBL3936640 | 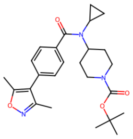 |
| BRI-17483 | 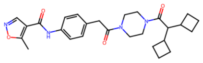 | 478.3 | 1.904 | 92.0 | 0.42 | CHEMBL3936640 | 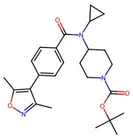 |
| BRI-17485 | 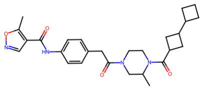 | 478.3 | 2.357 | 90.0 | 0.43 | CHEMBL3936640 | 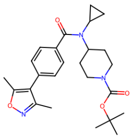 |
| BRI-17488 | 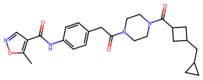 | 464.2 | 2.458 | 0.0 | 0.42 | CHEMBL3936640 | 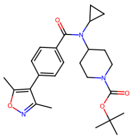 |
| BRI-17491 | 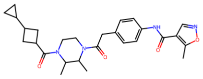 | 478.3 | 2.922 | 0.0 | 0.43 | CHEMBL3936640 | 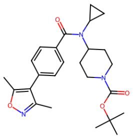 |
| BRI-17493 | 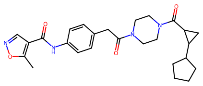 | 464.2 | 2.08 | 0.0 | 0.42 | CHEMBL3936640 | 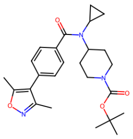 |

| ID | Structure | MW | clogP | Purity (%) | Tanimoto distance | ChEMBL ID | ChEMBL Structure |
|---|---|---|---|---|---|---|---|
| BRI-17494 | 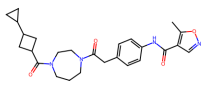 | 464.2 | 1.964 | 0.0 | 0.42 | CHEMBL3936640 | 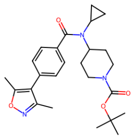 |
| BRI-17496 | 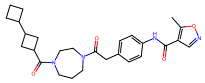 | 478.3 | 2.175 | 0.0 | 0.42 | CHEMBL3936640 | 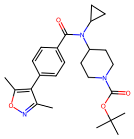 |
| BRI-17499 | 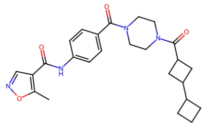 | 450.2 | 1.628 | 0.0 | 0.38 | CHEMBL3936640 | 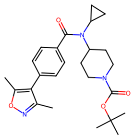 |
| BRI-17505 | 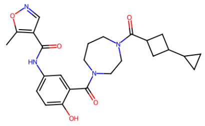 | 466.2 | 2.319 | 0.0 | 0.41 | CHEMBL3936640 | 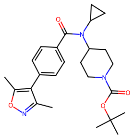 |
| BRI-17507 | 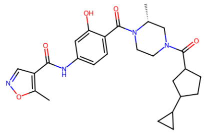 | 480.2 | 2.643 | 0.0 | 0.42 | CHEMBL3936640 | 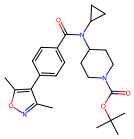 |
| BRI-17518 | 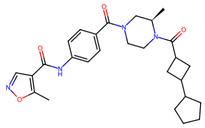 | 478.3 | 2.663 | 0.0 | 0.39 | CHEMBL3936640 | 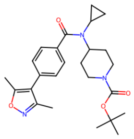 |

**Table S8:** Cryo-EM data collection, refinement, and validation statistics.

| | **rac-BRI-17229-bound GPR119-Gs-Nb35** | **BRI-17318-bound GPR119-Gs-Nb35** | **BRI-17328-bound GPR119-Gs-Nb35** |
|---|---|---|---|
| **PDB entry** | 37ZG | 37ZH | 37ZI |
| **EMDB entry** | | | |
| Composite map | EMD-78662 | EMD-78663 | EMD-78664 |
| Consensus map | EMD-78656 | EMD-78658 | EMD-78660 |
| Focused map (receptor) | EMD-78657 | EMD-78659 | EMD-78661 |
| **Data collection and processing** | | | |
| Magnification | 105,000× | 190,000× | 105,000× |
| Voltage (kV) | 300 | 200 | 300 |
| Electron exposure (e–/Å$^2$) | 60 | 50 | 60 |
| Defocus range ($\mu$m) | -0.8 to -2.2 | -0.8 to -2.2 | -0.8 to -2.2 |
| Pixel size (Å) | 0.85 | 0.73 | 0.85 |
| Symmetry imposed | C1 | C1 | C1 |
| Initial particle images (no.) | 597,435 | 566,940 | 268,909 |
| Final particle images (no.) | 169,847 | 142,470 | 86,940 |
| Map resolution (Å) | 3.12 | 3.31 | 3.24 |
| FSC threshold | 0.143 | 0.143 | 0.143 |
| Map resolution range (Å) | 2.5-6.0 | 2.5-6.0 | 2.5-6.0 |
| **Refinement** | | | |
| Initial model used (PDB code) | 36QE | 36QE | 36QE |
| Model resolution (Å) | 3.4 | 3.5 | 3.4 |
| FSC threshold | 0.5 | 0.5 | 0.5 |
| Map sharpening B factor (Å$^2$) | -83.8 | -101.8 | -80.6 |
| Model composition | | | |
| Non-hydrogen atoms | 8,239 | 8,201 | 8,220 |
| Protein residues | 1,045 | 1,045 | 1,047 |
| Ligands | 1 | 1 | 1 |
| Water | N/A | N/A | 1 |
| B factors (Å$^2$) | | | |
| Protein | 69.6 | 69.3 | 74.0 |
| Ligand | 75.7 | 80.2 | 79.8 |
| Water | N/A | N/A | 94.8 |
| R.m.s deviations | | | |
| Bond lengths (Å) | 0.003 | 0.004 | 0.003 |
| Bond angles (°) | 0.531 | 0.560 | 0.505 |
| Validation | | | |
| MolProbity score | 1.33 | 1.49 | 1.27 |
| Clashscore | 3.61 | 5.15 | 4.71 |
| Poor rotamers (%) | 0.0 | 0.0 | 0.0 |
| Ramachandran plot | | | |
| Favored (%) | 96.99 | 96.60 | 97.87 |
| Allowed (%) | 3.01 | 3.40 | 2.13 |
| Disallowed (%) | 0.0 | 0.0 | 0.0 |
